# Childhood brain tumours instruct cranial haematopoiesis and immunotolerance

**DOI:** 10.1101/2025.09.25.678472

**Authors:** Elizabeth Cooper, David A. Posner, Colin Y.C. Lee, Linda Hu, Sigourney Bonner, Jessica T. Taylor, Oscar Baldwin, Rocio Jimenez-Guerrero, Katherine E. Masih, Katherine Wickham Rahrmann, Jason Eigenbrood, Gina Ngo, Luca Porcu, Valar Nila Roamio Franklin, Clive S. D’Santos, Richard Mair, Thomas Santarius, Claudia Craven, Ibrahim Jalloh, Julia Moreno Vicente, Timotheus Y. F. Halim, Li Wang, Arnold R. Kreigstien, Fredrik J. Swartling, Javed Khan, Menna R. Clatworthy, Richard J. Gilbertson

**Author notes:** Correspondence to: EC and RJG.

## Abstract

Recent research has revealed a remarkable role for immunosurveillance in healthy and diseased brains, dispelling the notion that this organ is a passive immune-privileged site^1–3^. Better understanding of how this immunosurveillance operates could improve the treatment of neurological diseases. Here, using a novel genetically engineered mouse model of *ZFTA-RELA* ependymoma^4^–a childhood brain tumour–we characterised an immune circuit between the tumour and antigen presenting, haematopoietic stem/progenitor cells (HSPCs) in the skull bone marrow. The presentation of antigens in the cerebrospinal fluid (CSF) by HSPCs to CD4^+^ T cells, biased HSPC lineages toward myelopoiesis and polarised CD4^+^ T-cells to regulatory T cells (T- regs), culminating in tumour immunotolerance. Remarkably, a single infusion of antibodies directed against cytokines enriched in the CSF of mice bearing *ZFTA-RELA* ependymomas, choroid plexus carcinomas or Group-3 medulloblastoma–all aggressive childhood brain tumours–disrupted this process and caused profound tumour regression. These data unmask a mechanism by which skull bone marrow-derived HSPCs and CD4^+^ T cells cooperate to promote the immunotolerance of childhood brain tumours. Antibodies that disrupt this immunosurveillance could prove an effective therapy for these cancers that are less toxic than current treatments.

## MAIN TEXT

Almost all childhood brain tumours–the leading cause of childhood cancer death–are initiated in the embryonic brain^5^. These tumours retain the transcriptomic, morphological and functional characteristics of developing neural tissues, including the propagation of cell lineages from perivascular niches^6–8^. Therefore, since these tumours are initiated *in utero* before the immune system is fully mature^9^, it is possible that these cancers are tolerated as ‘self’. Understanding how the immune system interacts with childhood brain tumours may therefore improve the use of alternative treatments, including immunotherapies.

Immunotherapies have had limited success in the treatment of childhood brain tumours, but they could ultimately prove more effective and less toxic than conventional surgery, radiation and chemotherapy^10^. The failure of immunotherapies to elicit an immune response in childhood brain tumours suggests a local source of immunosuppression: a notion supported by evidence that immune circuits suppress autoreactive inflammatory responses in the brain and certain adult brain tumours^7,11–15^. Indeed, CSF flows directly to the skull bone marrow, feeding immune cells with brain-derived signals and driving immunosurveillance^11,16,17^; but the mechanisms by which these signals influence haematopoiesis remain unclear. Here, we provide evidence that this circuit operates in childhood brain tumours promoting the tolerance of these aggressive cancers, unmasking a new therapeutic vulnerability.

### HSPCs in *ZFTA-RELA* ependymoma

To understand how immunosurveillance might operate in childhood brain tumours, we developed a genetically engineered mouse model of supratentorial ependymoma (hereon, EP*^ZFTA-RELA^*) in which a conditional allele of the *ZFTA-RELA* fusion gene^4^ *(Nestin^Flx-STOP-FlxZFTA-RELA^*) is recombined in embryonic day (E)9.5 radial glia by the *Nestin^CreERT^*^2^ allele (**Supplementary Fig. 1a-e; Supplementary Table 1**): we originally identified radial glia as the cell of origin of ependymomas^4,6^. All *Nestin^CreERT2^*; *Nestin^Flx-STOP-FlxZFTA-RELA^* (hereon, *Nestin^Cre-ZFTA-RELA^*) mice developed EP*^ZFTA-RELA^* tumours with a median survival of 90 ± 9.5 days: these tumours contain neural progenitor-like cells highly enriched for a previously established human *ZFTA-RELA* ependymoma gene signature (**Supplementary Fig. 1f-h**)^4,6^.

Flow cytometric analysis of both *Nestin^Cre-ZFTA-RELA^* and *Nestin^CreERT2-WT^* (hereon, control) mouse brains at E12.5 identified similar populations of CD45^+^ cells, including lineage-negative/Sca-1^+^/c-Kit^+^ (LSK^+^) HSPCs that were previously described in normal embryonic mouse brain and human glioblastomas^21,22^ (**Fig. 1a,b; Supplementary Fig. 2; Supplementary Tables 2 and 3**). Following birth, CD45^+^ cell populations in the brains of *Nestin^Cre-ZFTA-RELA^* and control mice diverged markedly. By postnatal day (P)5, control mouse brains from which circulating CD45^+^ cells were excluded during flow cytometry by CD45 intravenous labelling^23^, lacked HSPCs and certain other immune cell types. In stark contrast, most immune cell populations, including HSPCs and Tregs, persisted and/or expanded in *Nestin^Cre-ZFTA-RELA^* brains. This began before appreciable tumour development (histologically undetectable before P19; **Supplementary Fig. 1c**) and persisted as tumours formed (**Fig. 1a-d**). Single cell RNA sequencing (scRNA-seq) of normal human foetal brain^24,25^ and 14 different types of human childhood brain tumour^26–30^, including ZFTA-RELA ependymoma and Group-3 medulloblastoma, confirmed the presence of similar immune populations in these tissues **(Fig. 1e,f; Supplementary Fig. 3a-f)**.

**Fig. 1:**
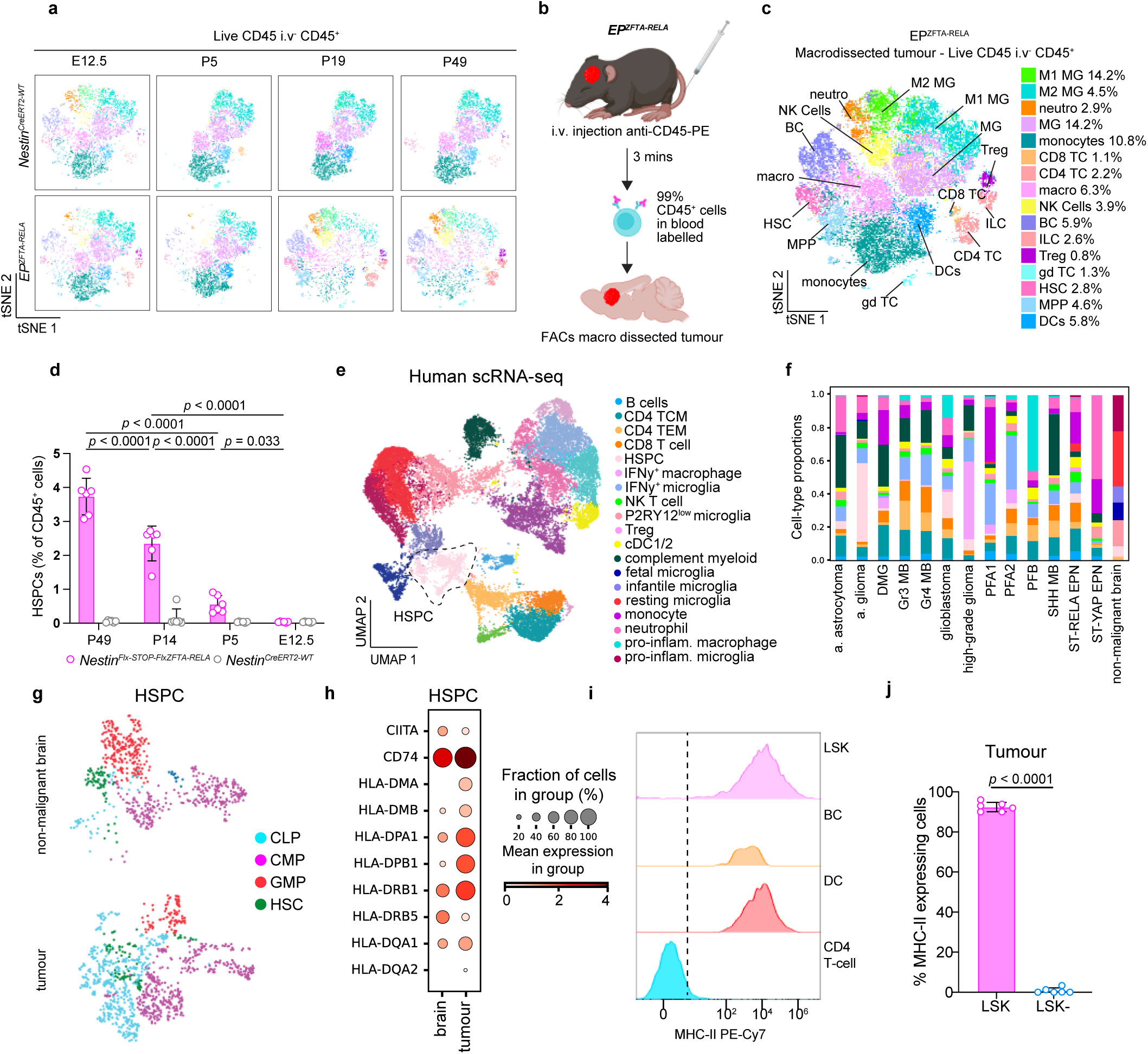
Intratumoural MHC-II^+^ haematopoietic stem progenitor cells (HSPCs) in childhood brain tumours. a, Representative t-SNE of flow cytometry data from developmental time points (E12.5, P5, P19, P49), with 3000 events concatenated per sample, manually gated (key shown to right in (c)). **b**, Schematic of the flow cytometry approach for characterizing the tumor microenvironment in a de novo ZFTA-RELA fusion ependymoma model (image created with Biorender.com) **c**, t-SNE analysis of 18,000 events from endpoint EP*^ZFTA-RELA^* mice, showing proportions of CD45^+^ cells; M1 microglia (MG), M2 MG, neutrophils (neutro.), MG, monocytes, CD8 T cells (TC), CD4 TC, macrophage (macro.), natural killer (NK) cells B cells (BC), innate lymphoid cells (ILC), regulatory T cell (Treg), gamma-delta (gd)-TC, haematopoietic stem cell (HSC), multipotent progenitor (MPP) and dendritic cell (DC). **d**, Quantification of LSK^+^ within the CD45^+^ population in EP*^ZFTA-RELA^* and *Nestin^CreERT2^* control mice (n=5; mean±s.e.m; one-way ANOVA with Šídák’s test). **e,** uniform-manifold approximation and projection (UMAP) of human single-cell RNAseq data integrated from fetal brain tissue and tumour tissue from childhood brain tumour patients, coloured by cell type annotations; B cells CD4 tissue central memory (TCM), CD4 Tissue effector memory (TEM). CD8 T cell, HSPCs, interferon-gamma (IFN-y) macrophages, IFN-y microglia, natural killer (NK) T cell, Purinergic Receptor P2Y12 (P2RY12) low microglia, Treg, conventional dendritic cell (cDC) ½, complement myeloid, foetal, infantile and resting microglia, monocyte, neutropjil and pro-inflammatory (pro- inflam.) macrophage and microglia. **f,** Quantification of the proportion of cell types across each disease group; anaplastic astrocytoma (a.astrocytoma), anaplastic glioma (a.glioma), diffuse intrinsic pontine glioma (DMG), group 3 (Gr3) and group 4 (Gr4) medulloblastoma (MB), posterior-fossa -type A (PFA) ependymoma type 1 and 2, posterior-fossa -type B (PFB) ependymoma, sonic-hedghog (SHH) medulloblastoma, Supratentorial-REL-associated protein (ST-RELA) ependymoma (EPN), ST-Yes1 associated transcriptional regulator (YAP) EPN. **g**, Dotplot of average and percentage expression of MHC-II antigen presentation machinery across HSPC cell clusters in malignant and non-malignant brain tissue. **h**, UMAP of HSPC clusters in malignant and non-malignant brain tissue, coloured by cell type annotation; common lymphoid progenitor (CLP), common myeloid progenitor (CMP), granulocyte-monocyte progenitor (GMP) and HSC. **i,** Representative histogram of MHC-II cell-surface expression in LSKs, BC, DCs and CD4 T cells. **j**, Quantification of the proportion of MHC-II expression in intratumoural LSK^+^ relative to LSK^-^ (n=6/group, mean±s.e.m, unpaired two-tailed Student’s t-test).

HSPCs in human foetal brain and childhood brain tumours expressed major the histocompatibility complex class II (MHC-II) and regulatory genes including *CITTA*, *HLA-DPA1*, *HLA-PB1*, *HLA-DRB1, HLA-DRB5, HLA-DQA1, HLA-DQA2* and the antigen-loading chaperone *CD74* (**Fig. 1g,h**). Similarly, >90% of HSPCs in EP*^ZFTA-RELA^*tumours, but not LSK^-^ committed progenitors, expressed MHC-II at levels similar to professional antigen-presenting cells (APCs; **Fig. 1i,j**). MHC-II expressing HSPCs were also observed in the tumour parenchyma, skull bone marrow and dura mater resected from a child with human choroid plexus papilloma (**Supplementary Fig. 4a**). EP*^ZFTA-RELA^*-resident HSPCs also expressed the antigen loading machinery at levels 6-7-fold higher than in LSK^-^ cells (**Extended data Fig. 1a; Supplementary Table 4**). HSPCs with antigen presentation capacity have been described previously in the peripheral bone marrow where they can eliminate pre-malignant HSCs^31^; however, whether HSPCs with antigen presentation capacity exist in solid tissues and malignancies, including the brain, is not known.

To better understand the characteristics of brain tumour-resident MHC-II^+^ HSPCs, we first tested their long-term self-renewal capacity. Lineage-depleted, CD45.1^+^ MHC-II^+^, but not CD45.1^+^ MHC-II^-^, cells isolated by fluorescence activated cell sorting (FACS) from EP*^ZFTA-RELA^*tumours reconstituted haematopoiesis for up to 16 weeks in non-myeloablative busulfan conditioned (NMC) mice carrying the CD45.2 allele, confirming the stem cell credentials of brain tumour-resident MHC-II^+^ HSPCs (**Extended Data Fig. 1b-e**).

Given the anatomical proximity of EP*^ZFTA-RELA^* tumours to the skull bone marrow, we reasoned that local haematopoiesis in the skull may contribute immune progenitors to the tumour microenvironment. Indeed, skull bone marrow has been implicated as a reservoir of immune cells in non-malignant brain pathologies^32,33^. In keeping with this notion, flow cytometric analysis of mice injected with 5-ethynyl- 2’-deoxyuridine (EdU), detected significantly more EdU-labelled HSPCs as well as total CD45^+^ cells in the skull, but not tibial (hereon, peripheral), bone marrow of postnatal EP*^ZFTA-RELA^*-bearing mice relative to controls (**Supplementary Fig. 5a,b**). Significantly more EdU-labelled monocytes, macrophages and immature B cells were also found in the skull, but not peripheral, bone marrow of EP*^ZFTA-RELA^*-bearing mice (**Supplementary Fig. 5c-d**). We extended these findings to two additional mouse models of childhood brain tumours–choroid plexus carcinoma and Group-3 medulloblastoma^34,35^–where we identified significantly more EdU-labelled HSPCs in the skull bone marrow, relative to sham control (**Supplementary Fig. 5e-f**). Thus, the presence of EP*^ZFTA-RELA^* tumours appeared to selectively engage the skull bone marrow niche, activating local haematopoiesis.

To characterize transcriptional changes within haematopoietic lineages in the presence of an EP*^ZFTA-RELA^* tumour, we generated scRNA-seq profiles of extravascular CD45^+^ cells isolated from the skull, dura, deep-cervical lymph nodes (dCLNs), tumour/brain and peripheral bone marrow of EP*^ZFTA-RELA^*-bearing mice (n=52,140 cells) and compared these with those in controls (n=72,935 cells; **Supplementary Fig. 6a-i**). scRNA-seq profiles of skull bone marrow HSPCs, macrophages, neutrophils, monocytes and Tregs isolated from EP*^ZFTA-RELA^-*bearing mice were enriched for chemotaxis, cytokine signalling, and myelopoiesis gene programs relative to controls (**Supplementary Fig. 6h-i; Supplementary Tables 5-20**). These changes were absent in peripheral bone marrow and dCLNs, underscoring a spatially confined reprogramming of immune activity within central nervous system (CNS)-proximal niches. Thus, the presence of an EP*^ZFTA-RELA^* tumour appeared to activate HSPCs in the skull bone marrow, potentially biasing haematopoiesis toward myelopoiesis and promoting chemotaxis.

### Brain-tumour-derived CSF cues inform skull haematopoiesis

CSF-borne proteins have been shown to educate the skull bone marrow to regulate CNS immune responses^11,17,36^. Therefore, we asked if EP*^ZFTA-RELA^* derived cues might similarly modulate local haematopoiesis through the CSF.

In keeping with this notion, two independent proteomic profiling approaches (multiplex Luminex® and tandem mass tag mass spectrometry) detected significantly higher levels of myelopoietic (e.g., G-CSF, GM-CSF) and chemotactic (MIP-1a/b), as well as T cell activation/polarisation (IL12/23, IL-4) cytokines in the CSF of EP*^ZFTA-RELA^*-bearing mice relative to controls (**Fig. 2a**; **Extended Data Fig. 2a**). Integration of these CSF profiles with skull bone marrow CD45^+^ cell scRNA-seq profiles, identified potential CSF cytokine cross-talk with receptors on HSPCs, monocytes, neutrophils and B cells that could direct leukocyte migration, adhesion, integrin signalling and phagocytosis (**Extended Data Fig. 2b-e; Supplementary Table 21-24**).

**Fig 2:**
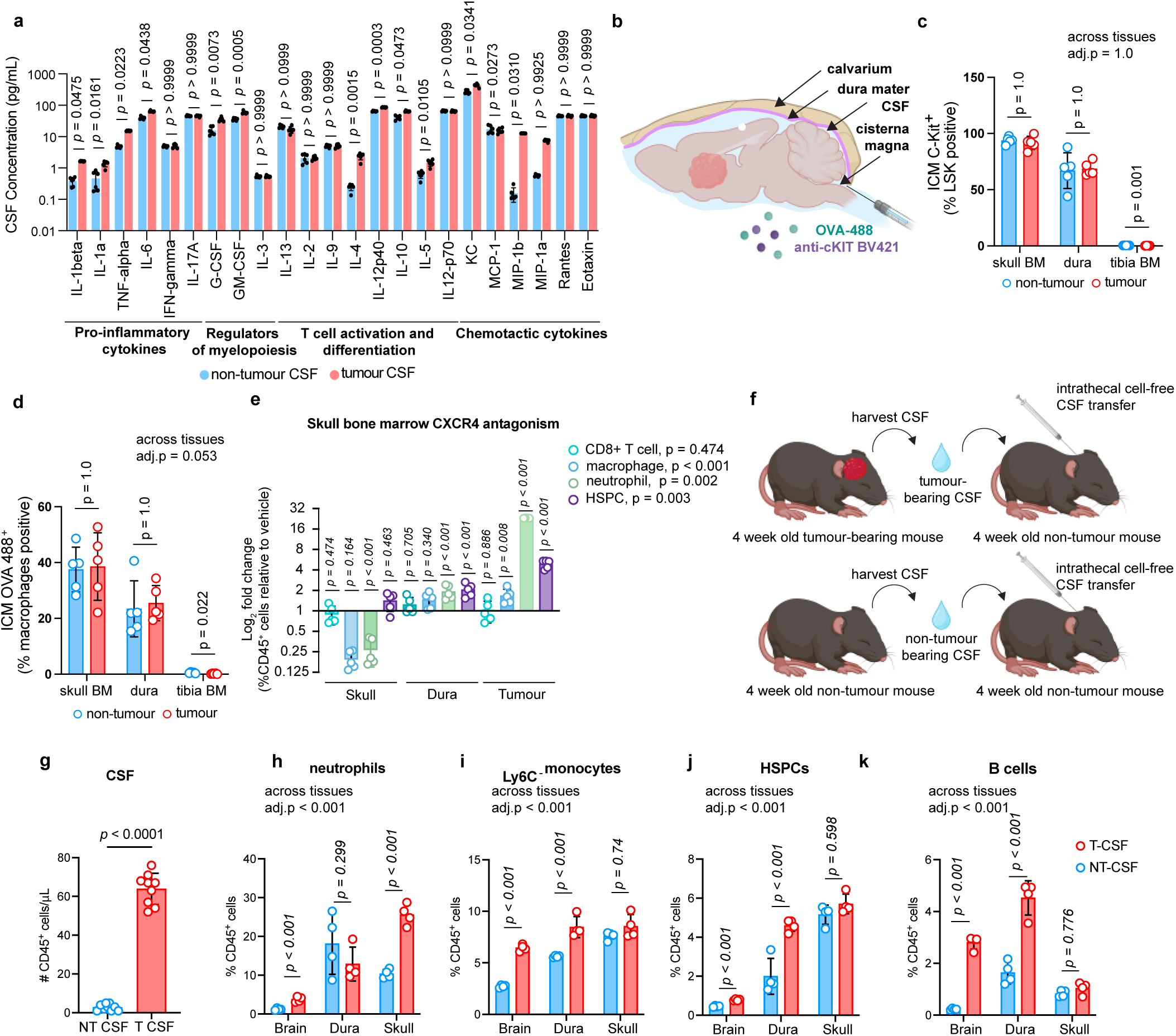
S**k**ull **bone marrow cells access brain-derived solutes via the CSF and supply CNS tumours with HSPCs and myeloid cells**. **a**, Multiplexed measurement of cytokines and chemokines in the CSF of EP*^ZFTA-RELA^*-bearing and control mice using Luminex. n = 6 per group, data are means ± S.E.M. *p* values represent two-sided t tests with Holm-Sidak’s multiplicity adjustment. **b**, Experimental design of intracisterna magna (ICM) injections of anti-c-Kit BV421 and OVA–488 into the cerebrospinal fluid (CSF) tumor and non-tumor-bearing mice (n=5 per group); image created with Biorender.com. **c-d**, Quantification of ICM-injected c-Kit+ Lin-Sca-1+c-Kit+ cells and ovalbumin (OVA)+ macrophages in skull, tibia, and dura (n=5; mean±s.e.m; linear mixed-effects model with Wald z-tests and Bonferroni correction). **e**, Log_2_-fold change of the proportion of intratumoral macrophages, CD8 T cells, haematopoietic stem progenitor cells (HSPCs) and neutrophils following intracalvarial AMD3100 (2 mg/kg, 6 h) or artificial CSF (aCSF) treatment (n=5/group, mean±s.e.m linear mixed-effects model with Wald z-tests and Bonferroni correction) relative to aCSF). **f**, Design for (**g-k**): CSF transfer from tumor or non-tumor mice (n=4/group). **g**, Quantification of CD45+ cells in the CSF (mean±s.e.m; students t-test). **h-k**, Quantification of CD45+ neutrophils (**h**), Ly6C- monocytes (**i**), HSPCs (**j**), and B cells **(k)** in dura, tibia, and skull (mean±s.e.m; linear mixed-effects model with Wald z-tests and Bonferroni correction).

To test more directly if CSF-borne signals could be carried to skull bone marrow HSPCs in our mice, we injected an anti-c-KIT antibody into the intrathecal space and looked for labelling of HSPCs (**Fig. 2b-c; Supplementary Figure 7a-d**). Within two hours of intrathecal injection, 85% and 60% of skull bone marrow and dural HSPCs, respectively, were labelled with anti-c-KIT regardless of tumour presence (**Fig. 2c**). Using a separate approach, we also show that fluorescence-tagged ovalbumin (OVA), injected intrathecally labelled F4/80^+^ CD64^+^ skull bone marrow and dural macrophages (**Fig. 2d; Supplementary Figure 7e**). HSPCs and macrophages in the tibial bone marrow were unlabelled.

Finally, we took two approaches to test if brain tumour-derived signals carried in the CSF might direct skull bone marrow haematopoiesis and promote the migration of skull-derived immune cells to the underlying tumour. First, we injected AMD3100–a CXCR4 antagonist that regulates leukocyte trafficking–directly into the skull bone marrow or intrathecal space of EP*^ZFTA-RELA^* -bearing mice and quantified immune cell populations in the brain, dura and skull bone marrow by flow cytometry (**Fig. 2e; Supplementary Fig. 7f**). AMD3100, but not control, treatment significantly increased the number and proportion of macrophages, neutrophils and HSPCs in EP*^ZFTA-RELA^* tumours and dura mater, while these populations decreased in the skull bone marrow. Once again, no such changes were seen in the blood or peripheral bone marrow. The number of intra-tumoral and dural CD8^+^ T cells was unaffected by AMD3100, consistent with a blood-trafficked origin or CXCR4-independent migratory pathway for these cells (**Fig. 2e**)^37,38^.

Second, to test if signals present specifically in the CSF of EP*^ZFTA-RELA^*-bearing mice can educate the skull bone marrow, we transferred cell-free CSF from EP*^ZFTA-RELA^*-bearing or control mice into the intrathecal space of naïve, age- and sex-matched C57BL/6 recipients and quantified changes in immune cell populations using flow cytometry (**Fig. 2f**). Six hours after CSF transfer, the total number of CD45^+^ cells was significantly increased in the CSF of mice receiving EP*^ZFTA-RELA^*-donor CSF relative to controls, suggesting the CSF is a likely route for tumour-driven skull-brain trafficking (**Fig. 2g**). This was associated with an increased number of neutrophils, Ly6C^-^ monocytes, HSPCs and B cells in the brains of mice receiving CSF from EP*^ZFTA-RELA^*-bearing donors relative to control (**Fig. 2h-k**).

Together, these data support the hypothesis that EP*^ZFTA-RELA^*-derived signals are carried in the CSF to the skull bone marrow, promoting the mobilization of HSPCs and myeloid cells to the dura and tumour.

### Dural sinuses are hubs of anti-tumour immunosurveillance

During normal brain homeostasis, infection, ageing and neuroinflammation, the dura plays a central role in CNS immune surveillance^3,39–41^. In particular, CSF-derived antigens drain through meningeal lymphatics to facilitate presentation to T cells in the dura mater and draining lymph nodes^17,20^.

Therefore, we looked to see if the dura might be a site of immune activation in EP*^ZFTA-RELA^*-bearing mice. We observed a pronounced accumulation of lymphoid aggregates surrounding the confluence of the sinuses, bridging veins and rostral-rhinal hub–sites of known lymphatic-CSF interface^3^–in EP*^ZFTA-RELA^*- bearing mice. These aggregates contained dense infiltrates of CD4^+^ and CD8^+^ T cells, IBA1^+^ macrophages, and MHC-II^+^ APCs which were significantly expanded relative to control mice (**Fig. 3a- h**). Lymphoid dural aggregates in EP*^ZFTA-RELA^*-bearing mice were also characterised by significant increases in IFNγ expressing CD4^+^ T cells, T_H_1 and Treg cells and increased IFNγ secretion was confirmed in fresh dural explants (**Fig. 3i-k)**. Furthermore, single cell T cell receptor (TCR) analysis identified expanded CD4^+^ and CD8^+^ T cell clones that were common to the skull bone marrow and dura, suggesting coordinated anti-tumour immune responses between these two sites (**Extended Data Fig. 3a-i**).

**Fig. 3:**
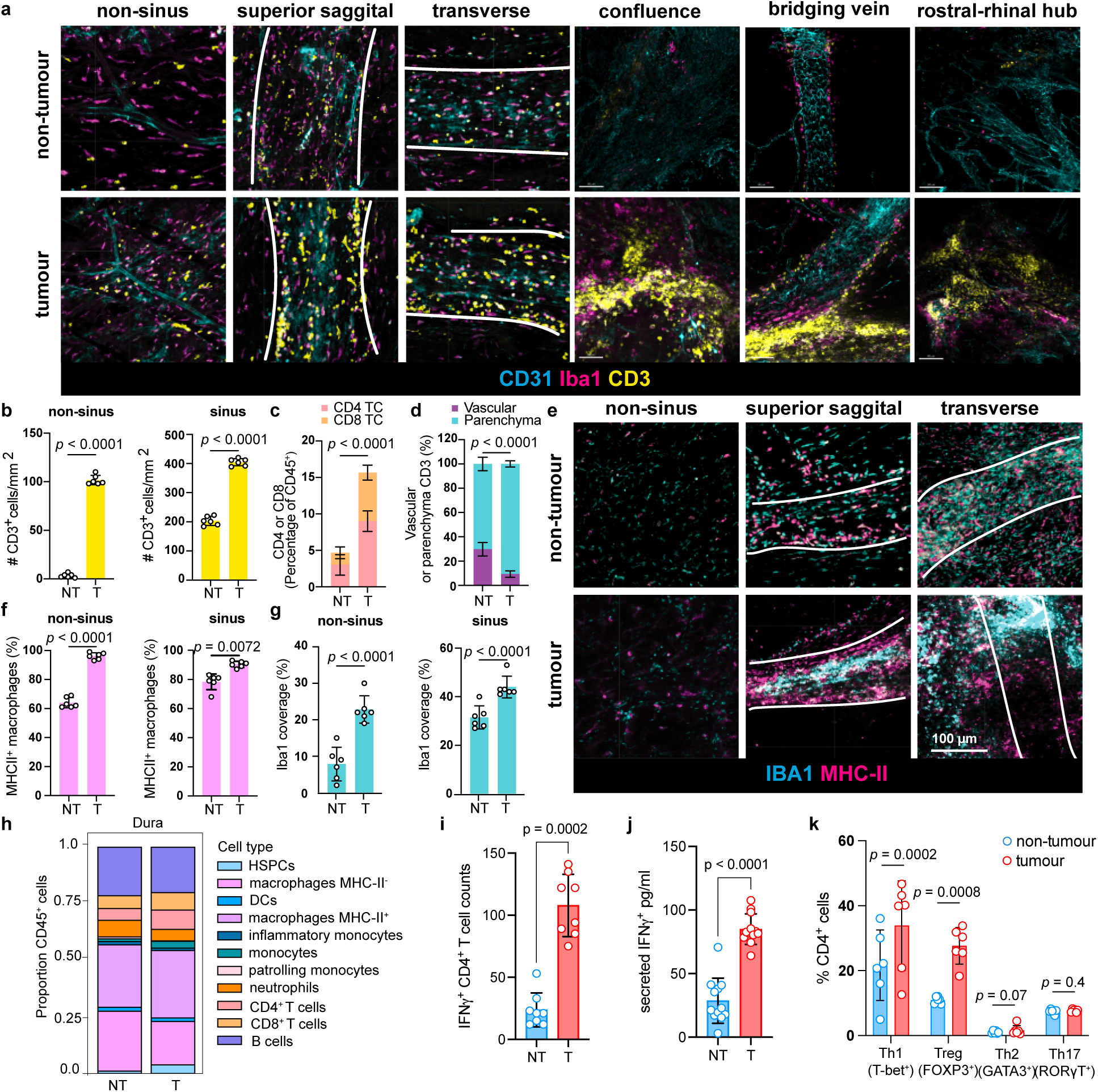
D**u**ral **sinuses are regional hubs for meningeal anti-tumour immunosurveillance. a**, Immunohistochemistry of CD3^+^ T cells, Iba1^+^ macrophages, and CD31^+^ endothelium, at key regions of immunosurveillance in the dura mater in tumour bearing, and control bearing mice. **b-c**, Quantification of the number and proportion of CD3^+^, CD4^+^ and CD8^+^ T cells at dural sinus and non-sinus in *EP^ZFTA-RELA^*-bearing and control bearing mice. **d**, Flow cytometry quantification of the proportion of vascular (CD45 i.v^+^) or parenchyma (CD45 i.v^-^) CD3^+^ T cells in *EP^ZFTA-RELA^*-bearing and control bearing mice, n = 6. **e,** Immunohistochemistry of MHC-II*^+^* Iba1^+^ macrophages in sagittal sinus, transverse sinus and non-sinus regions of the dura mater. **f-g**, Quantification of the proportion of Iba1^+^ cells that co-express MHC-II in the non-sinus and peri-sinus regions in *EP^ZFTA-RELA^*-bearing mice (n=6/group, mean±s.e.m, unpaired two-tailed Student’s t-test). **h**, Flow cytometry analysis and quantification of the proportion of immune cell types in the dura (n = average of 12 mice per group). **I**, Quantification of meningeal CD4+ IFN-y+ T cells in tumour and control bearing mice (n = 8 mice per group, 3 independent experiments) **j,** Quantification of IFN-y in culture supernatants following *ex vivo* stimulation of dural wholemounts (n = 10 mice per group, 2 independent experiments). **k,** Flow cytometry analysis and quantification of T cell phenotypes; T-helper (Th)1, regulatory T cell (Treg), Th2 and Th17 cells in the meningeal dura mater of *EP^ZFTA-RELA^*-bearing and control bearing mice (n = 6 mean±s.e.m.; one-way ANOVA with Šídák’s test).

HSPCs (Sca-1^+^Kit^+^CD150^+^) were also enriched in EP*^ZFTA-RELA^*-bearing dura relative to control mice (**Extended Data Fig. 4a,b**). In contrast to lymphoid aggregates that formed around venous sinuses, HSPCs were located adjacent to non-sinus blood vessels suggesting that in contrast to skull HSPCs, dural HSPCs might monitor peripheral signals. Direct contact between APCs and CD3^+^ T cells was significantly more frequent in the dura of EP*^ZFTA-RELA^*-bearing than control mice, and scRNA-seq profiles of CD45^+^ cells isolated from the dura of EP*^ZFTA-RELA^*-bearing mice were significantly more enriched for regulators of antigen receptor signalling and T cell differentiation pathways relative to control mice (**Extended Data Fig. 4c,d; Supplementary Tables 7, 8**).

Thus, like observations in the skull bone marrow, the immune composition and landscape of the dura was markedly altered by the presence of an EP*^ZFTA-RELA^* tumour, supporting the notion that the dura serves as a site of tumour antigen engagement.

### Brain tumours drive Treg polarisation in the skull bone marrow

Having established that MHC-II^+^ HSPCs in EP*^ZFTA-RELA^*-bearing mice are pluripotent, likely derived from the skull bone marrow, and potentially regulated by CSF signals, we turned our attention to establishing if they possessed antigen-presenting capacity. First, we made use of the Y-Ae monoclonal antibody that specifically recognises presentation of the exogenous Eα_52-68_ peptide presented on MHC- II allele I-A^b^ ^42^ (**Fig. 4a**). EP*^ZFTA-RELA^*-bearing, or control, mice were injected intrathecally with Eα- peptide and its presentation by MHC-II expressing cells quantified in the skull bone marrow, dura and tumour by Y-Ae-based flow cytometry. As expected, CD11c^+^ dendritic cells, but not CD4^+^ T cells, in EP*^ZFTA-RELA^* tumours efficiently presented the Eα-peptide via MHC-II (**Fig. 4b**). As previously described for HSPCs with APC capacity in the peripheral bone marrow^31^, HSPCs in the tumour and the skull bone marrow of EP*^ZFTA-RELA^*-bearing mice also efficiently presented the MHC-II restricted Eα-peptide and in a further functional assay showed these cells efficiently processed DQ-OVA (**Fig. 4b-d**). This antigen presentation could be blocked *in vitro* by pre-incubating HSPCs isolated from EP*^ZFTA-RELA^* tumours with anti-MHC-II antibody prior to treatment with Eα-peptide, confirming that these cells present exogenous peptides via MHC-II (**Extended Data Fig. 5a-c**).

**Fig. 4:**
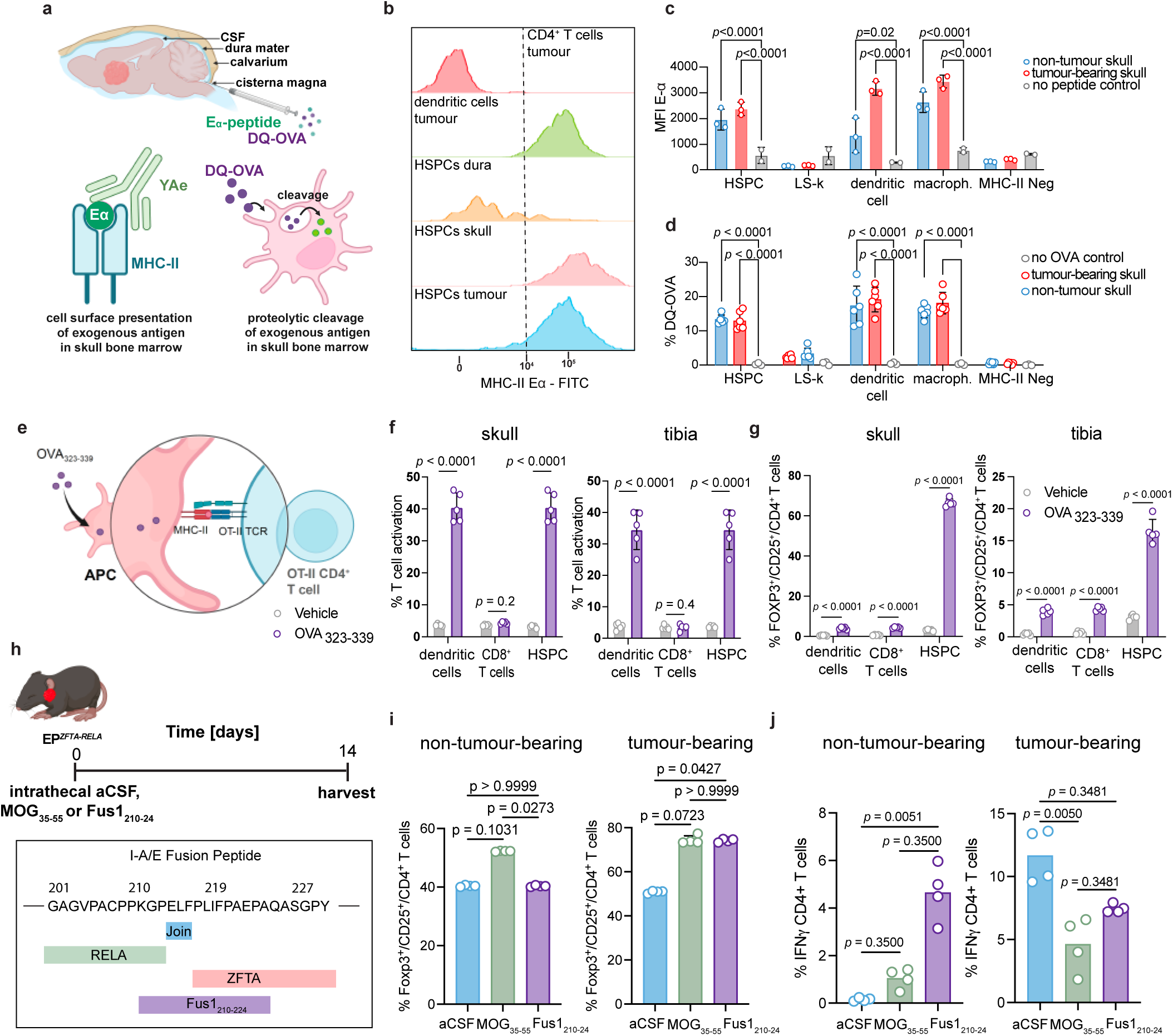
A**n**tigen **presentation of central nervous system and tumour neoantigen drives Treg polarisation in the skull bone marrow a,** Experimental design for profiling antigen uptake and processing via intracisternal magna (ICM) injections of exogenous peptides, created with Biorender.com. **b,** Flow-cytometry histograms of MHC-II Eα immunoreactivity in FACS-isolated CD4^+^ T cells, B cells, dendritic cells (DCs), and HSPCs from tibia and skull bone marrow after Eα peptide immunization (500 µg/mL i.v., 5 h). **c,** Quantification of MHC-Eα^+^ cells by genotype with or without anti-MHC-II antibody (n=3, 10 mice/replicate; mean±s.e.m.; one-way ANOVA with Šídák’s test). **d**, Proportion of DQ-OVA^+^ cells in tumor and non-tumor mice following ICM injection of DQ-OVA for 2 h (n=5, 10 mice/replicate; mean±s.e.m.; one-way ANOVA with Šídák’s test). **e**, Experimental design for *ex vivo* peptide pulsing with vehicle or OVA_323-339_ created with biorender.com. **f**, T-cell activation (CD44^+^ cells) after co-culture of skull/tibia-derived DCs, CD8^+^ T cells, and HSPCs with OT-II CD4^+^ T cells (n=5, 10 pooled mice/replicate; mean±s.e.m.; one-way ANOVA with Šídák’s test). **g**, FOXP3 expression in vehicle or OVA-immunized DCs, CD8^+^ T cells, and HSPCs after OT-II CD4^+^ T cell co-culture (n=5). **h**, Experimental design for intrathecal injection of aCSF, MOG*35-55* or Fus1*210-24* created with biorender.com. **i**, FOXP3 expression in skull CD4 T cells in immunised mice 14 days after injection with aCSF, MOG_35-55_ or Fus1*210-24* . **j**, Quantification of skull CD4*^+^* IFN-y*^+^* T cells following intrathecal injection in tumour or control bearing mice (n = 4 per group, 2 independent experiments).

A defining feature of APCs is their capacity to activate CD4^+^ T cells in an antigen-specific manner; therefore, we looked to see if HSPCs in EP*^ZFTA-RELA^*-bearing mice might present antigens to CD4^+^ T cells. To do this, we made use of CD4^+^ T cells isolated from OT-II mice that specifically recognise the ovalbumin peptide (OVA_323–339_) in the context of MHC-II^43^ (**Fig. 4e**). Skull bone marrow HSPCs, dendritic cells, and CD8^+^ T cells were isolated by FACS from EP*^ZFTA-RELA^*-bearing and control mice and incubated *ex vivo* with OVA_323–339_. HSPCs and dendritic cells, but not CD8^+^ T cells, robustly activated OT-II CD4^+^ T cells *in vitro,* regardless of tumour presence, at levels similar to that achieved by these cells isolated from tibial bone marrow (**Fig. 4f; Extended Data Fig. 5d**). T cell activation correlated with increasing T cell:HSPC ratios and could be abrogated by pre-incubating HSPCs with anti-MHC- II antibody, confirming the MHC-II dependency of this interaction (**Extended Data Fig. 5e-g**). Direct HSPC-CD4^+^ T cell interaction was required for HSPCs to activate T cells (**Extended Data Fig. 5h**). To ensure that skull antigen presentation by HSPCs was not restricted to OVA-OT II interactions, we confirmed activation of CD4^+^ T cells in an MHC-II-dependent manner using FACS-isolated HSPCs inoculated with a different, CNS antigen (myelin oligodendrocyte glycoprotein [MOG]_35-55_; **Extended Data Fig. 5i-j)**. Thus, like HSPCs with APC capacity previously identified in long bones^43^ we show that HSPCs with APC capacity exist in postnatal skull bone marrow; however, these cells appear to exist in the postnatal brain only in the presence of a brain tumour.

To determine how antigen-presentation by HSPCs might impact tumour surveillance, we looked to see if this resulted in the polarisation of naive CD4^+^ T cells into proinflammatory or immunosuppressive T helper subsets. OVA_323–339_ and MOG_35-55_ HSPCS-primed *ex vivo*, that we isolated from the skull bone marrow and tumour, potently upregulated FOXP3 expression in OT-II and 2D2 T cells, indicating Treg polarisation (**Fig. 4g; Extended Data Fig. 5k-m**). Furthermore, scRNA-seq profiles of CD45^+^ cells isolated from the skull and dura of EP*^ZFTA-RELA^*-bearing mice were significantly enriched for genes associated with Treg cell polarisation relative to control mice (**Supplementary Fig. 6h; Supplementary Tables 19-20**). HSPCs exhibited low-to-moderate expression of conventional costimulatory molecules, but markedly elevated levels of the coinhibitory receptor PD-L1 (**Supplementary Fig. 6i**). Furthermore, comprehensive flow cytometry profiling of central and peripheral bone marrow sites in EP*^ZFTA-RELA^*-bearing and control mice revealed selective enrichment of FOXP3^+^ CD4^+^ T cells in the skull bone marrow of EP*^ZFTA-RELA^*-bearing mice (**Extended Data Fig. 5n,o**). These data suggest that skull bone marrow-derived HSPCs act as APCs, but the expression of co-inhibitory molecules, such as PD-L1 modulate T cell responses towards immunosuppression.

To test directly if CD4^+^ T cells in the skull bone marrow are polarised toward Tregs in response to brain-derived peptides *in vivo*, we utilised an adoptive transfer model in RAG2-knock out (KO) mice that lack mature B and T cells (**Extended Data Fig. 6**). CD90.1^+^CD4^+^FOXP3^-^ cells were adoptively transferred into RAG2-KO mice then injected intrathecally at days 14 and 17 post transfusion with either artificial CSF (aCSF; vehicle) or 10 μg of MOG_35-55_. Donor cells isolated 38 days post-infusion from the dura and skull of MOG_35-55_ injected mice demonstrated a marked increase in CD4^+^ T cell polarisation toward FOXP3^+^ Tregs relative to aCSF-treated mice (**Extended Data Fig. 6b-e**). No such polarisation was observed among cells harvested from the spleen, blood or brain. We were able to corroborate this effect in wild-type mice, demonstrating that CNS-derived peptides could also drive the polarization of conventional CD4^+^ T cells into Tregs within the skull (**Extended Data Fig. 6 f-i**).

Given the response of naïve CD4^+^ T cells to CNS antigens in the skull bone marrow, we reasoned that CNS tumour neoantigens may favour the same immunosuppressive response. To test this, we identified a MHC-II (I-A/E) predicted neoantigen unique to the ZFTA-RELA fusion protein in our EP*^ZFTA-RELA^* mouse model and inoculated EP*^ZFTA-RELA^*-bearing and control mice with this peptide (Fus1_360-71_), MOG_35-55_ or aCSF intrathecally (**Fig. 4h; Supplementary Table. 25**). Consistent with a classical response to a foreign antigen in control mice, Fus1_210-24_ failed to promote Treg polarisation in the skull bone marrow and favoured a Th1 response indicated by elevated IFNγ^+^ CD4^+^ T cells (**Fig. 4i,j**). Remarkably, mice harbouring EP*^ZFTA-RELA^* tumours inoculated with Fus1_210-24_ mirrored the response of mice inoculated with the CNS self-peptide MOG_35-55_, including Treg polarisation and a lack of IFNγ^+^ CD4^+^ T cell induction. Hence, in the context of EP*^ZFTA-RELA^*, the fusion neoantigen is recognised as a self-antigen by the developing skull bone marrow. In further support of this notion, only 4.8% of EP*^ZFTA-RELA^* tumour cells engrafted in the brains of syngeneic adult C57BL/6 mice compared to an engraftment rate of 98.5% in perinatal pups (χ²=96.685, p<0.00001, and data not shown), strongly supporting the notion that local and early immunotolerance is important in tumorigenesis. Thus, skull HSPCs and other antigen-presenting cells take up, process and present CSF antigens via MHC-II, activating and polarizing T-cells to provide a local supply of immunosuppressive Tregs.

### Ependymoma drives aberrant myelopoiesis in the skull bone marrow

In addition to subverting HSPC-T cell interactions, we hypothesised that the presence of a brain tumour may also reshape skull bone marrow HSPC lineage commitment. As a first step to test this, we interrogated the regulatory mechanisms underpinning haematopoiesis using single-nuclear assay for transposase-accessible chromatin with sequencing (snATAC-seq) and transcriptomic analysis of CD34^+^LSK^+^ cells that we FACS-isolated from the skull and peripheral bone marrows of EP*^ZFTA-RELA^*- bearing and control mice. UMAP projections of chromatin accessibility landscapes revealed discrete clustering of hematopoietic subpopulations, with skull HSPCs from EP*^ZFTA-RELA^*-bearing mice displaying a marked shift toward myeloid fate commitment relative to controls (**Fig. 5a-b; Supplementary Fig. 8a-f**). Pseudotime analysis further supported a trajectory marked by reduced lymphoid output and dominance of myeloid programs (**Fig. 5c,d**). Motif enrichment analysis across pseudotime revealed a myeloid-specification programme that included genes downstream of GM-CSFRα e.g., *Fos*, *Bach1/2*, and *Smarcc1* consistent with elevated GM-CSF levels we had observed in the CSF (**Fig. 5e,f**; **Fig. 2a; Extended Data Fig. 2a**), while gene ontology analysis revealed a concomitant suppression of lymphoid-associated programmes in skull relative to peripheral bone marrow (**Fig. 5g,h**). Together, these data suggest that tumour-derived signals in the CSF reprogram local HSPCs toward a myeloid- skewed state at the expense of lymphopoiesis.

**Fig. 5:**
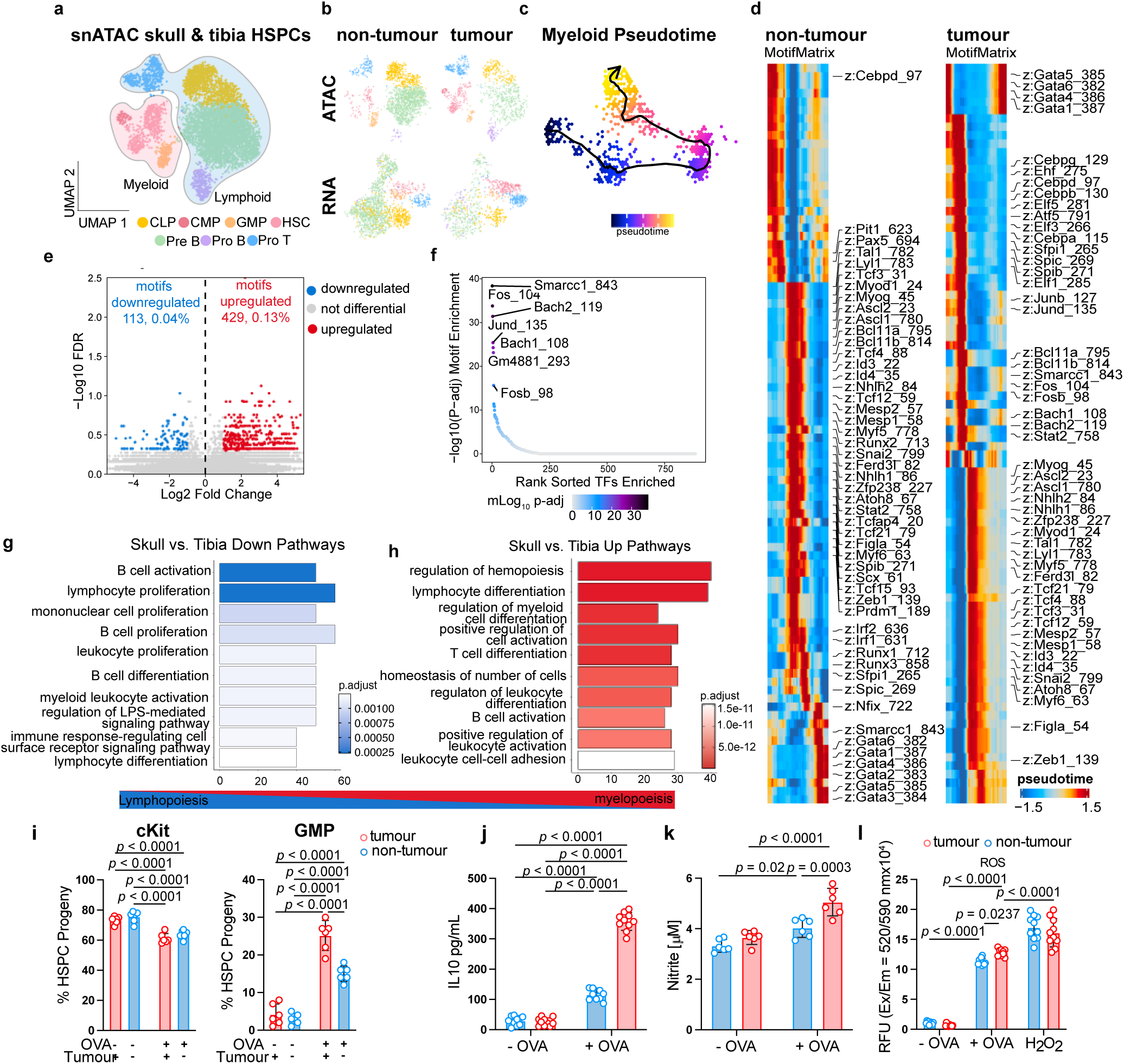
Combined analysis of chromatin accessibility and gene expression in HSCs from skull and tibia of *EP^ZFTA-RELA^*-bearing and control-bearing mice reveals myelopoiesis bias in skull HSCs. a, uniform manifold approximation and projection (UMAP) visualisation of the single nucleus assay for transposase-accessible chromatin using sequencing (snATAC) dataset (1623 nuclei from LSK^+^ CD34^+^haematopoietic stem cells (HSCs) sorted cells from the skull and the tibia of *EP^ZFTA-RELA^*-bearing and control bearing mice), coloured by cluster; common lymphoid progenitors (CLP), common myeloid progenitor (CMP, granulocyte-monocyte precursor (GMP), HSC, Pre B cell, Pro B cell and Pro T cell. **b**, UMAP separated by genotype and modality. **c**, ArchR pseudotime visualisation of the differentiation trajectory of haematopoietic cells from the skull of *EP^ZFTA-RELA^*-bearing mice from. **d**, Heatmap of motifs identified across the myeloid cell trajectory with ArchR split by control skull and tumour-skull. **e**, Motif enrichment in differential peaks upregulated in skull of tumour relative to control bearing mice visualised by volcano plot. **f**, Significant rank-sorted transcription factor motifs enriched in skull HSCs of tumour bearing mice. **g-h**, Gene Ontology (GO) pathway analysis top differentially downregulated and upregulated genes in skull-derived HSCs relative to tiba. **I**, Indicated populations derived from *ex-vivo* ovalbumin (OVA)-primed HSPCs following co-culture with OT-II CD4^+^ T cells (n = 6 mean±s.e.m.; one-way ANOVA with Šídák’s test). **j-l**, Cytometric bead array measurements of IL-10 production, nitrite production and reactive oxygen species (ROS) production in OVA primed HSPC-derived populations following co-culture with OT-II CD4^+^ T cells (n = 10 mean±s.e.m.; one-way ANOVA with Šídák’s test).

To validate the myeloid bias observed in tumour-associated skull HSPCs, we employed two complementary approaches. First, colony-forming unit assays confirmed that HSPCs isolated from the skull bone marrow of EP*^ZFTA-RELA^*-bearing produced significantly more granulocyte-macrophage colonies (CFU-GM) than controls, consistent with a myeloid bias fate (**Supplementary Fig. 8g**). Second, OVA-primed skull HSPCs that we co-cultured with OT-II CD4+ T cells *ex vivo* differentiated toward granulocyte-monocyte progenitor (GMP) and myeloid-lineage cells (**Fig. 5i**). This cellular reprogramming was accompanied by increased production of IL-10, nitrite (NO_2_^-^) and reactive oxygen species (ROS), consistent with a myeloid-derived suppressor cell (MDSC)-like phenotype^44^ (**Fig. 5j-l**).

Finally, we further tested the contribution of HSPC antigen presentation to tumour progression by generating an inducible MHC-II knockout mouse in which H2-Ab1^tm1Koni/J^ mice were bred to SLC- CreER^T2^ mice (**Extended Data Fig. 7a**). Tamoxifen induction at P0 and P1 achieved a 60% knockdown of MHC-II expression on LSK^+^ cells within the skull bone marrow, which was partially restored by P21 (**Extended Data Fig. 7a,b**). The growth of orthotopic EP*^ZFTA-RELA^* tumour allografts in these mice was significantly delayed relative to those in mice with intact MHC-II, resulting in prolonged survival (median survival: 67 days vs. 37.5 days, *p*=0.0025; **Extended Data Fig. 7c-e**).

Together, these findings indicate that EP*^ZFTA-RELA^* tumours reprogram local HSPC chromatin and transcriptional landscapes in favour of myeloid lineages, linking skull marrow haematopoiesis to tumour-induced immune remodelling. Sustained antigen presentation by skull HSPCs promotes myelopoiesis and immune suppression in the context of EP*^ZFTA-RELA^*tumours, identifying this axis as a driver of local immunotolerance.

### CNS immunosurveillance is a therapeutic vulnerability in childhood brain tumours

Given the profound relationship between local immunotolerance and brain tumorigenesis, we looked to see if this might represent a therapeutic vulnerability of childhood brain tumours. We reasoned that elevated GM-CSF in the CSF–likely produced by cells across the tumour, dura, and skull–participates in a circuit in which tumour-derived signals drive aberrant haematopoiesis in adjacent marrow (**Supplementary Fig. 9a,b**). Indeed, our transcriptomic data identified high-level expression of the key myelopoiesis regulator Csf2ra (encoding GM-CSFRα) in the skull bone marrow (**Supplementary Fig. 9c,d**). While not exclusive to HSPCs, this receptor is well positioned to convert local GM-CSF signals into a myelopoietic program^45,46^. Therefore, we randomised mice bearing EP*^ZFTA-RELA^* tumours (confirmed by bioluminescence exploiting the *Nestin^Cre-ZFTA-RELA^*-IRES-Luciferase allele; **Supplementary Fig. 1**) to receive a single intrathecal injection of anti-GM-CSF antibody (5 mg/kg), mavrilimumab (5 mg/kg; an anti-GM-CSFRα antibody) or control antibody (**Fig. 6a**). Remarkably, single-dose antibody blockade of either GM-CSF, or its receptor, induced a near-complete regression of EP*^ZFTA-RELA^* tumours that was sustained for around six weeks and associated with a greater than three-fold increase in survival time (**Fig. 6 b,c; Supplementary Table 28**). GM-CSF targeted therapy also decreased CSF GM-CSF levels, skull HSPC proliferation, and tumour-associated myeloid cells, while increasing tumour CD8+ T cell infiltration increased within 21 days (**Extended Data Fig. 8a-h**).

**Fig. 6:**
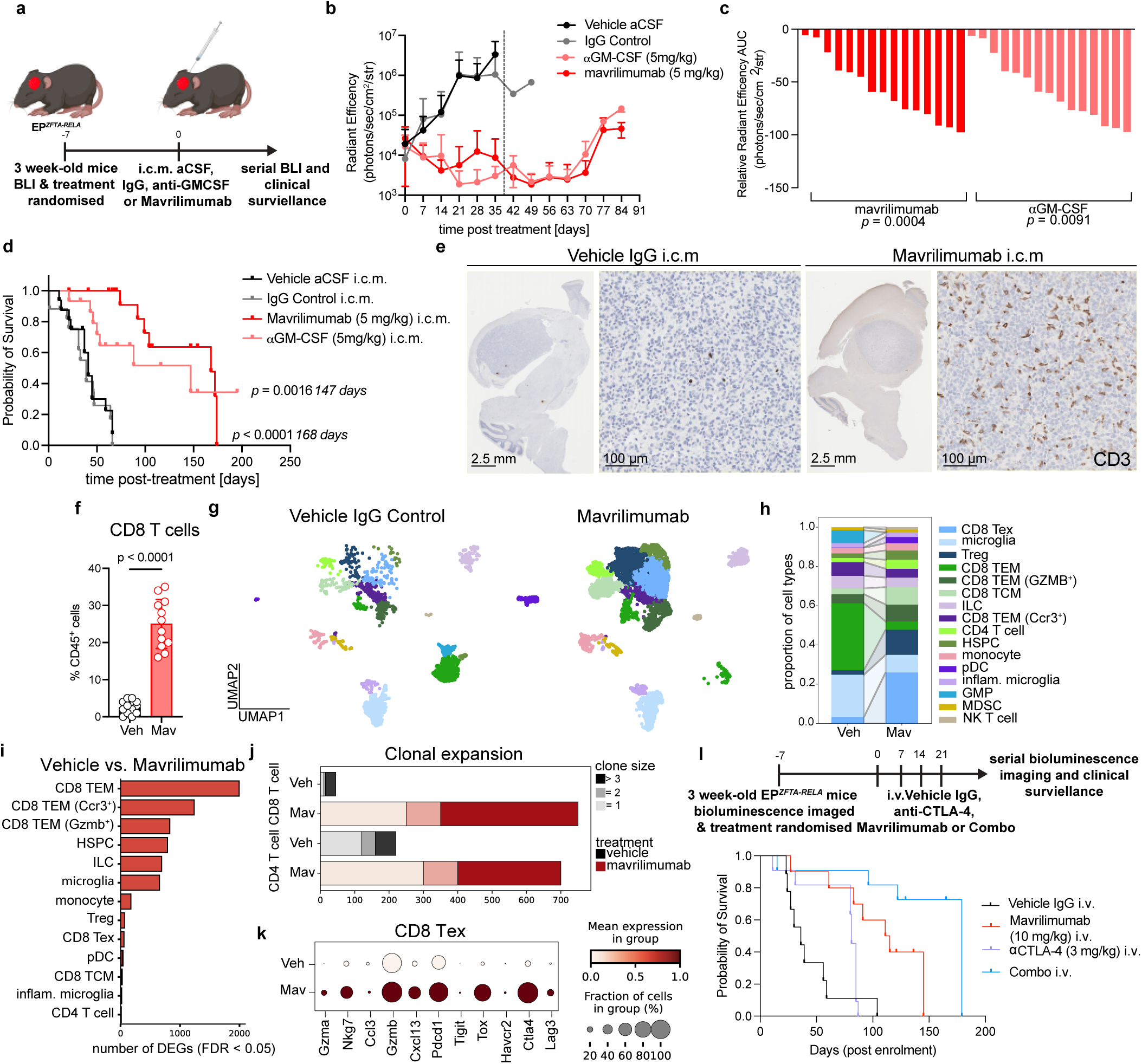
N**o**rmalising **skull haematopoiesis improves the survival of ZFTA-RELA fusion-driven ependymoma-bearing mice. a,** Experimental design for the treatment of 3 week-old ZFTA-RELA *EP^ZFTA-RELA^*-bearing mice with a single intracisternal magna (ICM) injection of 10 uL of Vehicle aCSF (*n* =17), IgG isotype control (5 mg/kg, *n* = 16), anti-GM-CSF (5 mg/kg, *n* = 14) or mavrililumab (5 mg/kg, *n* = 17) created with biorender.com. **b**, Weekly bioluminescence tracking of treated mice. **c**, Waterfall plots of the area under the curve (AUC) for anti-GM-CSF (*p* = 0.0004, *n* = 15) and mavrilimumab- treated mice (*p* = 0.0091, *n* = 16) up to 42 days, Welch’s t-test. **d**, Kaplan–Meier survival plot of *EP^ZFTA-RELA^* -treated mice, log rank Mantel Cox test. **e**, Representative immunohistochemical images of brains from mice treated with vehicle or mavrilimumab at recurrence. **f**, Flow cytometry quantification of CD8 T cells as a proportion of CD45^+^ cells, n = 12 mice per group. **g**, uniform manifold approximation and projection (UMAP) visualisation of single-cell RNAseq of extravascular CD45^+^ cells isolated from the tumour parenchyma of vehicle or mavrilimumab-treated mice at recurrence, coloured by cell type; exhausted CD8 T cell (CD8 Tex), microglia, CD8 tissue-effector memory (TEM), CD8 TEM GZMB+, CD8 tissue central memory (TCM), CD4 T cell, haematopoietic stem progenitor cell (HSPC), monocyte, plasmacytoid (pDC), inflammatory (inflame) microglia, granulocyte-monocyte precursor (GMP), myeloid-derived suppressor cell (MDSC) and natural killer (NK) T cell. **h**, Quantification of the proportion of cell types in each group. **i**, Quantification of the number of differentially expressed genes in each cell type in vehicle relative to mavrilimumab-treated mice. **j**, Single-cell T-cell receptor (TCR) sequencing clonotype analysis of the number of expanded clones in CD4^+^ and CD8^+^ T cells in vehicle and mavrilimumab-treated mice (n = 6-10 tumours pooled per group). **k**, Dotplot of average and percentage expression of exhaustion markers in CD8 Tex population in vehicle and mavrilimumab- treated mice. **l**, Protocol and Kaplan–Meier survival plot of *EP^ZFTA-RELA^* -treated mice, with four, weekly intravenous injections of IgG control (n = 9), mavrilimumab (10 mg/kg, n = 10), anti-CTLA-4 (3 mg/kg, n = 11) or combo (mavrilimumab + anti-CTLA-4, n = 11), log rank Mantel Cox test.

Since we observed evidence of aberrant proliferation in the skull bone marrow of other childhood brain tumours (**Supplementary Fig. 5e-f**), we tested if the GM-CSF signaling axis might represent a therapeutic vulnerability in other aggressive childhood brain tumours. Remarkably, mice harboring accurate models of CPC and Group-3 medulloblastoma^34,35^ also displayed profound tumour growth suppression; reduced CSF GM-CSF levels; decreased skull HSPC and tumour myeloid cell numbers; and increased CD8^+^ T cell infiltration in response to mavrilimumab therapy (**Extended Data Fig. 8i– t**). These data reveal GM-CSF signalling as a key mediator of skull-bone marrow driven myelopoiesis and tumour tolerance in a broad array of childhood brain tumours. To our knowledge, this is the first report of skull-directed therapy for the treatment of an intracranial malignancy.

The rapid tumour regression and increased CD8^+^ T cell infiltration that followed GM-CSF axis blockade suggested skull-derived myeloid derived suppressor cells might impede cytotoxic T cell function. Indeed, immunohistochemistry and flow cytometry of mavrilimumab-treated tumours confirmed elevated CD8^+^ T cells in the tumour parenchyma persisted even in relapsed tumours (**Fig. 6e,f**). Furthermore, TCR profiling of extravascular CD45^+^ cells revealed clonal expansion of CD8^+^ and CD4^+^ T cells within the tumour parenchyma of mavrilimumab-treated mice and gene ontology analysis of CD8^+^ T cell populations revealed upregulation of the type 1 interferon response, regulation of cell killing and T cell activation (**Fig. 6g–j; Supplementary Fig. 10a-h; Supplementary Tables 29-34**). CD8^+^ T cells with high clonal expansion displayed elevated cytotoxicity (GZMK, GZMB), activation (NKG7, CCL5), inflammation and chemotaxis (Ccr5, Xcl1, S100a6), as well as exhaustion (Lilrb4a, Pdcd1; **Extended Data Fig.9**). Global changes in the transcriptome of other intratumoural immune cell subsets–including monocytes, ILCs, microglia–revealed upregulation of antigen processing and presentation, myeloid leukocyte activation and response to interferon beta, suggesting mavrilimumab- driven reprogramming of the tumour microenvironment in favour of T-cell activation (**Supplementary Tables 35-44)**. Consistent with an on-target effect, intratumoural HSPCs downregulated haematopoiesis, lymphocyte differentiation and immune system development programs following mavrilimumab treatment (**Supplementary Table 35-36)**. These data demonstrate for the first time that skull-bone marrow directed therapy can modulate the local supply of MDSCs and induce the clonal expansion and activation of T cells within the tumour parenchyma.

CD8^+^ T cells in relapsed tumours upregulated exhaustion markers (*Ctla4*, *Lag3*, *Pdcd1*, *Tigit*), suggesting that this might explain the subsequent failure of GM-CSF blockade therapy (**Fig. 6k; Supplementary Fig. 10i**; **Supplementary Table 29-46**). Therefore, we tested if an immune checkpoint inhibitor (anti-CTLA-4) might increase the therapeutic benefit of mavrilimumab. To facilitate the clinical translation of this combination treatment regimen, we delivered mavrilimumab (10 mg/kg) intravenously (rather than intrathecally), either alone or with anti-CTLA-4 (3 mg/kg). Intravenous mavrilimumab monotherapy proved highly effective (p < 0.0001, n = 11), but this was further increased by co-treatment with anti-CTLA-4 (p < 0.0223, n = 11) (**Fig. 6l; Supplementary Table 47**). Anti-CTLA-4 alone providing no significant survival benefit (p < 0.4379, n = 11).

Despite intravenous administration, CD45⁺ cell proportions in peripheral tissues (spleen, tibial bone marrow) were unchanged **(Extended Data Fig. 10)**. In contrast, mavrilimumab, alone or in combination, significantly reduced intratumoural monocytes, B cells, neutrophils, and CD4⁺ T cells (all *p* < 0.001), while increasing CD8⁺ T cells in tumour parenchyma (*p* < 0.001) and dura (*p* = 0.037). In the skull bone marrow, we observed reduced macrophages, monocytes, neutrophils, and T cells across both CD4⁺ and CD8⁺ subsets, with interaction effects between compartments suggesting a redistribution of immune cells from bone marrow to dura and tumour **(Extended Data Fig. 10)**. These findings demonstrate that normalizing skull bone marrow haematopoiesis drives profound remodelling of local immune niches, favouring anti-tumour immunity. Importantly, we show on-target activity of intravenous mavrilimumab in combination with immune checkpoint blockade, supporting its translational potential.

Together, these data identify the skull marrow as a druggable immunological niche for children with a brain tumour. The combination of mavrilimumab and anti-CTLA-4 provide a rationale combination treatment that could be readily translated to the clinic to treat children with a broad array of brain tumours.

## Discussion

Historically, the skull bone marrow has been thought of solely as a site of haematopoiesis. But this compartment is connected to CNS border tissues through channels in the skull, enabling CSF to facilitate communication between the brain, meninges and skull bone marrow^20,41,47^. Here, we show for first time that the CSF has an instructional role in shaping skull bone marrow haematopoiesis. Using a spontaneous mouse model of EP*^ZFTA-RELA^* we identify a new type of skull bone marrow HSPC, similar to one previously identified in peripheral bone marrow^43^, that presents CNS antigens via MHC-II. Tumour-derived signals carried in the CSF educated skull bone marrow such that interactions between these HSPCs and CD4^+^ T cells during antigen presentation directed HSPC myeloid-biased differentiation and T cell polarisation to T-regs, contributing to tumour immunotolerance. While this dynamic immunosurveillance system likely evolved to protect the brain from neuroinflammation, we propose that this mechanism is subverted by childhood brain tumours to promote tumour tolerance. Indeed, antibody therapy targeting a tumour-derived cytokine in three very different types of aggressive childhood brain tumour in mice, disrupted these processes to produce profound tumour growth suppression and survival benefit. Therefore, we propose that chronic, tumour-antigen exposure in the skull bone marrow creates sustained tumour tolerance in children with brain tumours and that this represents a novel therapeutic vulnerability. Our observations that the skull bone marrow participates in CNS tolerance through HSPCs antigen presentation and local Treg polarisation extends evidence that CNS-antigen autoreactive B cells are negatively selected in the dura^18,48,49^ and that CNS autoimmunity is regulated by skull bone marrow-derived cells^32,33^.

HSPCs with APC capacity have been described previously in the peripheral bone marrow in the context of leukaemia^31^. In the peripheral bone marrow this process protects the integrity of the stem cell pool by editing out malignant progenitors. In contrast, we show that the *in utero* development of a brain tumour subverts normal antigen-presenting HSPC haematopoiesis in the skull, in part by changing the chromatin and transcriptional landscape of these cells to favour myelopoiesis. While this process involves HSPC-CD4^+^ T cell interactions that educate the local stem cell pool, the broad changes we observed across the dura, CSF and skull bone marrow suggest broader, multi-factorial effects are at work. These findings are consistent with emerging evidence that the skull bone marrow is highly sensitive to intracranial and peripheral cues, including bone resorption-mediated HSPC expansion, exchange of molecules via arachnoid cuff exit points from the subarachnoid space to the dura mater^50,51^ and systemic myelopoiesis induced by non-small cell lung cancer^52^.

In addition to promoting immunotolerance through the differentiation of antigen presenting HSPCs, we show that these cells polarise CD4^+^ T cells toward Tregs that are known to inhibit antitumor immunity. Thus, brain tumour immunotolerance might also be achieved by the subversion of normal CNS- homeostatic mechanisms in which regulatory CNS peptides are presented within the skull and dura to reinforce a population of unconventional suppressor CD4^+^ T cells^13^. We show that this process can be driven by exogenous peptides, endogenous CNS peptides, and the ZFTA-RELA fusion; thereby demonstrating that tumour neoantigens can be recognised as self as part of tumour immunotolerance. In line with recent observation that FOXP3^+^CD4^+^ T_reg_ cells are readily detectable in the dura mater of homeostatic mice and that their ablation led to an enrichment of IFNγ-producing cells, we demonstrate that polarisation of CD4 T cells towards Tregs can occur in the skull bone marrow and dura in response to endogenous CNS and neoantigens^53^. Together these data extend the notion that the dura mater serves as a critical site for immunotolerance, favouring immunosuppression, which is subverted in the context of childhood brain tumours. Further work will be required to define precisely which peptides are active in the tolerance of the different types of childhood brain tumours in patients and how these insights might facilitate vaccine and peptide-based immunotherapeutics.

Targeting skull bone marrow GM-CSF-driven myelopoiesis through intrathecal delivery of anti-GM- CSF or anti-GM-CSFRα resulted in profound tumour growth suppression. Whilst we cannot rule-out modulation of other myeloid cells, the prolonged efficacy achieved with a single intrathecal dose of antibody, coupled with on-target reduced haematopoiesis and concurrent reduction in intratumoural MDSCs, strongly suggests that this treatment targeted HSPC-function. While all mice treated in this manner relapsed this could be mitigated by treatment with anti-CTLA-4, overcoming the exhaustion associated with intratumoural CD8^+^ T cell clonal expansion. The combination of mavrilimumab and anti-CTLA-4 provide a rationale combination treatment that could be readily translated to the clinic to treat children with a broad array of brain tumours. The immunotolerance axis described here may also have implications for the development of other immunotherapies for childhood brain tumour patients. Further exploration of other immune-checkpoint blockades and CAR-T therapy may result in highly effective new therapies that spare the child from the severe neurotoxicity of conventional surgery, chemotherapy and radiation treatment.

In summary, we identify a novel therapeutic vulnerability in childhood brain tumours, where the tumour educates the local immune supply from the skull bone marrow at the apex of haematopoiesis. Our findings demonstrate that CNS-derived signals can regulate skull haematopoiesis to enforce a locally tolerant immune repertoire, fostering an immunosuppressive environment conducive to tumour development.

## Methods

### Materials availability

This study did not generate new unique reagents.

### Experimental model and subject details Human participants

Human brain tissue was obtained from patients undergoing neurosurgical procedures at Cambridge University Hospitals NHS Foundation Trust. Tissue collection and use for research were approved by the East of England–Cambridge Central Research Ethics Committee (REC reference: 23/EE/0241). Written informed consent was obtained from all participants prior to inclusion, in accordance with the Declaration of Helsinki.

### Mice

All animal work was carried out under the Animals (Scientific Procedures) Act 1986 in accordance with the UK Home office license (Project License PP9742216) and approved by the Cancer Research UK Cambridge Institute Animal Welfare and Ethical Review Board. Mice were housed in individually ventilated cages with wood chip bedding plus cardboard fun tunnels and chew blocks under a 12 hour light/dark cycle at 21 ± 2°C and 55% ± 10% humidity. Standard diet was provided with ad libitum water. Mice were allowed to acclimate for at least one week in the animal facility prior to the beginning of any experiment. Adult males and females between 4-6 weeks of age were primarily used for our studies unless stated otherwise. Sample sizes were determined on the basis of a power analysis in accordance with previously published experiments. Experimenters, where necessary, were blinded to experimental groups during both scoring and quantification. Mouse strains used are listed in Supplementary Table 1.

## Method Details

### Generation of Rosa26-locus targeted conditional, Nestin-driven, C11orf95-RELA fusion expressing mice

A conditional C11orf95:RelA fusion (Nestin-(lsl)-C11orf95:RelA) construct with homology to the Rosa26 locus was generated by conventional molecular techniques. The targeting plasmid was linearised and gel purified before being nucleofected into C57BL/6J embryonic stem cells. After G418 sulfate selection clones were isolated and subjected to genotyping and karyotyping. Four correctly targeted clones were injected into wild type CD1 8-cell stage embryos. The microinjected embryos were cultured in KSOM+AA (KCl, enriched simplex optimisation medium with amino acid supplement, Zenith Biotech) at 37°C with 95% humidity and 5% CO_2_ until the blastocyst stage and then were transferred into pseudopregnant recipients. The resulting F0 mice were bred to C57BL/6J mice, proving germline transmission, and establishing the colony (see Supplementary Tables for details on animals, Supplementary Table 1 and oligonucleotide sequence, Supplementary Table 4).

### Orthotopic allotransplantation models

Choroid plexus carcinoma (CPC) and group 3 medulloblastoma orthotopic allotransplantation models were generated from *in vitro* cultures of cells generated from genetically engineered mouse models of each of these tumours. Briefly, as an extension of our previous work^34^ in Tong et al., 2015, CPC cell lines were derived from *Pten*^-/-^*Rb*^-/-^*Tp53*^-/-^ mice, generously donated by Suzzane J. Baker^54^ crossed with TTR CreEsr1;TdT mice (MGI:3046546)^55^, in which tamoxifen induction at P1/2 results in recombination within the choroid plexus and CPC tumours postnatally. Cell lines derived from these tumours were expanded in Neurobasal medium with B27 without vitamin A, N2 Supplement and rEGF, FGFb and IGF-2 (20 ng/mL), on extracellular matrix -coated flasks. Group 3 medulloblastoma lines were generously donated by Fredrik J. Swartling^535^ and were cultured in Neurobasal medium with B27 without vitamin A, N2 Supplement and rEGF, FGFb and IGF-2 (20 ng/mL). Both lines were transfected with luciferase (pLenti CMV Puro LUC (w168-1)) prior to implantation. P1 C57B/L6 mice were orthotopically implanted with 400 cells (G3 medulloblastoma) or 4000 cells (CPC) on the right temporal lobe, or lateral ventricle, respectively under anaesthesia.

### *In vivo* preclinical studies

EP*^ZFTA-RELA^*, CPC and Gr3 medulloblastoma tumour-bearing mice were randomised at 3 weeks old, confirmed by bioluminescence exploiting the endogenous NestinCre-ZFTA-RELA-IRES-Luciferase allele in EP*^ZFTA-RELA^* or the lenti-viral introduction of luciferase *in vitro* in the CPC and Gr3 medulloblastoma as described above. Mice received a single intrathecal injection of mavrilimumab (5 mg/kg; an anti-GM-CSF receptor alpha antibody) or control antibody, and were monitored with bi- weekly bioluminescence imaging to monitor tumour burden until mice reached a humane endpoint based on the accumulation of clinical signs. For intravenous administration, EP*^ZFTA-RELA^* bearing mice were randomised at 5 weeks old to receive mavrilimumab (10 mg/kg), alone or with anti-CTLA-4 (3 mg/kg) or control antibody, once-weekly for 4 weeks.

### Tissue Processing and immunohistochemistry

Mice were given a lethal dose of anaesthesia via IP (Dolethal 10% v/v) and transcardially perfused with 0.025% heparin PBS. Mice were decapitated posterior to the occipital bone and following removal of overlying skin and muscle from the skull, the skull cap was removed and drop-fixed in paraformaldehyde (4% w/v in PBS) for 1 hour at 4 °C. Duras were peeled from the skull and washed three times in PBS before being mounted on Superfrost Plus slides (Fisher Scientific). For blocking and permeabilization, wholemounts were incubated in buffer containing 10% Tris, 1% BSA (R&D Systems, Cat# 841380), 1% serum, 1% saponin, and 0.5% Triton X-100 in MilliQ water for 1 hour at room temperature within a hydrophobic barrier. Wholemounts were then incubated with primary antibodies in the block/stain buffer at 4 degrees for 16 hours, washed three times for ten minutes each in PBS with 0.2% triton and, if required, incubated with secondary antibodies at room temperature for two hours. Duras were mounted with Fluoromount-G™ Mounting Medium.

### Confocal Microscopy and Image Analysis

Tile scan images of whole-mount meninges were acquired on the Stellaris SP8 laser-scanning confocal microscope equipped with four detectors, six laser lines (405, 458, 488, 515, 559 and 635 nm) and five objectives (×4/0.16 NA, ×10/0.4 NA, ×20/0.75 NA and ×40/0.95 NA, and chromatic-aberration-corrected ×60/1.4 NA) or a STELLARIS 8 confocal microscope with a white-light laser, a 405 nm diode laser, five HyD detectors and four objectives (×5/0.15 NA, ×20/0.75 NA, ×40/1.3 NA and ×60/1.4 NA) (Leica Microsystems). Images from confocal tile scans were imported into Imaris v.9.5 (Bitplane). To differentiate cell subsets in the peri-sinus region, including the superior sagittal and transverse sinuses, and lobe regions, a mask was manually drawn using the surface function, and the surface area in μm^2^ was determined by exporting surface statistics. Absolute cell numbers were quantified within surfaces by identifying and thresholding on positively stained cells within the three-dimensional surface of each respective channel using the spot-detection function. Resultant statistics were then exported into Excel (Microsoft) for further analysis.

For histocytometry plot generation, a surface was created for the entire meningeal whole mount, creating a value-based visual surface for all positively stained cells in each channel, allowing for quantification of fluorescence intensity and the frequency of labelled and unlabelled cells, as well as their position in an *xy* plane. Channel statistics were then exported into Excel (Microsoft) and plotted in FlowJo software using the text to FCS conversion function (TreeStar).

### Sample processing for single cell RNA-sequencing

Age-matched 6-8 week-old EP*^ZFTA-RELA^* and Nestin*^CreERT2^* mice were intravenously injected with CD45- PE 3 minutes prior to schedule 1. For blood, a single eye was removed using fine curved forceps, rupturing the retro-orbital sinus. Three drops of blood were collected into 1 mL of PBS with 0.025% heparin to prevent coagulation. Samples were kept on ice, for the entirety of collection. Blood was centrifuged at 400 x g for five minutes, and red blood cell (RBC) lysis was performed by resuspension in 1 mL of ACK lysis buffer(Quality Biological) for one minute, then 2 mL of ice-cold PBS was added, samples were centrifuged, and lysed red blood cells were aspirated from the leucocyte-containing pellet. The pellet was resuspended in fluorescence activated cell sorting (FACS) buffer (0.1 M, pH 7.4 PBS with 1% BSA and 1 mM EDTA) until use. Meningeal dura was carefully collected under a dissection microscope. Meninges and calvaria were then digested for 15 minutes at 37°C with constant agitation using 1 mL of pre-warmed digestion buffer (DMEM, with 2% FBS, 1 mg/mL collagenase D (Sigma

Aldrich), and 0.5 mg/mL DNase I (Sigma Aldrich)), filtered through a 70 μm cell strainer, and enzymes neutralized with 1 mL of complete medium (DMEM with 10% FBS). An additional 2 mL of FACS buffer was added, samples were centrifuged at 400 x g for five minutes, and samples were resuspended in FACS buffer and kept on ice until use. For peripheral bone marrow, both tibia were flushed with 0.05% BSA PBS with 0.05% EDTA, filtered through 100 µm meshes and washed with 2% fetal bovine serum in RPMI and resuspended on 0.05% BSA PBS solution. The whole intact deep cerbical lymph nodes were mashed through a 70 μm cell strainer, using a sterile syringe plunger, and washed with 5 mL of FACS buffer. Deep cervical lymph node samples were then filtered through 100 µm meshes and washed with 2% fetal bovine serum in RPMI and resuspended on 0.05% BSA PBS solution. Brains were macrodissected based on TdTomato fluorescent signal to harvest the tumour and region-matched brain in control bearing animals. Tumour/brain samples were macrodissected, removing choroid. Plexus from the third, fourth and lateral ventricles and were mechanically dissociated using sterile surgical scalpels into ∼1 mm^3^ cubes, and dissociated using the mouse tumour dissociation kit (Miltenyi Biotec) using the gentleMACS Octo Dissociator (Miltenyi Biotec). After dissociation, the samples were then filtered through 100 µm meshes and washed with 2% fetal bovine serum in RPMI, spun down 420 g for 5 minutes. Samples were resuspended in 40% percoll and centrifuged at 600 x g for 10 minutes. Supernatant was removed and washed with 2% fetal bovine serum in RPMI and resuspended on 0.05% BSA PBS solution. Samples were stained with DAPI (0.2 µg/ml). Samples were centrifuged, resuspended in FACS buffer with anti-CD16/32 (FC block; Biolegend) diluted 1:50 in FACS buffer and fluorescently conjugated antibodies (anti-CD45 APC and anti-Ter 119 FITC) at 4°C, followed by for 20 minutes. Cells were sorted using the Influx Cell Sorter (BD Biosciences) or FACsAria II (BD Biosciences) into 1% BSA coated 1.5 mL Eppendorf tubes with 500 μL of DMEM.

### Sample processing for single nuclei RNA-sequencing

Single cell suspension of skull and tibial bone marrow cells were obtained as described above. Cells were sorted for Live CD45 i.v^-^Ter119^-^CD45^+^Lin^-^CD34^+^ using the Influx Cell Sorter (BD Biosciences) or FACsAria II (BD Biosciences) into 1% BSA coated 1.5 mL Eppendorf tubes with 500 μL of DMEM.

FACS-isolated cells were centrifuged at 500 x g for 5 minutes at 4 degrees Celsius before removing the supernatant and resuspending in 500 uL lysis buffer (10 mM Tris-HCl, 3 mM MgCl₂, 2 mM NaCl, 0.005% NP-40 substitute, 0.1 mM DTT, SUPERase RNase inhibitor 0.25 U/mL and protease inhibitor (A32965) for 2 minutes. Nuclei were pelleted at 500 × g for 5 min at 4 °C, washed in PBS with 1% BSA, and counted using Trypan Blue exclusion. Single-nucleus multiome libraries (RNA + ATAC) were prepared using the Chromium Next GEM Single Cell Multiome ATAC + Gene Expression kit (10x Genomics) according to the manufacturer’s protocol. Libraries were sequenced on an Illumina NovaSeq 6000.

### Sample processing of human neurosurgical tissue

To obtain flow cytometry of skull, dura and tumour shown in Supplementary Fig. 4, discard neurosurgical material from skull fragments, dura and tumour were collected from a 6 month old male patient undergoing routine tumour debulking for an atypical choroid plexus papilloma (WHO Grade 2). Skull fragments and dura were obtained during craniotomy into RPMI-1640 supplemented with 10% FBS and kept on ice. Larger bone fragments were roughly crushed using a sterile scalpel. The suspensions was centrifuged at 600 × g for 10 min at 4 °C, and the pellet was resuspended in 10–15 mL RPMI containing L-glutamine, 2 mg/mL Collagenase D, 2 mg/mL Collagenase A, and 0.5 mg/mL DNase I. Samples were digested for 1 h at 37 °C with rotation using a MACS rotator. Digested material was filtered through a 70 μm cell strainer, and enzymes were neutralized with 10 mL RPMI + 10% FBS. Cells were pelleted at 600 × g for 10 min at 4 °C. Debris was removed using MACS debris removal solution (130-109-398) prior to flow cytometry.

Freshly resected human brain tumor tissue was placed in a sterile Petri dish, and cauterized regions were removed with a scalpel. Remaining viable tissue was cut into ∼3–8 mm³ pieces, weighed, and transferred to a 50 mL conical tube. The dish was rinsed thoroughly with HBSS, and the volume was adjusted to 20 mL. Tissue was pelleted at 300 × g for 10 min at room temperature, and the supernatant was aspirated. Samples were resuspended in enzyme-containing dissociation buffer and transferred to C-tubes (Miltenyi Biotec) following the manufacturer’s protocol, with enzyme volumes adjusted to tissue weight. Mechanical and enzymatic dissociation was performed using the gentleMACS Octo Dissociator with Heaters (program: *37C_BTDK_1*). Cell suspensions were filtered through a 40 μm strainer into fresh 50 mL tubes and rinsed with ∼20 mL HBSS to maximize yield. Cells were pelleted at 300 × g for 10 min at 4 °C, leaving 1 mL residual volume. The pellet was resuspended in 7 mL of 40% Percoll (4.32 mL Percoll, 0.48 mL 10× PBS, 7.2 mL RPMI) and centrifuged at 600 × g for 10 min at 4 °C with no brake. The myelin-rich top layer was discarded, and the cell pellet was resuspended in 500 μL PBS, then transferred to a new tube containing 5 mL PBS and processed as per below for flow cytometry. The samples were obtained from Cambridge University Hospital, REC reference 23/EE/024.

### Flow cytometry

Cell suspensions were prepared as described above and transferred into a V-bottom plate. Viability staining was performed using Zombie NIR (1:500 in PBS, 10 minutes, room temperature; BioLegend). Suspensions were then pelleted (450*g* for 5 minutes) and resuspended in anti-CD16/32 antibody (1:100, BioLegend) diluted in FACS buffer to block Fc receptor binding. Antibodies against cell surface epitopes were then added for 10 minutes at room temperature. For a full list of antibodies, see **Supplementary Table 3**. Flow cytometry was performed using an Aurora spectral flow cytometer (Cytek Biosciences), and data were analyzed with FlowJo (version 10, BD Biosciences).

### CSF collection and intra-cisterna magma injections

Mice were anesthetized via intraperitoneal injection of ketamine (100 mg kg^−1^) and xylazine (10 mg kg^−1^) in saline and placed on a stereotactic frame. The fur over the incision site was clipped, and the skin was disinfected with three alternating washes of alcohol and Betadine. For intracerebral injections, a midline incision was made along the scalp, exposing the dorsal skull. A burr hole was carefully made using a dental drill. A 1:1 ratio of OVA-594 and anti-c-Kit-PE (1 µl) was injected using a glass capillary attached to a microinjector (World Precision Instruments) over 2 minutes at the following coordinates: +1.5 A/P, −1.5 M/L, −2.5 D/V. The glass capillary was left in place for another 2 minutes to prevent backflow. For ICM injections, the posterior scalp and neck was shaved and prepared with iodine antiseptic. The head was placed in a stereotactic frame with the neck flexed. A midline incision was made and the posterior nuchal musculature divided, exposing the inferior, dorsal aspect of the occipital bone and the posterior dura overlying the cisterna magna. A glass capillary attached to a microinjector (World Precision Instruments) was used. 5 ul was infused and injection rates were adjusted to achieve a 5-minute injection, followed by a 5-minute wait period to prevent backflow. For CSF collection, a glass capillary was inserted through the dorsal dura mater into the superficial cisterna magma and approximately 50 µL of CSF was drawn by capillary action.

For CSF transfer experiments 10 µl of CSF was transferred. Mice were allowed to recover on a heating pad. For animals undergoing survival surgery, the skin was sutured and ketoprofen was subcutaneously injected for postoperative analgesia. 6 hours after transfer, mice were sacrificed with an overdose of anaesthesia, CSF was obtained as described above, and mice were necropsied, including the careful removal of the leptomeninges and choroid plexus from the 3^rd^, 4^th^ and lateral ventricles.

### CSF collection and multiplex analyte analysis

Mice were anesthetized via intraperitoneal administration of ketamine/xylazine and placed on a stereotactic frame. CSF was collected from the cisterna magna with a 30 gauge needle. CSF (12.5 μl) was obtained from each mouse, and analyte quantification was performed using Luminex magnetic beads with the Bio-Plex Pro Mouse Cytokine Panel 23-plex instruction (Bio-Rad). Data were acquired with the Luminex FLEXMAP 3D and analyzed with xPONENT software version 4.2 (Luminex).

### Sample dissolution, TMT labelling and Reverse-Phase fractionation

A 50µl aliquot of CSF was mixed with lysis buffer containing 100mM Triethylammonium bicarbonate (TEAB, Sigma), 10% Isopropanol, 50mM NaCl, 1% SDC with Nuclease and Protease/Phosphatase Inhibitors and incubated at RT for 15 min followed by water bath sonication. Later the samples were reduced and alkylated with 2ul of 50mM tris-2-caraboxymethyl phosphine (TCEP, Sigma) and 1ul of 200mM Iodoacetamide (IAA, Sigma) respectively for 1 hour. Then digested overnight at 37°C using trypsin solution at ratio protein/trypsin ∼ 1:30. The next day, protein digest was dried and suspended in 0.1M TEAB was labelled with the TMTpro reagents (Thermo Scientific) for 1 hour. 1µl of each labeled sample was taken and analysed as pre-mix to normalize the sample volume across all the conditions. Later the reaction was quenched with 4 μL of 5% hydroxylamine (Thermo Scientific) for 15 min at room temperature (RT). All the normalized sample volume were mixed and acidified using Formic Acid for SDC removal. Samples were fractionated with Reversed-Phase cartridges at high pH (Pierce #84868). Nine fractions were collected using different elution solutions in the range of 5–50% ACN and dried completely by vacuum centrifugation. Each fraction was reconstituted in 0.1% formic acid for liquid chromatography tandem mass spectrometry (LC–MS/MS) analysis.

### LC-MS/MS

Peptide fractions were analysed on a Vanquish Neo UHPLC system coupled with the Orbitrap Ascend (Thermo Scientific) mass spectrometer. Peptides were trapped on a 100μm ID X 2 cm microcapillary C18 column (5µm, 100A) followed by 90 minutes elution using 75μm ID X 25 cm C18 RP column (3µm, 100A) at 300nl/min flow rate. A Real Time Search (RTS)-MS3 method was used for the analysis; The MS1 spectra were acquired in the Orbitrap (R=120K; scan range 400-1600 m/z; AGC target = 400000; maximum IT = 251ms) and the MS2 spectra in the Ion Trap (isolation window 0.7Th; collision energy of 30%; maximum IT = 35ms; centroid data). For RTS, Trypsin/P digestion was selected using static cysteine carbamidomethylation and TMTpro modification on lysine and peptide N-terminus. The search was conducted for a maximum of 35ms with the following thresholds: Xcorr =1.4, dCn = 0.1, precursor ppm 10, charge state = 2. MS3 spectra were collected in the Orbitrap (R= 45K; scan range 100−500m/z; normalized AGC target = 200%; maximum IT = 200ms; centroid data. Phospho fractions were subjected to MS2 analysis without RTS.

### Data processing

The Proteome Discoverer 3.0. (Thermo Scientific) was used for the processing of CID tandem mass spectra. The SequestHT search engine was used, and all the spectra searched against the Uniprot Mouse FASTA database (taxon ID 10090 - Version October 2024). All searches were performed using a static modification of TMTpro (+304.207 Da) at any N-terminus and at lysine and Carbamidomethyl at Cysteines (+57.021 Da). Methionine oxidation (+15.9949Da) and Deamidation (+0.984) on Asparagine were included as dynamic modifications. Mass spectra were searched using precursor ion tolerance 20 ppm and fragment ion tolerance 0.5 Da. For peptide confidence, 1% FDR was applied, and peptides uniquely matched to a protein were used for quantification.

### Single-cell library construction and sequencing

For scRNA-seq, gel bead in emulsions were prepared by loading up to 10,000 cells per sample onto the Chromium Chip G (10x Genomics 1000073) and run using the Chromium Controller (10x Genomics). cDNA libraries were generated using the Chromium Single Cell 5′ GEM, Library and Gel Bead Kit V3 (10x Genomics) with V(D)J. Libraries were sequenced using the Novaseq600 Kit v2.5 (Illumina) on the Illumina Novaseq600 system.

### scRNA-seq analysis of mouse samples

Single-cell gene expression data from Cell Ranger (v.7) output was analysed using standard scanpy (v.1.8.2) workflow. Doublet detection was performed using scrublet (v.0.2.3) with adaptations outlined previously. In brief, after scrublet was performed, the data were iteratively subclustered, and a median scrublet score for each subcluster was computed. Median absolute deviation scores were computed from the cluster scrublet scores and a one-tailed *t*-test was performed with Benjamini–Hochberg correction applied and cells with significantly outlying cluster scrublet scores (Benjamini–Hochberg-corrected *P* < 0.1) were flagged as potential doublets. Quality control and filtering were performed in Scanpy using the sc-dandelion pre-processing module (dandelion.pp,recipe_scanpy_qc) with max. genes = 6000, min. genes = 200 and a Gaussian mixture model distribution to determine mitochondrial filtering value .Genes were retained if they were expressed by at least three cells. Gene counts for each cell were normalized to contain a total count equal to 10,000 counts per cell. This led to a working dataset of 125,000 cells. Highly variable genes were selected based on the following parameters: minimum and maximum mean expression are ≥0.0125 and ≤3 respectively; minimum dispersion of genes = 0.5. Batch correction was performed using harmonypy (v.0.0.5) after principal component analysis. Clustering was performed using the Leiden algorithm (v.0.8.2) with the resolution set at 1.0. The size of local neighbourhoods was set at 10. Uniform manifold approximation and projection (UMAP; v.0.5.1) was used for dimensional reduction and visualization, the minimum distance was set at 0.3 and all other parameters as per default settings in scanpy. Cell type identification was guided by manual inspection of canonical marker gene expression and top marker genes identified with Wilcoxon rank-sum tests.

For analysis of differentially expressed genes between conditions, cells or tissues were subset based on sub-cluster identity and gene expression and each subset was filtered to include genes that had at least 4 transcripts in at least 4 cells. Then the top 2000 highly variable genes were determined and included for further analysis using the SingleCellExperiment (v.3.20) modelGeneVar (scran, v1.35.0) and getTopHVGs (scran, v.1.35.0) functions. After filtering, limma (v.3.2.0) and edgeR (v.4.0) were used to build a model and conduct differential expression testing with the lmFit, contrasts.fit, and eBayes functions. Results were then filtered using a Benjamini-Hochberg adjusted p-value threshold of less than 0.05 as statistically significant. Over representation enrichment analysis with Fisher’s Exact test was used to determine significantly enriched Gene Ontology (GO) terms (adj. p < 0.05) for the sets of significantly differentially expressed genes. For each gene set, genes were separated into up- and downregulated and separately, the enrichGO function from the clusterProfiler (v.3.2.0) package was used with a gene set size set between 10 and 500 genes and p-values adjusted using the Benjamini-Hochberg correction

The list of proteins identified in the CSF LC-MS/MS (Supplementary Table 16) was converted to coding genes with the use of biomaRt (v.3.20) using the UniProt ID and Ensembl gene name as conversion factors. This list was then filtered to include only genes contained in the list of ligands in the annotated reference provided by RNAMagnet (v.0.1.0) with the function getLigandsReceptors with the cellularCompartment parameter set to ‘secreted’, ‘ECM’ or ‘both’ and the version set to 3.0.0. The receptors matching those ligands were mined from the reference (Supplementary Table 17), and their expression was plotted for cell types of interest as average normalized mRNA transcripts per population with the circlize (v.0.4.16) package in R.

To compare cytokine signalling activity between tumour and control tissues, we applied CytoSig to our scRNA-seq data^56^. Normalized gene expression matrices were generated following standard preprocessing (log-normalization, scaling, and batch correction) and used as input for CytoSig (https://cytosig.jhubiostatistics.org) using default parameters. Cytokine activity scores were inferred for each cell based on curated transcriptional signatures of cytokine responses derived from perturbation experiments. Scores were aggregated across cell types and tissues to identify differential cytokine usage between tumour and matched control environments.

Raw sequencing data were processed with Cell Ranger (v7.1.0) to generate V(D)J contigs. Annotated clonotype data were further analysed using Dandelion (v0.1.5)^57^. TCR α and β chains were matched per cell, and only high-confidence, productive clonotypes were retained. Clonotype expansion, overlap, and diversity metrics were computed using Dandelion’s built-in functions. Clonotype integration with transcriptomic data was performed by mapping TCR metadata onto the corresponding cell barcodes from RNA-seq datasets.

### scRNA-seq integration and analysis of human data

Single-cell gene expression data from Cell Ranger (v7.0) were processed with Scanpy (v1.8.2). Doublets were identified using Scrublet (v0.2.3) with adaptations as described previously: data were iteratively subclustered, median Scrublet scores were computed per subcluster, and clusters with significantly outlying scores (Benjamini–Hochberg–corrected P < 0.1, one-tailed t-test) were flagged. Quality control was performed with sc-dandelion (dandelion.pp.recipe_scanpy_qc; max. genes = 6000, min. genes = 200), with mitochondrial thresholds determined by Gaussian mixture modelling. Genes expressed in <3 cells were removed. Counts were normalized to 10,000 per cell, yielding 125,000 high-quality cells for analysis.

Highly variable genes were selected (mean expression ≥0.0125 and ≤3; dispersion ≥0.5). Data were log-transformed, scaled, and subjected to PCA. Batch correction was performed with harmony (v0.0.5). Clustering was performed with the Leiden algorithm (v0.8.2; resolution = 1.0, neighbourhood size = 10). Dimensionality reduction and visualization used UMAP (v0.5.1; minimum distance = 0.3, otherwise default parameters). Cell types were annotated based on canonical marker genes and Wilcoxon rank-sum testing of cluster markers.

For cross-dataset integration, we implemented a two-step approach. First, batch and confounder effects were regressed out using a custom Ridge regression model (scikit-learn, v1.3.0). One-hot encoded batch and confounder variables were fitted against the expression matrix, and the predicted contributions were subtracted. Second, BBKNN was applied to principal components from the corrected data to generate a batch-balanced neighborhood graph, with data origin as the variable. UMAP and Leiden clustering were then performed on the BBKNN graph to define shared cell states across scRNA-seq and snRNA-seq datasets.

### snATAC + GEX mutliome analysis

Libraries were sequenced on an Illumina NovaSeq 6000 platform to achieve a minimum of 25,000 unique fragments per nucleus, as recommended by the manufacturer. Raw sequencing data were processed using the 10x Genomics Cell Ranger ARC pipeline (v.2.0), which aligns the ATAC reads to the reference genome (mm10) and identifies barcoded fragments.

The raw data were imported into R (v4.4.0) for downstream analysis using the Signac and Seurat R packages. First, fragment files were processed using Signac to create a fragment object linked to the associated barcodes. Quality control (QC) metrics were calculated, including the total number of fragments per cell, fraction of reads in peaks (FRiP), and nucleosome signal. Low-quality nuclei were excluded based on the following criteria; Less than 1000 fragments, nucleosome signal >2. Doublets were identified and removed using the DoubletFinder (v.2.0.4) package.

Peaks were called on aggregated ATAC-seq data using MACS2 (v2.2.9.1) to define a unified set of reproducible peaks. A peak-by-cell matrix was constructed by quantifying the number of fragments overlapping each peak per nucleus. For dimensionality reduction, Latent Semantic Indexing (LSI) was applied to the binary peak matrix, followed by principal component analysis (PCA) to identify major axes of variation. Uniform manifold approximation and projection (UMAP) was then used to visualize chromatin accessibility profiles in a low-dimensional space.

For datasets containing paired ATAC-seq and gene expression (RNA-seq) profiles, multimodal integration was performed using Seurat (v4.4.0). A shared UMAP embedding was generated by linking RNA and ATAC modalities through anchor-based integration with the FindMultiModalNeighbors function. Joint clustering of the RNA and ATAC datasets was achieved using the FindClusters function, enabling visualization of shared cell states across both modalities in the same low-dimensional space. Differentially accessible regions (DARs) between tumor-bearing and non-tumor-bearing samples were identified using a Wilcoxon rank-sum test, implemented in ArchR’s getMarkerFeatures() function. Motif enrichment analysis was performed using chromVAR (v1.16.0), leveraging cisBP motif annotations to uncover transcription factor activity associated with condition-specific chromatin states. To examine dynamic changes in accessibility, pseudotime trajectories were constructed using ArchR’s addTrajectory() function, allowing for the visualization of progressive changes in chromatin states along differentiation or activation pathways. All analyses were performed in R (v4.3.1), and visualization of results was conducted using ggplot2 (v3.4.3) and patchwork (v1.1.3)

### Quantitative Polymerase Chain Reaction (qPCR)

For qPCR analyses, cells were directly sorted into RNA lysis buffer (Arcturus PicoPure RNA isolation Kit (Invitrogen)), incubated for 30 min at 42°C and processed for cDNA synthesis using SuperScript VI cDNA synthesis kit (Invitrogen) according to manufacturer’s instructions. The newly synthesized cDNA was diluted 1:10 in RNAse free H_2_O and 6 μL were mixed in technical triplicates in 384-well plates with 0.5 μl of forward and reverse primer (10 μM) (Supplementary Table 4) and 7 μl PowerUP SybrGreen Mastermix (ThermoFisher). Program: 50°C for 2 minutes, 95°C for 10 minutes and 40 cycles of 95°C for 15 seconds, 60°C 1 minute. Primers were designed to be intron spanning whenever possible using the Universal ProbeLibrary Assay Design Center (Roche) and purchased from IDT. Experiments were performed on the Quant StudioTM 7 (Applied Biosystems) and analysis of gene amplification curves was performed using the Quant StudioTM Real-Time PCR Software v1.3 (Applied Biosystems). RNA expression was normalized to the housekeepers *Gapdh/Actb* for murine gene expression analysis. Relative expression levels are depicted in 2^-ΔCt^ values, ΔCt = (geoMean Housekeeper Ct) - (gene of interest Ct).

### Bone Marrow Transplantation

To test the stem cell potential of intratumoural MHC-II populations, equal numbers (5x10^4^) of CD45+Lin- MHC-II+ or MHC-II- intratumoural cells were transplanted intravenously into non-myeloablative busulfan (25 mg/kg)-conditioned mice. 16 weeks post-transplantation total bone-marrow cells (1 x 10^6^) were transplanted into secondary recipients. Mice were bled every 4 weeks and cells were stained as described above to assess engraftments.

### *In vitro* expansion of naïve CD4⁺ T cells

Cryopreserved C57BL/6 or 2D2 splenocytes were thawed using standard protocols and washed in FACS buffer (PBS with 2% FBS and 2 mM EDTA). Cells were centrifuged at 300 × *g* for 10 min and resuspended at 10⁷ cells per 80 µL. For T-cell enrichment, cells were incubated with 20 µL of biotinylated anti-mouse pan T-cell antibody (Miltenyi Biotec) for 15 min on ice, followed by 20 µL of anti-biotin microbeads (Miltenyi Biotec) per 10⁷ cells for an additional 15 min on ice with intermittent agitation. Labeled cells were washed and subjected to magnetic separation using MS columns on a magnetic stand. Negative fractions were collected during column loading, and positively selected T cells were eluted by removing the column from the magnet and flushing with buffer using a plunger.

Viable CD4⁺ T cells were counted and cultured in AIM-V media supplemented with 5% heat-inactivated FBS, penicillin-streptomycin, L-glutamine, and recombinant mouse IL-2 (40 IU/mL; R&D Systems). Cells were maintained at a density below 1.5 × 10⁶ cells/mL and stimulated with Dynabeads® Mouse T-Expander CD3/CD28 beads (1 × 10⁸/mL; Invitrogen, Cat# 111.41D) following the manufacturer’s protocol.

### Adoptive transfer and immunisation

To test the ability of endogenous CNS antigens to polarize skull CD4 T cells, expanded naïve CD4⁺Foxp3⁻CD90.1⁺ T cells were adoptively transferred into RAG2⁻/⁻ recipient mice. Mice were immunised intrathecally with artificial cerebrospinal fluid (aCSF), MOG₃₅₋₅₅ peptide, or a MOG-based fusion peptide at days 21 and 42 post-transfer. At day 49, dura and skull bone marrow tissues were harvested and analysed for Foxp3 expression within the transferred CD90.1⁺ T cell population by flow cytometry.

To test the ability of endogenous CNS antigens to polarise antigen-specific CD4 T cells; naïve CD4⁺ T cells (5×10⁴) from 2D2 TCR transgenic mice were expanded in vitro and transferred into wild-type recipient mice. Mice received intrathecal injections of aCSF or MOG₃₅₋₅₅ peptide (2.5 μL of 2 mg/mL) on days 1 and 7. On day 14, dura and skull bone marrow were harvested and analyzed for Foxp3⁺CD4⁺ T cells.

### *In vivo* antigen presentation assays

For analysis of presentation of exogenous antigens on HSPCs, DQ-OVA (100μg/mL, Invitrogen), Eα peptide (52-68) (500μg/mL, Mimotopes) or a control IgG2b antibody (100μg/mL, eB149/10H5, ThermoFisher) were injected intrathecally as described above, and mice were sacrificed after 6 hours.

### Murine *ex vivo* cultures

Cells were cultured at 37°C and 5% CO_2_ in U-bottom plates in a total volume of 200μL of Dulbecco’s Modified Eagle’s Medium GlutaMAX (DMEM GlutaMAX, Gibco) supplemented with 10% heat-inactivated Fetal Calf Serum (FCS, Gibco), sodium pyruvate (1.5mM, Gibco), L-glutamine (2mM, Gibco), L-arginine (1x, Sigma), L-asparagine (1x, Sigma), penicillin/streptomycin (100 U/mL, Sigma), folic acid (14μM, Sigma), MEM non-essential amino acids (1x, ThermoFisher), MEM vitamin solution (1x, ThermoFisher) and β-mercaptoethanol (57.2μM, Sigma). Cells were sorted and, when mentioned, labelled with cell trace violet (ThermoFisher) according to manufacturer’s instructions. 5x10^4^ naïve CD4^+^ T cells were cultured with 2x10^4^ HSPCs, DCs or CD8^+^ T cells, unless stated otherwise. When stated, ovalbumin peptide (323-339) (25μg/mL, Invivogen), DQ-OVA (100μg/mL, Invitrogen), Eα peptide (52-68) (100μg/mL, Mimotopes), αMHC-II blocking antibody (10μg/mL, M5/114.15.2, BioXCell) or a control IgG2b antibody (10μg/mL, eB149/10H5, ThermoFisher) were added to the cultures.

LSK cells were isolated as described above and cultured for 12 hours in presence or absence of ovalbumin peptide (50μg/mL) in culture medium supplemented with TPO (50 ng/mL, PreproTech) and SCF (50 ng/mL, PreproTech) at 37°C, 5% CO2 levels. These cells were then co-cultured with OT-II CD4+ T cells for 72 hours

### Griess assay and ROS production in LSK progeny

To quantify the levels of nitrite and ROS in LSKs, isolated by FACS and cultured for 12 hours in presence or absence of ovalbumin peptide (50μg/mL) in culture medium supplemented with TPO (50 ng/mL, PreproTech) and SCF (50 ng/mL, PreproTech) at 37°C, 5% CO2 levels. These cells were then co-cultured with OT-II CD4+ T cells for 72 hours. Following treatment, cells were washed twice with PBS and incubated with 100 μL of ROS Detection Solution (containing the ROS-specific fluorogenic probe) at 37°C for 30 min in the dark. After incubation, fluorescence intensity was measured at Ex/Em

= 520/605 nm using a microplate reader (Clariostar). Following plate reading, ROS-medium was washed, cells were lysed and Nitrite (NO₂⁻), a stable oxidation product of nitric oxide (NO), was quantified using the Griess Reagent Kit (Abcam, ab234044) following the manufacturer’s protocol .

### IL-10 Enzyme-linked immunosorbent assay (ELISA)

To quantify the secretion of IL-10 in LSKs, isolated by FACS and cultured for 12 hours in presence or absence of ovalbumin peptide (50μg/mL) in culture medium supplemented with TPO (50 ng/mL, PreproTech) and SCF (50 ng/mL, PreproTech) at 37°C, 5% CO2 levels. These cells were then co-cultured with OT-II CD4+ T cells for 72 hours. Interleukin-10 (IL-10) levels in samples were quantified using a sandwich enzyme-linked immunosorbent assay (ELISA) kit (e.g., Abcam Mouse IL-10 ELISA Kit, ab100697), following the manufacturer’s instructions. Optical density (OD) was measured at 450 nm using a microplate reader (Clariostar, BMG Labtech)

### Colony-forming unit-assay

To observe hematopoietic colony-forming unit (CFU) formation, the cell suspension obtained from skull bone marrow was seeded in methylcellulose media: (MethoCult H4230 and MethoCult SF H4236, Stemcell Technologies) according to manufacturer’s protocol. Both media were supplemented with IL-3 (20 ng/mL), IL-6 (20 ng/mL), G-CSF (20 ng/mL), GM-CSF (20 ng/mL), SCF (50 ng/mL) and erythropoietin (3 units/mL). After incubation for 14–16 days at 37 °C with 5 % CO2, the colonies were characterized and scored according to their morphology on a ZEISS AX10 Inverted Microscope (Zeiss).

### Intracalvarial injection

Skull bone marrow delivery of AMD3100 was performed as previously described^58^. Briefly, mice were anesthetized with a ketamine (100 mg/kg)–xylazine (10 mg/kg) mixture. The head of the mouse was shaved, a skin midline incision was made, and skull was exposed. Using an electrical drill, the outer periosteal layer was thinned on top of skull bone marrow near the bregma and lambda on five spots, without damaging the bone marrow. One microliter of 1 mg/ml AMD3100 (ab120718, Abcam) or vehicle was applied on each spot for 5 min. The skin was sutured, and mice were sacrificed 24 hours later.

### EdU and Ki67 staining for proliferation analysis

Mice received two intraperitoneal injections of 10 mg of EdU per kilogram of body weight 24 hours apart and were sacrificed 24 hours after the final injection. After generation of single-cell suspensions for flow cytometry and surface staining as described earlier, fixation, permeabilization, and EdU staining were performed following the manufacturer’s instruction (Click-iT Plus EdU Alexa Fluor 488 Flow Cytometry Assay Kit, C10632, ThermoFisher Scientific). Intracellular staining for Ki67 and additional staining for PE conjugates was performed for 10 min at room temperature after EdU staining.

### Statistics and reproducibility

Statistical methods were not used to recalculate or pre-determine study sizes but were based on similar experiments previously published. Experiments were blinded, where possible, for at least one of the independent experiments. No data were excluded for analysis. For all experiments, animals from different cages were randomly assigned to different experimental groups. All experiments were replicated in at least two independent experiments of at least five mice per group, and all replication was successful. For all representative images shown, images are representative of at least three independent experiments. Statistical tests for each experiment are provided in the respective figure legends. Data distribution was assumed to be normal, but this was not formally tested. In all cases, measurements were taken from distinct samples. Statistical analysis was performed using Prism (version 10.0, GraphPad Software).

To test for differences in cell numbers among tissues subject to studies e.g., OVA uptake, we deployed a linear mixed-effects model to logit transformation of proportions p [i.e. logit outcome = log(p/(1-p))]. This choice was motivated by the presence of proportions near boundaries 0-1, which is a risk for the normality assumption. The following approaches were used to confirm the results: a) natural scale (i.e. proportions as outcome); b) outliers’ rejection; c) rank transformation; d) generalised linear mixed-effects models with beta distribution as residuals’ distribution; e) distribution-free methods (i.e. Wilcoxon rank-sum test).

Where linear and generalised linear mixed-effects models were used, variability was considered different between strata (i.e. random error, is clearly different between Tibia, Skull and Dura strata). To control the familywise error rate a Bonferroni correction was performed on the two outcomes (percentage of C-Kit^+^ LSK cells and percentage of OVA^+^ macrophage cells). As such, the type I error (false positive result) is controlled at the 5% nominal level. For each outcome I used Wald z-tests on contrasts. Because correlation between different test statistics on the same regression model have been considered, hypotheses testing was more efficient.

### Data availability

The accession number for the Fastq files and quantified gene counts for single-cell sequencing reported in this paper is GEO: GSE28237, GSE82459, GSE300889 and GSE300890. All data are available in the main text or the Supplementary Information files. Data were also sourced from the following published accessions: NEMO; dat-oii74w and dat-3ah9h9x, GEO; GSE141460, GSE126025, GSE156053, GSE226961 and GSE231860. The mass spectrometry proteomics data have been deposited to the ProteomeXchange Consortium via the PRIDE partner repository with the dataset identifier PXD058239.

## Acknowledgments

This work was supported by grants to: R.J.G Cancer Research UK (CRUK) Centre, CRUK Children’s Brain Tumour Centre of Excellence, CRUK Cambridge Institute Core Award; the Brain Tumour Charity Quest for Cures and EC Little Princess Trust; the Brain Tumour Charity Expanding Theories Award. We thank all members of the Gilbertson lab for their continued discussion to push the current study forward. We would like to acknowledge the microscopy core within the Cancer Research UK Cambridge Institute for their support in the acquisition and analysis of meningeal wholemounts, Flow cytometry Core for sorting the numerous cell types for various experiments in this project, Proteomics and Genomics and bioinformatics core for performing the single-cell RNA sequencing and pre- processing, and proteomic analysis. We further extend our thanks to the genome editing core for the generation of the EP*^ZFTA-RELA^* mouse model, specifically Dr Alasdair Russel, Dr Claire King generated the targeting construct. Dr Xiangang Zou and Panagiotis Papadopolous targeted the ES cells and injected these to generate founder mice. We extend our gratitude to the CRUK Cambridge Institute Biological Resource Unit for their immense support with the mouse work. And finally, we extend our gratitude to the patients, donors and families that made this work possible.

## Author contributions

EC conducted the bulk of the experimental procedures. DAP, LH, JT, KWR, JE, OB, GN, LP, KEM, JMV, V.N.R.F and C.S.D conducted/advised on important experimental procedures. SB, JK, RM, CC, IJ, TS, LW, ARK, FJS, TYFH and MRC provided important data and reagents. RJG conceived the research and with EC designed the approach and oversaw the research. All authors contributed to the writing of the manuscript.

## EXTENDED DATA FIGURE LEGENDS

**Extended Data Fig. 1.**
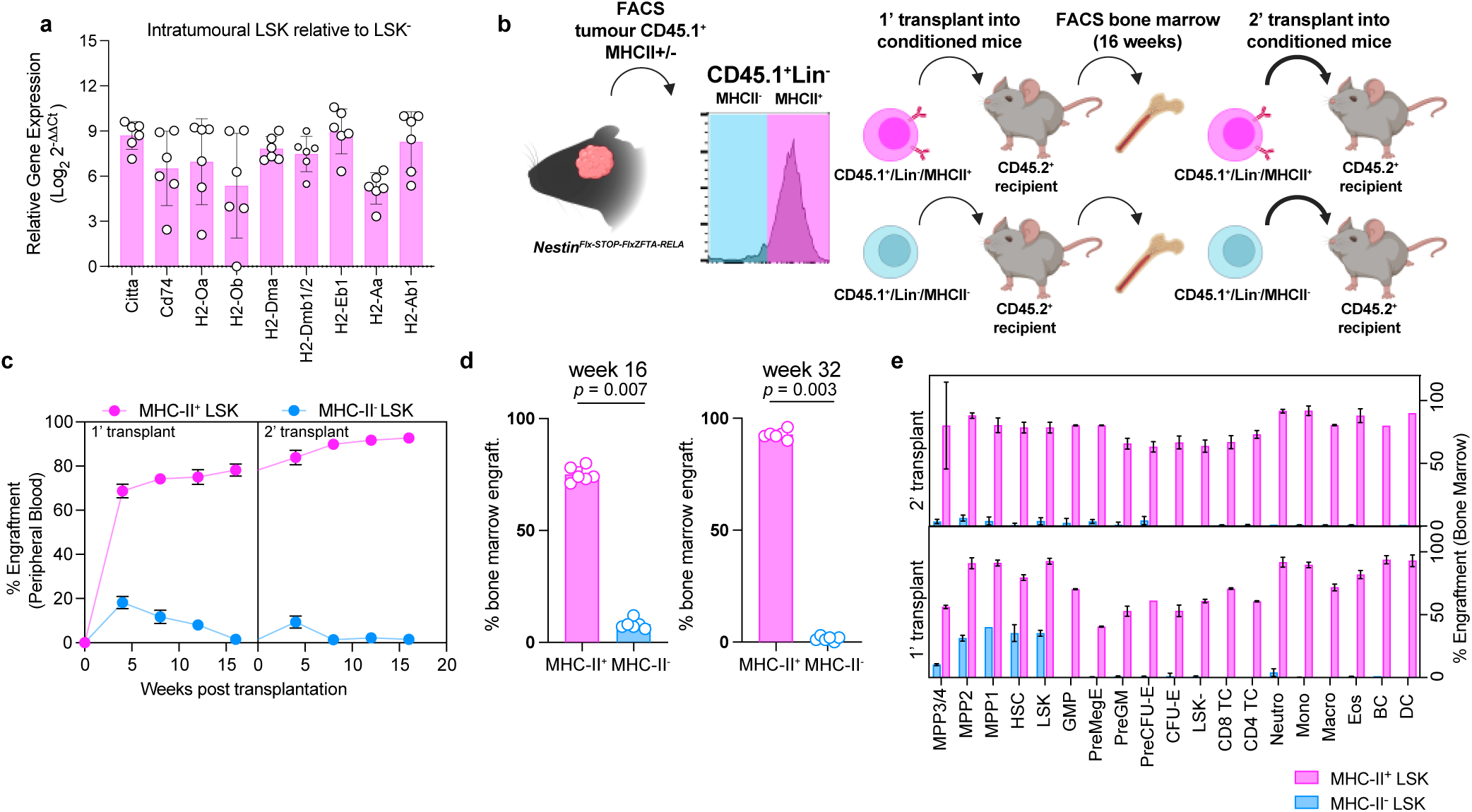
Intratumoural pluripotent haematopoietic stem progenitor cells possess antigen-presentation machinery. **a**, Messenger ribonucleic acid (mRNA) transcript expression of genes encoding antigen presentation molecules and machinery in fluorescent activated cell-sorting (FACS)-isolated LSK^+^ cells, isolated from EP^ZFTA-RELA^ tumours, compared to LSK^-^ cells (n = 7, 10 pooled mice per replicate). **b**, Experimental design pertaining to Figures **b-e,** image created in biorender.com. **c**, Peripheral blood engraftment of MHC-II Lin- cells transplanted from tumors over 16 weeks (n=6, 6-8 pooled mice per replicate) **d**, Quantification of bone marrow engraftment of MHC-II^+^ and MHC-II- tumor-derived Lin- cells (n=6/group, mean±s.e.m., unpaired two-tailed t-test). **e**, Bone marrow engraftment of MHC-II^+^ Lin- cells in different cell types after primary and secondary transplantation; multipotent progenitor (MPP)3/4, MPP2, MPP1, haematopoietic stem cell (HSC), granulocyte-monocyte precursor (GMP) pre-megakaryocyte/erythrocyte (PreMegE), pre- granulocyte monocyte (preGM), pre-colony-forming unit-erythroid cell (CFU-E), CFU-E, LSK-, CD8 T cell (TC), CD4 TC, neutrophil (neutro.), monocyte (mono.), macrophage (macro.) eosinophil (Eos), B cell (BC) and dendritic cell (DC).

**Extended Data Fig. 2.**
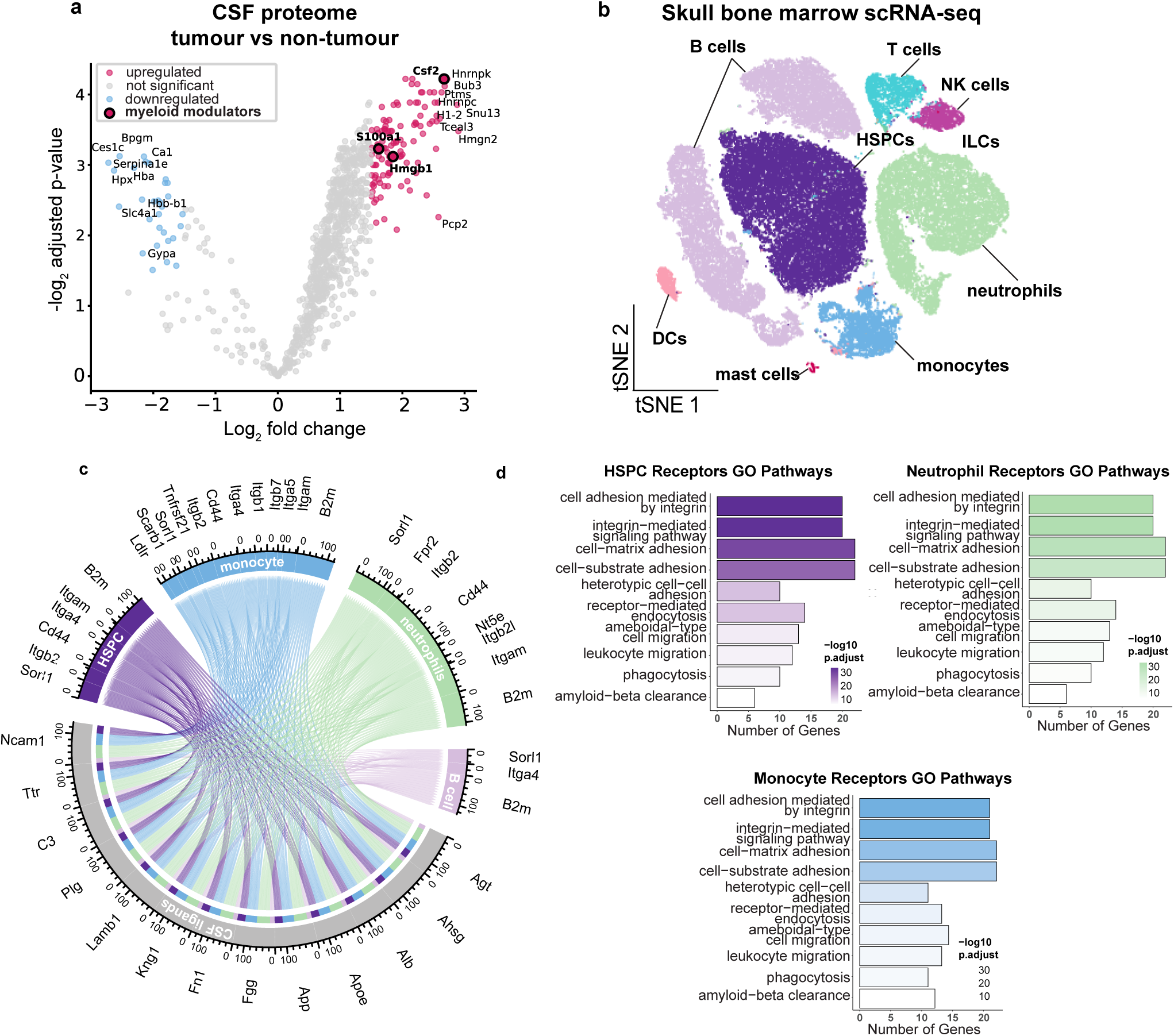
Cerebrospinal fluid (CSF) interacts with the skull bone marrow in EP*^ZFTA-RELA^*-bearing mice. **a**, Volcano-plot of differential expression analysis of CSF proteins in *EP^ZFTA-RELA^*- bearing relative to control bearing mice (n = 6/group). **b**, t-distributed stochastic neighbour embedding (t-SNE) visualizations of scRNA-seq from dorsal skull and tibial bone marrow from 2-month-old mice coloured by cell type; B cells, T cells, natural killer (NK) cells, haematopoietic stem progenitor cells (HSPCs), innate lymphoid cells (ILCs), neutrophils, monocytes, mast cels and dendritic cells (DCs). **c**, Chord plot detailing between CSF ligands identified by tandem mass tag (TMT) Liquid chromatography–mass spectrometry (LC–MS) and receptors on skull bone marrow HSPCs, B cells, monocytes and neutrophils identified by scRNA-seq. **d**, Gene Ontology (GO) pathway analysis on receptor genes with at least one CSF ligand in HSPCs, neutrophils and monocytes.

**Extended Data Fig. 3.**
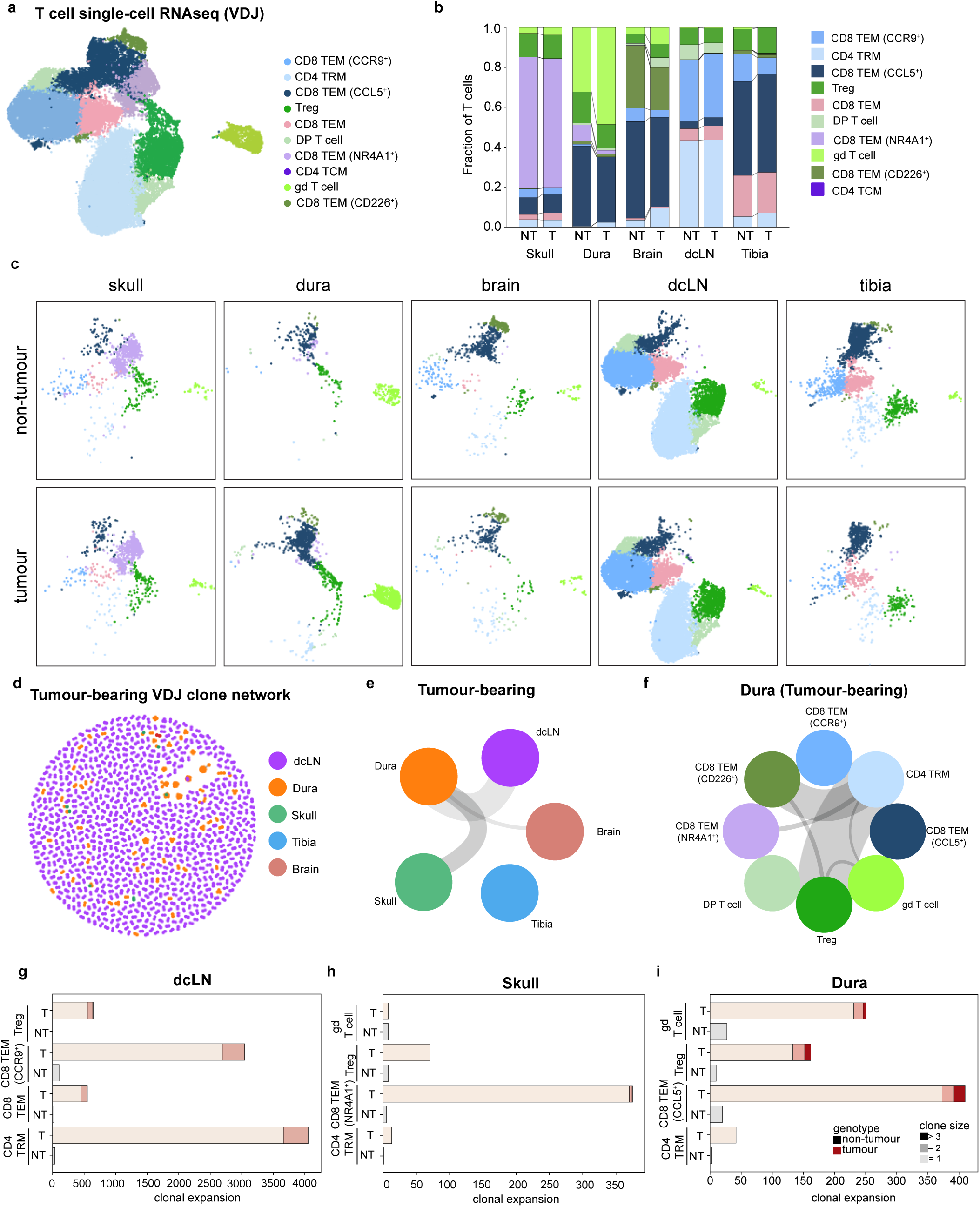
Clonal expansion of T cells in the dura and skull bone marrow of EP*^ZFTA-RELA^*-bearing mice. **a**, uniform manifold approximation and projection (UMAP) visualisation of single- cell RNAseq of T cell subsets from extravascular CD45^+^ cells isolated from the skull and tibia bone marrow, dura, deep cervical lymph nodes (dcLN) and brain/tumour from EP*^ZFTA-RELA^*-bearing and control bearing mice, coloured by cell type; CD8 tissue effector memory (CD8 TEM) CCR9+,, CD4 tissue resident memory (CD4 TRM), CD8 TEM CCL5+, regulatory T cell (Treg), CD8 TEM, douple-positive (DP) T cell, CD8 TEM (NR4A1+), CD4 tissue central memory (TCM), gamma-delta (gd) T cells, CD8 TEM CD266+). **b**, Stacked barplot of T cell populations across tissues in EP*^ZFTA-RELA^*-bearing and control bearing mice. **c**, UMAP of single-cell RNAseq of T cell subsets split by tissue and genotype. **d**, variable– diversity–joining rearrangement (VDJ) T cell receptor (TCR) clone network across tissues in EP*^ZFTA-RELA^*-bearing mice. **e**, circle plot of shared T cell receptor clones between tissues in EP*^ZFTA-RELA^*-bearing mice. **f**, circle plot of shared T cell receptor clones between cell types in dura of EP*^ZFTA-RELA^*-bearing mice. **g-i**, Bar plot of expanded clones in top 4 expanded T cell populations in each tissue.

**Extended Data Fig. 4.**
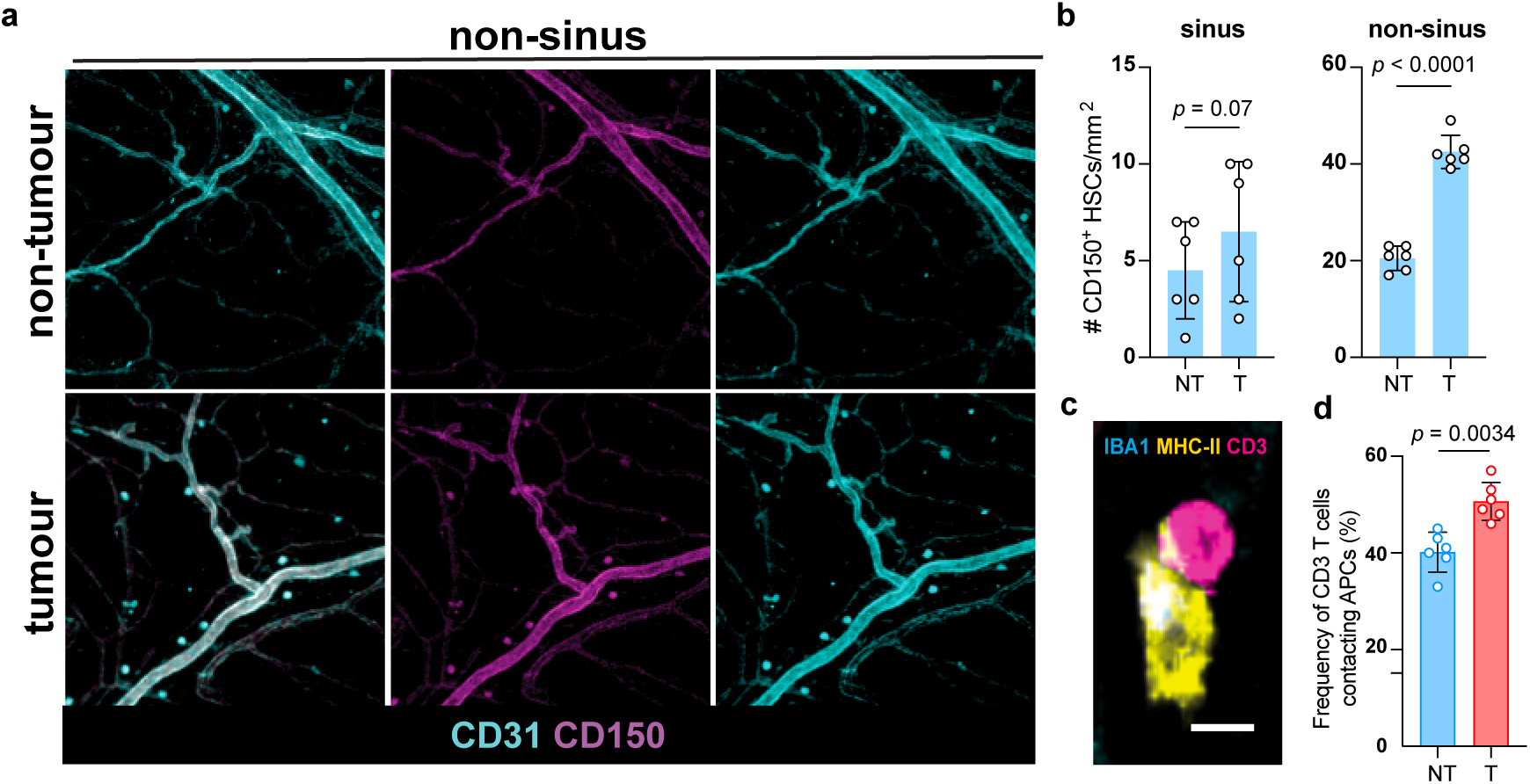
**Elevated numbers of haematopoietic stem progenitor cells located perivascular non-sinus regions in the dura mater. a-b**, Immunohistochemistry and quantification of CD150^+^ HSPCs at non-sinus and sinus regions of the dural meninges in EP*^ZFTA-RELA^*-bearing and control bearing mice (n=6/group, mean±s.e.m, unpaired two-tailed Student’s t-test). **c-d**, Quantification of CD3^+^ T cells contacting MHC II^+^ cells at dural sinuses, n = 6 mice, n = 382 and n = 460 T cell-MHC II interactions counted total (n=6/group, mean±s.e.m, unpaired two-tailed Student’s t-test).

**Extended Data Fig. 5.**
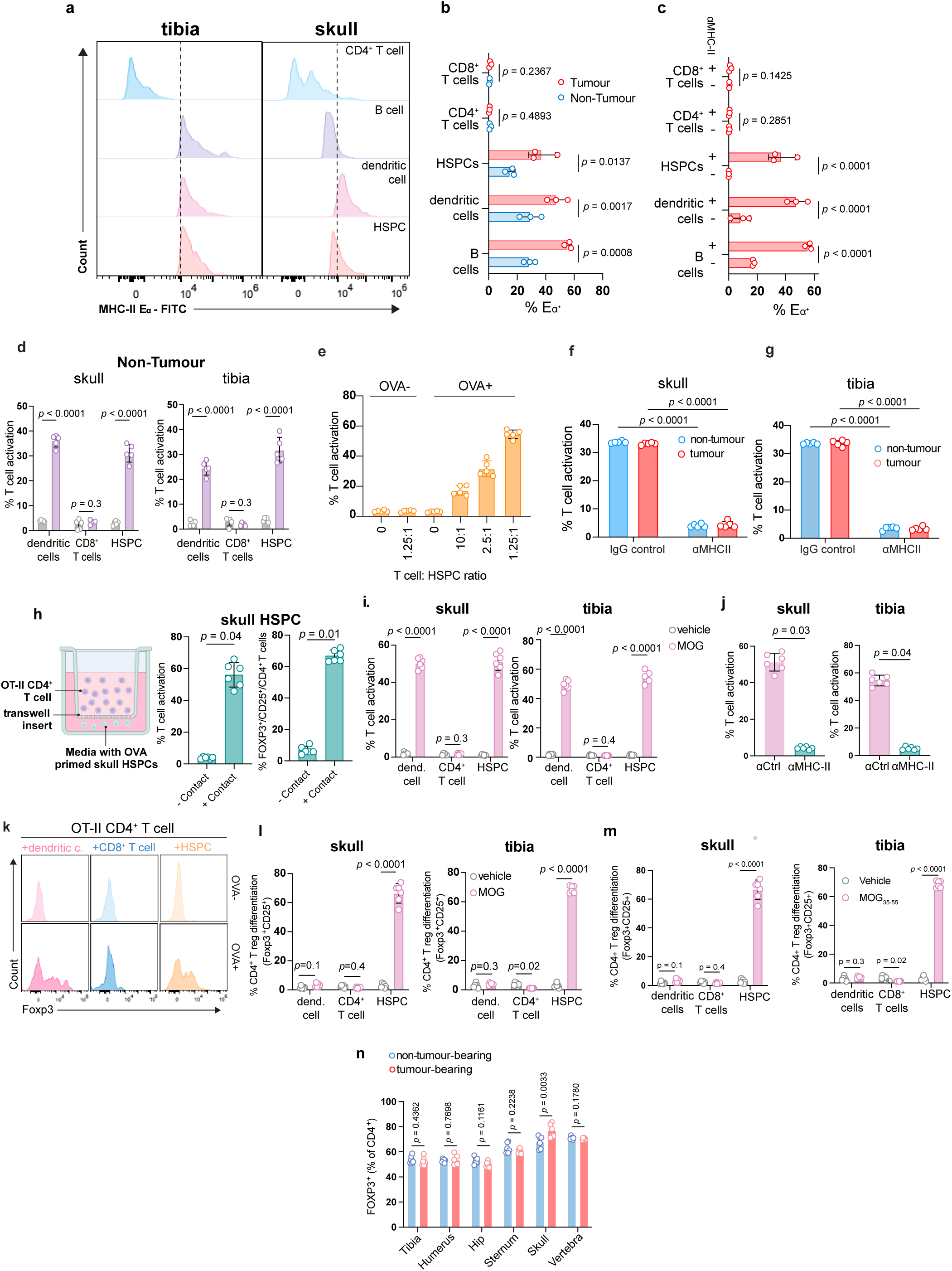
Skull-derived haematopoietic stem progenitor cells polarise CD4 T cells towards Tregs in an antigen-specific manner. **a**, Representative flow cytometry plots of fluorescence- activated cell sorting (FACS) isolated CD4^+^ T cells, B cells, dendritic cells and haematopoietic stem progenitor cells (HSPCs) from the tibia and skull of tumour and control bearing EP*^ZFTA-RELA^* mice pulsed with Eα peptide *ex vivo*. **b**, Quantification of Y-Ae-bound cells from each population as measured by flow cytometry in the presence (**b**) or absence (**c**) of pre-incubation with anti-MHC-II. **d**, T-cell activation (CD44^+^ cells) after co-culture of skull/tibia-derived DCs, CD8^+^ T cells, and HSPCs from control bearing mice with OT-II CD4^+^ T cells (n=5, 10 pooled mice/replicate; mean±s.e.m.; one-way ANOVA with Šídák’s test). **e**, T-cell activation across skull HSPC:OT-II CD4^+^ ratios. **f-g**, T-cell activation in HSPCs ± anti-MHC-II prior to co-incubation with OT-II CD4^+^ T cells (n=5; mean±s.e.m.). **h**, Contact-dependent assessment of OT-II CD4 T cell activation and Treg polarisation with OVA- primed HSPCs (n=5/group, mean±s.e.m, unpaired two-tailed Student’s t-test). **i**, T-cell activation (CD44^+^ cells) after co-culture of skull/tibia-derived DCs, CD8^+^ T cells, and HSPCs from control bearing mice with 2D2 CD4+ T cells (n=5, 10 pooled mice/replicate; mean±s.e.m.; one-way ANOVA with Šídák’s test). **j** T-cell activation in HSPCs ± anti-MHC-II prior to co-incubation with 2D2 CD4^+^ T cells (n=5; mean±s.e.m.). **k-m**, Treg polarisation (FOXP3^+^ CD25^+^ CD4^+^ T cells) after co-culture of skull/tibia-derived DCs, CD8^+^ T cells, and HSPCs pulsed *ex vivo* with MOG_35-55_ with 2D2 CD4^+^ T cells (n=5, 10 pooled mice/replicate; mean±s.e.m.; one-way ANOVA with Šídák’s test). **n**, Quantification of the proportion of FOXP3^+^ CD4^+^ T cells across bone marrow compartments in EP*^ZFTA-RELA^*-bearing and control bearing mice (n = 6 per group, 2 independent experiments), # denotes comparison of non-CNS bone marrow to skull bone marrow in each genotype.

**Extended Data Fig. 6.**
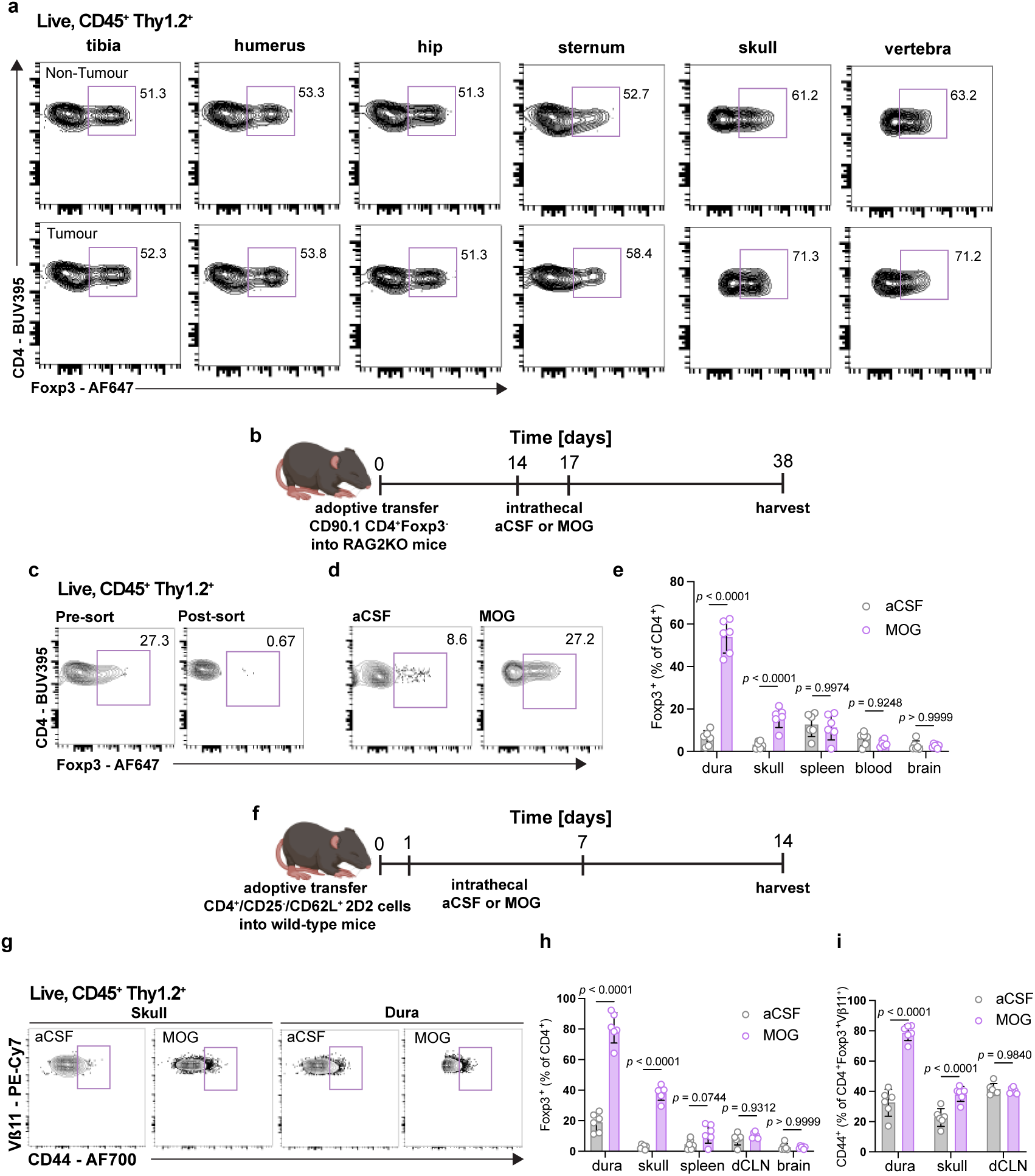
Naïve CD4 T cells are polarised to Tregs in the meningeal dura mater and skull bone marrow. **a**, Representative flow cytometry analysis of CD4^+^FOXP3^+^ T cells across different bone marrow niches in *EP^ZFTA-RELA^*-bearing and control bearing mice. **b**, Schematic of the adoptive transfer system from non-Treg CD4^+^ T cells (Thy1.2) cells into RAG2 knockout recipients (KO), created in biorender.com. **c**, Post-sort validation of isolation of CD4^+^Foxp3^-^ Thy1.2^+^ cells. **d**, Flow cytometry analysis and quantification of CD4^+^FOXP3^+^ T cells in different tissues, n = 6/group, Data are means ± S.E.M, p values represent two-way ANOVA with Sidak’s post hoc test. **e**, Quantification of the proportion of Foxp3^+^ CD4 T cells across bone marrow niches in *EP^ZFTA-RELA^*-bearing and control bearing mice, combined in. n=8. Data are means ± S.E.M, p values represent two-way ANOVA with Sidak’s post hoc test. **f**, Schematic of adoptive transfer experiments using 2D2 naïve (CD4^+^VB5.1^+^CD25^-^CD62L^+^) T cells to wildtype recipients. **g**, Flow cytometry analysis and quantification of Tregs (CD4^+^Foxp3^+^). **h**, and antigen-specific activated Tregs (Foxp3^+^CD44^+^VB11^+^) (**i**), respectively. n = 6, data are means ± S.E.M, p values represent two-way ANOVA with Sidak’s post hoc test.

**Extended Data Fig. 7.**
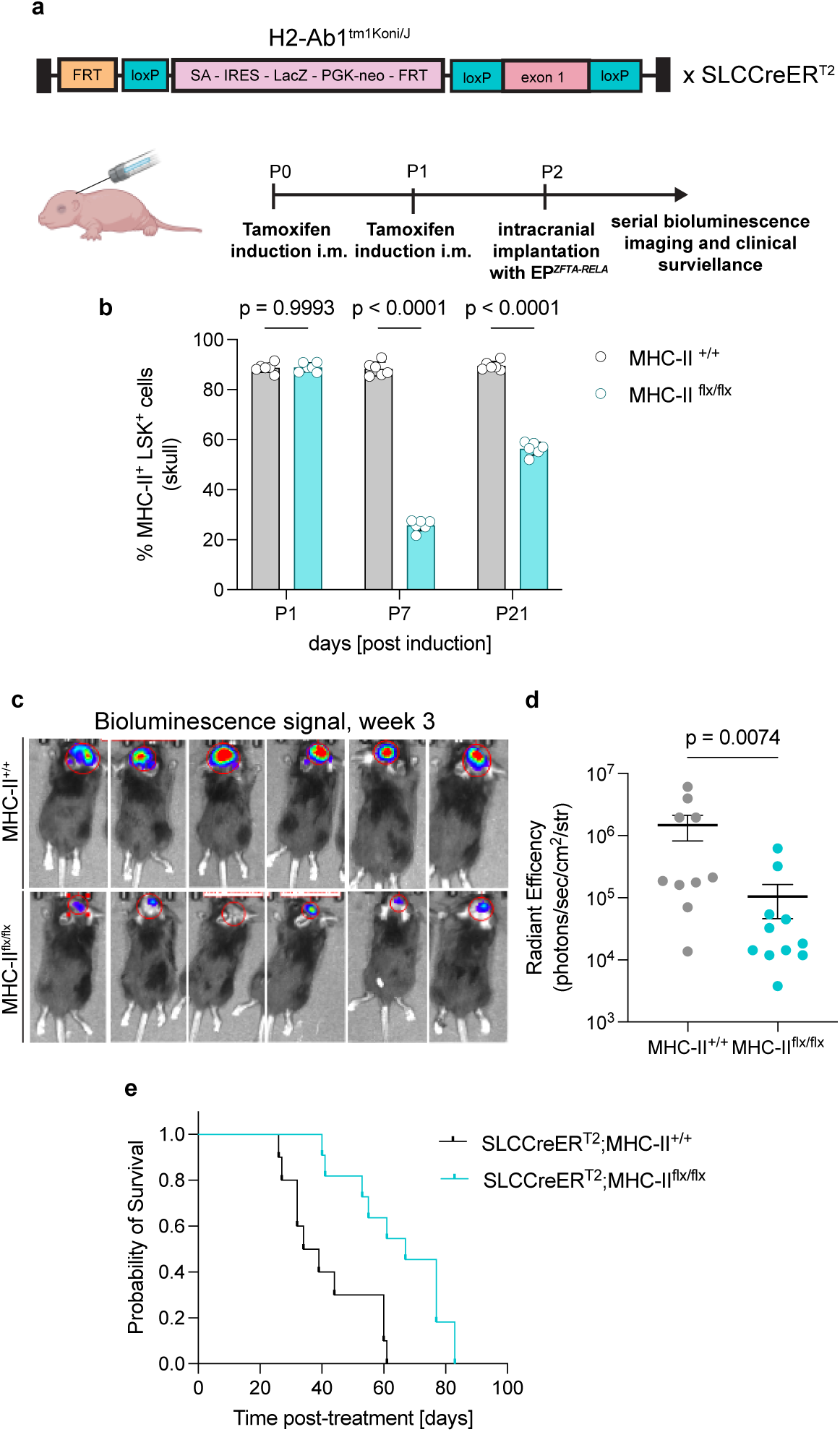
Conditional deletion of MHC-II on HSPCs reduces tumour burden in EP^ZFTA-RELA^. . **a**, Schematic of the H2-Ab1^tm1Koni/J^ cassette use to generation the MHC-II^flx/flx^ mouse following cross with SLC-CreER^T2^ model and induction and implantation timelines, created in biorender.com. **b**, Quantification of the proportion of MHC-II expressing LSK cells in the skull bone marrow of MHC-II^+/+^ and MHC-II^flx/flx^ mice up to 21 days (n = 6 per group, 2 independent experiments). **c**, Representative bioluminescent images of 3 week-old mice in each group. **d**, Quantification of bioluminescent signal at 3 weeks after induction and implantation. **e**, Kaplan–Meier survival plot of MHC-II^+/+^ (n = 10) and MHC-II^flx/flx^ (n = 11)

**Extended Data Fig. 8.**
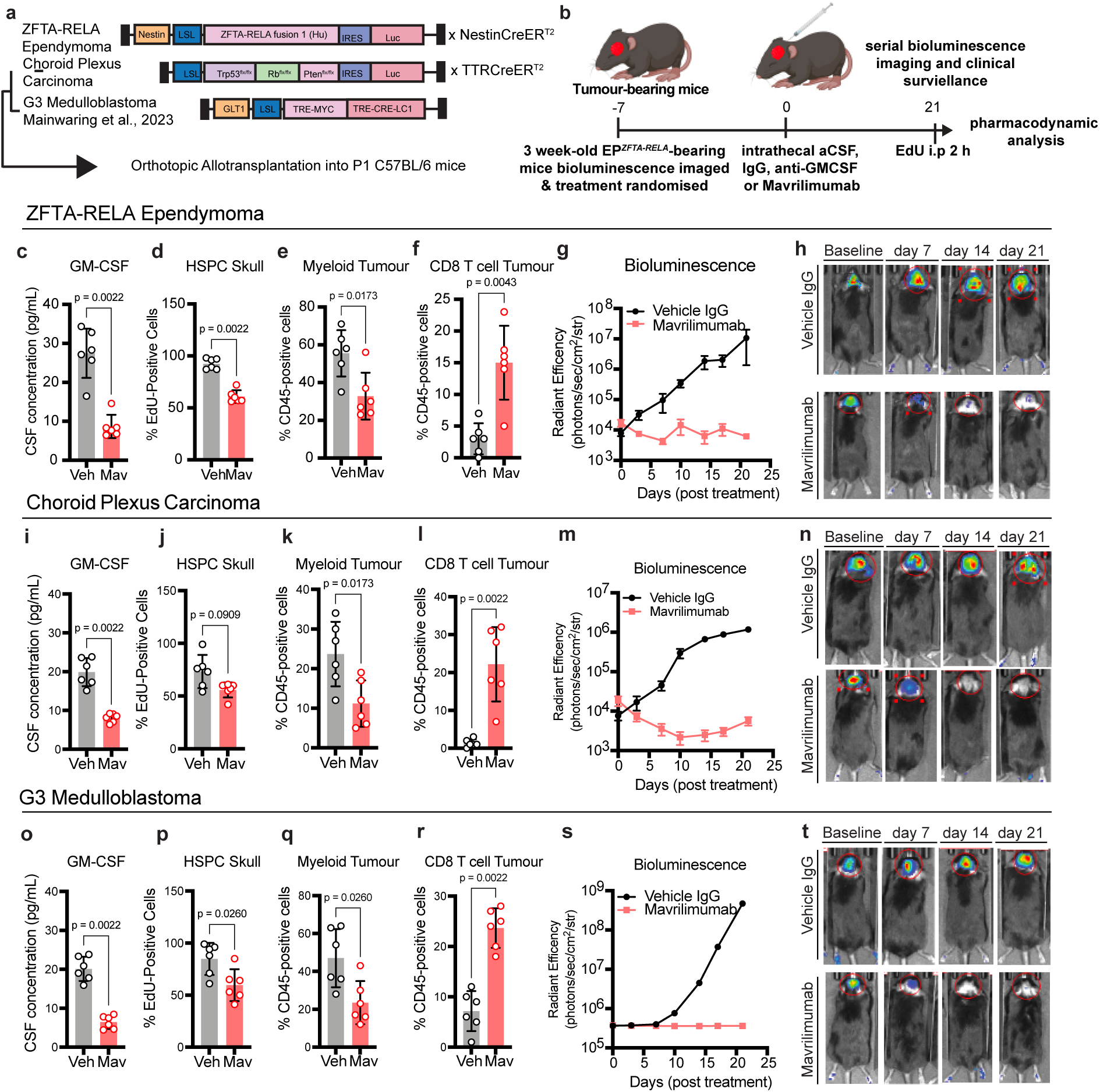
Anti-tumour effects of a single intrathecal dose of Mavrilimumab across multiple mouse models of rare childhood brain tumours. **a**, schematic of constructs of childhood brain tumour mouse models used in subsequent studies, including ZFTA-RELA ependymoma, choroid plexus carcinoma and group 3 medulloblastoma. **b**, Experimental design for the treatment of 3 week- old *EP^ZFTA-RELA^*-bearing mice with a single intracisternal magna (ICM) injection of 10 uL of IgG isotype control (Veh, 5 mg/kg, *n* = 6 per group or mavrililumab (Mav, 5 mg/kg, *n* = 6 per group) created with biorender.com. **c, i, o**,. Enzyme-linked immunosorbent assay (ELISA) quantification of CSF levels of GM-CSF 21 days after intra cisterna magna injection (icm) of Vehicle IgG Isotype control (5 mg/kg) or Mavrililumab (5 mg/Kg) in ZFTA-RELA ependymoma, G3 Medulloblastoma and Choroid Plexus Carcinoma mouse models. **d, j, p**, Flow cytometry quantification of the perecentage of EdU positive Live CD45^+^ CD45 i.v- Lineage-Sca1^+^c-Kit^+^ HSPCs in the skull bone marrow of mice 21 days post treatment. **e, k, q**, Flow cytometry quantification of the proportion of myeloid (Live CD45^+^ CD45 i.v- Ly6G- Cd11b^+^) cells in the tumour parenchyma. **f, l, r**, Flow cytometry quantification of the proportion of CD8 T cells in the tumour parenchyma after treatment for 21 days. **g, m, s**, Tumour progression was then followed by bi-weekly bioluminescence imaging and clinical surveillance. **h, n, t**, Representative bioluminescence signal across one mouse from each group in each tumour model. Data represents, mean± SD (n=6/group) unpaired two-tailed Student’s t-test).

**Extended Data Fig. 9.**
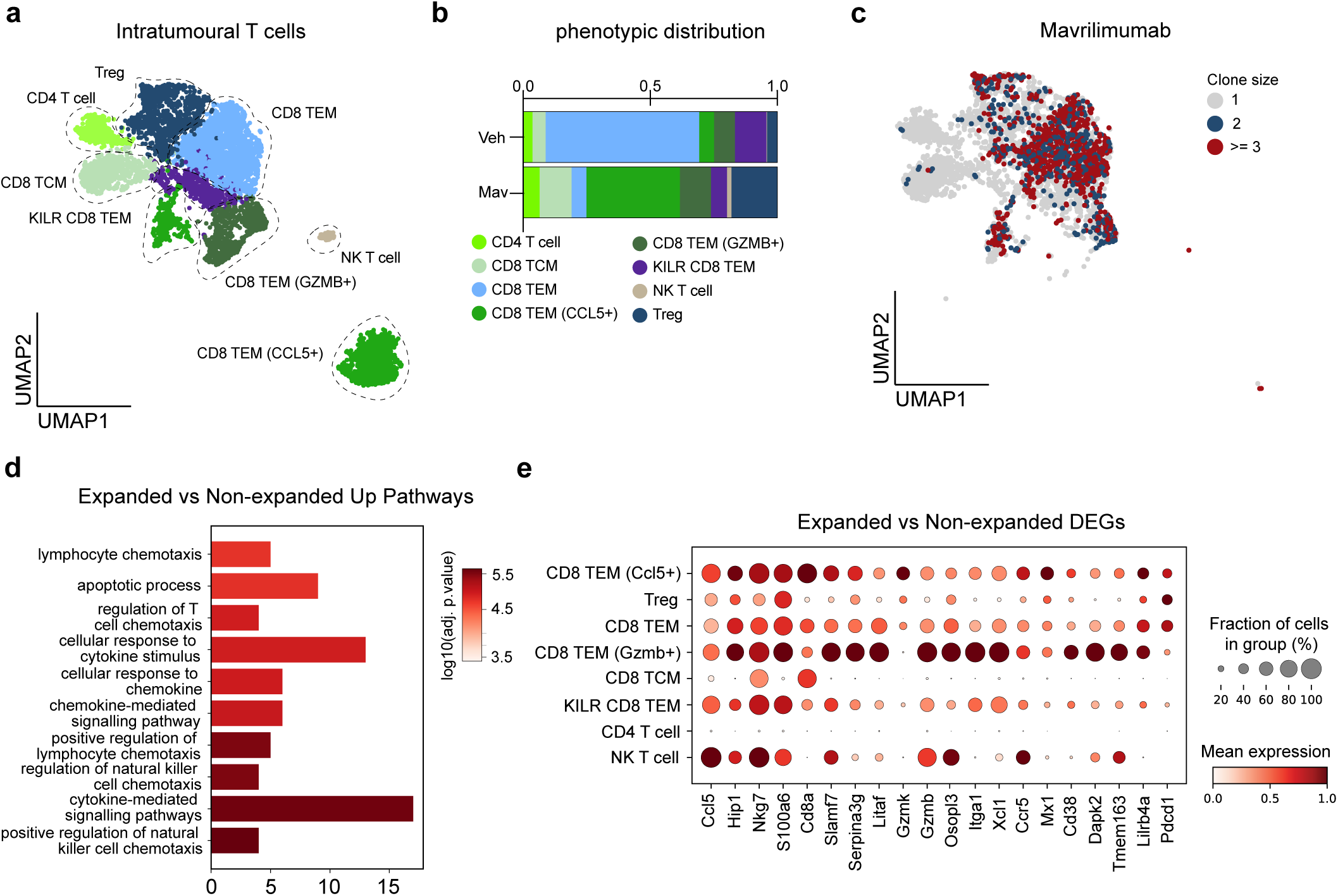
Normalizing skull haematopoiesis promotes clonal expansion and tumour infiltration of effector T cells. a, Uniform-manifold projection of intratumoural T cells in endpoint vehicle and mavrilimumab-treated EP^ZFTA-RELA^-bearing mice, coloured by phenotype and quantified in **b. c,** UMAP of clone size of intratumoural T cells in mavrilimumab-treated mice. **d,** Gene Ontology (GO) pathway analysis top differentially upregulated genes in expanded intratumoural T cell clones (>3) relative to non-expanded clones. **e,** dotplot of top 20 differentially upregulated genes across T cell phenotype clusters in mavrilimumab-treated mice, n = 2 pooled experiments per group.

**Extended Data Fig 10.**
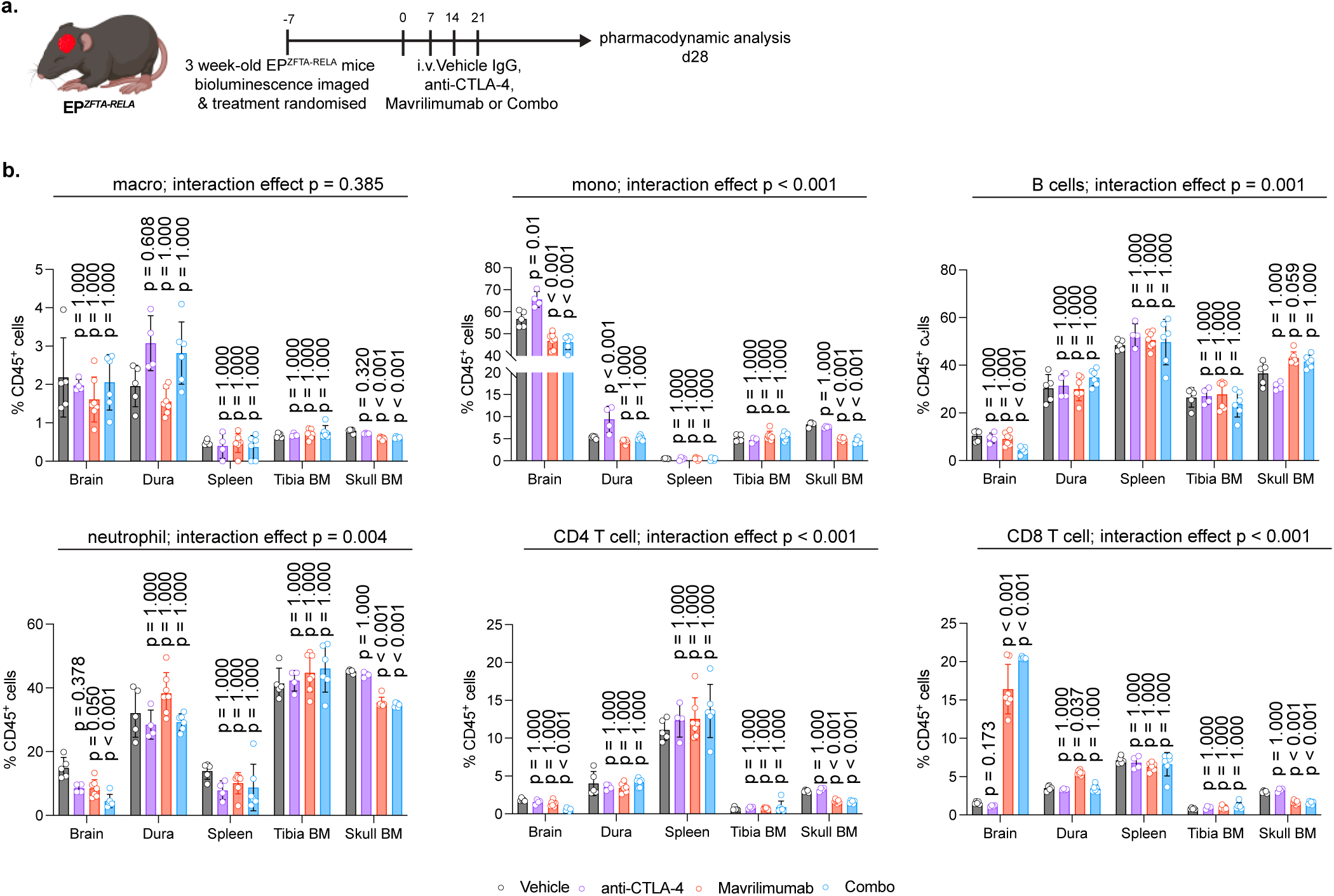
Combinatorial treatment of mavrilimumab with immune-checkpoint blockade synergizes for anti-tumour immunity without off-target effects in EP*^ZFTA-RELA^* mice. **a**, Experimental design for the treatment of 3 week-old ZFTA-RELA EP*^ZFTA-RELA^*-bearing mice with four weekly intravenous injections of Vehicle IgG isotype control (13 mg/kg, n = 5), anti-CTLA-4 (3 mg/kg, n = 4), mavrilimumab (10 mg/kg, n = 7) or combination of anti-CTLA-4 and mavrilimumab (n = 6), created with biorender.com. **b,** Quantification of the proportion of macrophages, monocytes, B cells, neutrophils, CD4 T cells and CD8 T cells, within the brain, dura, spleen, tibia and skull bone marrow of treated mice. A linear mixed-effects model was applied to test for interaction across strata, with skull bone marrow as the reference, Wald z-tests on contrasts relative to vehicle with Bonferroni correction for multiple comparisons.

## SUPPLEMENTAR FIGURE LEGENDS

**Supplementary Fig. 1 Autochthonous Nestin^CreERT2^;Nestin^Flx-STOP-FlxZFTA-RELA^ mouse model of ZFTA-RELA fusion-driven ependymoma. a,** Schematic of lox-stop-lox cassette and breeding strategy for the generation of the Nestin^CreERT2^;Nestin^Flx-STOP-FlxZFTA-RELA^ mouse model. **b,** Kaplein Meier survival of Nestin^CreERT2^;Nestin^Flx-STOP-FlxZFTA-RELA^ mouse model (n = 70), Log-rank MantelCox test relative to Nestin^CreERT2^ control mice (n = 50), median survival 92 ± 16 days. Representative bioluminecense signal of 28 day old EP^ZFTA-RELA^ mice. **c,** Representative brightfield TdTomato signal and haematoxylin and Eosin staining of endpoint tumour. **d,** Representative immunohistochemical staining of Nestin^CreERT2^;Nestin^Flx-STOP-FlxZFTA-RELA^ mouse tumours stained for canonical ZFTA-RELA ependymoma markers**. e**, uniform manifold approximation and projection (UMAP) visualisation of single-cell RNAseq of tumour parenchyma of EP^ZFTA-RELA^ mice (n = 4 mice), coloured by cell-type annotations; neural stem cell (NSC) proliferating (prolif.) radlial glia, neural precursor cell (NPC), astrocyte, NSC, oligodendrocyte precursor cell (OPC), stromal, myeloid, lymphoid, neuronal and oligodendrocyte. **f**, enrichment plot of ependymoma signature score across cell types. **g**, Dotplot of proportion of and mean expression of ZFTA-RELA gene across cell types.

**Supplementary Fig. 2 Representative flow cytometry gating strategies. a,** Gating strategy for the annotation of haematopoietic stem progenitor cell (HSPC) populations in murine tissue; multipotent progenitor (MPP)3/4, MPP2, MPP1, haematopoietic stem cell (HSC), granulocyte-monocyte precursor (GMP) pre-megakaryocyte/erythrocyte (PreMegE), pre- granulocyte monocyte (preGM), pre-colony- forming unit-erythroid cell (CFU-E), CFU-E, LSK-, CD8 T cell (TC), CD4 TC, neutrophil (Neutro.), monocyte (Mono.), macrophage (Macro.) eosinophil (EOS), B cell (BC) and dendritic cell (DC). **b**, Gating strategy for fluorescence-activated cell sorting (FACS) of immune cells for antigen presentation assays.

**Supplementary Fig. 3 Integration of PTPRC^+^ cells from human foetal brain and childhood brain tumour single-cell RNA sequencing datasets. a**, schematic summarising the sourcing of previously published datasets created in biorender.com. **b**, uniform manifold approximation and projection (UMAP) visualisation of single-cell RNAseq of integrated from human fetal brain and childhood brain tumour PTPRC^+^ cells, annotated by cell type; B cells CD4 tissue central memory (TCM), CD4 Tissue effector memory (TEM). CD8 T cell, HSPCs, interferon-gamma (IFN-y) macrophages, IFN-y microglia, natural killer (NK) T cell, Purinergic Receptor P2Y12 (P2RY12) low microglia, Treg, conventional dendritic cell (cDC) 1/2 complement myeloid, foetal, infantile and resting microglia, monocyte, neutrophil and pro-inflammatory (pro-inflam.) macrophage and microglia. **c**, Stacked barplot of the proportion of annotated cell types across developmental stages. **d-e**, UMAP annotated by pathology (anaplastic astrocytoma (a.astrocytoma), anaplastic glioma (a.glioma), diffuse intrinsic pontine glioma (DMG), group 3 (Gr3) and group 4 (Gr4) medulloblastoma (MB), posterior-fossa -type A (PFA) ependymoma type 1 and 2, posterior-fossa -type B (PFB) ependymoma, sonic-hedghog (SHH) medulloblastoma, Supratentorial-REL-associated protein (ST-RELA) ependymoma (EPN), ST-Yes1 associated transcriptional regulator (YAP) EPN) and development stage. **f**, Dotplot of mean expression and fraction of cells in group expressing cell type annotations genes across annotated cell types.

**Supplementary Fig. 4 Human choroid plexus carcinoma neurosurgical CNS immune tissue reveals HLA-DR-expressing haematopoietic stem progenitor cells (HSPCs) in skull, dura and tumour parenchyma. a**, Representative histogram showing HLA-DR expression levels in immune cell populations across tissues. **b**, Representative t-SNE of flow cytometry data from tumour parenchyma, dura mater and skull bone marrow with 3000 events concatenated per sample, manually gated; HSPCs, CD8 tissue central memory (TCM), CD45RA^+^ CD16^+^ monocytes, CD14^-^CD66b^+^CD16^+^CD45^lo^, CD14^-^ CD66b^+^CD16^+^CD45^hi^, CD66b^+^CD14^-^, B cells, neutrophils, non-classical monocytes, PDL-1^lo^ monocye-derived macrophages (MDMs), natural killer (NK) cells CD16^+^CD56^+^ NK cells, dendritic cells, CD4 T cells, double-negative (DN) T cells, CD8 T cell, regulatory T cell (Treg), CD4 Naïve T cell, CD4 tissue effector memory (TEM), CD45RA^+^ CD90^+^ T cell, CD4 TCM.

**Supplementary Fig. 5 Local increased proliferation of skull bone marrow cells in** *EP^ZFTA-RELA^***- bearing mice, which is absent in peripheral bone marrow niches. a,** Quantification of the percentage of 5-Ethynyl-2′-deoxyuridine (EdU)-positive flow cytometry profiled LSK^+^ HSPCs from the skull and tibia bone marrow of *EP^ZFTA-RELA^ -*bearing and *and Nestin^CreERT2^*, control bearing mice, n = 6 individual mice, average of 3 independent experiments, data are means ± s.e.m.; P values represent a two-sided Student’s t-test. **b**, Number of CD45^+^ cells in skull and tibia of *EP^ZFTA-RELA^*-bearing and control bearing mice, quantified from **a. c,** Quantification of the percentage of EdU-positive immune cell subsets, n = 6 individual mice, average of 3 independent experiments, data are means ± s.e.m.; P values represent a two-sided Student’s t-test. **d**, Quantification of the proportion of subsets of CD45^+^ cells within the skull of EP*^ZFTA-RELA^* -bearing *and Nestin^CreERT2^* mice as measured by flow cytometry, (n=6/group, mean ± s.e.m, unpaired two-tailed Student’s t-test). **e-f,** Quantification of the percentage of EdU-positive flow cytometry profiled LSK^+^ HSPCs from the skull and tibia bone marrow of orthotopic allotransplantation models of choroid plexus carcinoma and group 3 medulloblastoma, relative to sham control mice, n = 6 individual mice, average of 3 independent experiments, data are means ± s.e.m.; P values represent a two-sided Student’s t-test.

**Supplementary Fig. 6 Experimental pipeline and cell type annotation of single-cell RNA- sequencing of mouse neuroimmune tissues in** *EP^ZFTA-RELA^***-bearing and control bearing mice. a,** Schematic illustrating the methodology to isolate CD45^+^ cells from the tissues listed in *EP^ZFTA-RELA^*- bearing and control bearing mice, created in biorender.com. **b**, Representative gating strategy for the fluorescence-activated cell sorting (FACS) isolation of CD45 i.v- Terr119^-^ Live CD45^+^ cells. **c-e,** t- distributed stochastic neighbour embedding (t-SNE) visualizations of scRNA-seq from brain, deep- cervical lymph nodes (dcLN), dura, skull and tibial bone marrow from 4-week-old mice coloured by genotype, tissue – split by genotype and cell type. **f**, Stacked bar plot of the relative proportions of annotated scRNA-seq cell types within the skull and tibia bone marrow of *EP^ZFTA-RELA^*-bearing and control bearing mice. **g**, Mean expression dot plot of marker genes used to define cell type annotations in single cell RNAseq data from the these tissues in *EP^ZFTA-RELA^*-bearing and control bearing mice. **h**, Volcano plots of differentially expressed genes in neutrophils, monocytes, macrophages, and HSPCs within the skulls of tumour bearing relative to control bearing mice. Magenta dots represent upregulated transcripts, blue dots represent downregulated transcripts in skull populations compared to the tibia. y-axes represent adjusted log_2_ p value for cluster changes between tumour and control bearing mice. **i,** Dotplot of average and percentage expression of T cell instructing cytokines, co-inhibition and co-stimulation molecules across HSPC and myeloid progenitor cell clusters.

**Supplementary Fig. 7 CSF access and modulate skull bone marrow niche in** EP*^ZFTA-RELA^***-bearing mice. a**, Experimental design of intracisterna magna (ICM) injections of anti-c-Kit BV421 and OVA– 488 in tumor and non-tumor-bearing mice (n=5 per group); image created with Biorender.com. **b**, Gating strategy to quantify dual-positive c-Kit^+^ cells, summarized in (f). **c**, Gating strategy to quantify OVA-488^+^ macrophages in dura, tibia, and skull bone marrow. **d-e**, Representative flow cytometry plots of dual-positive c-Kit cells (**d**) and OVA-positive F4/80^+^ macrophages (**e**) across experimental conditions. **f**, Quantification of the proportion and number of intratumoral macrophages, CD8 T cells, haematopoietic stem progenitor cells (HSPCs) and neutrophils following intracisternal magna injection of AMD3100 (2 mg/kg, 6 h) or artificial CSF (aCSF) treatment (n=5/group, mean±s.e.m, unpaired two- tailed Student’s t-test) relative to aCSF).

**Supplementary Fig. 8 Combined analysis of chromatin accessibility and gene expression in HSCs from skull and tibia of** EP*^ZFTA-RELA^***-bearing and control-bearing mice. a**, Uniform manifold approximation and projection (UMAP) visualization of combined single nucleus assay for transposase- accessible chromatin using sequencing (snATAC-seq) and RNA data using latent semantic indexing (LSI) from fluorescence-activated cell sorting (FACS)-isolated CD34^+^ LSK^+^ (haematopoietic stem cell (HSC), showing distinct clustering of hematopoietic subpopulations, common myeloid progenitor (CMP), granulocyte monocyte progenitor (GMP), Pro B, and Pro T) across skull and tibia samples. **b**, UMAP plots of snATAC (top) and RNA (bottom) data, stratified by tumour and control conditions, highlighting the distribution of hematopoietic subpopulations in skull and tibia. **c**, Feature plots of key lineage-specific genes (Pax5, Gata1, Ms4a1, Cd14, Cd3d, Cd8a, Cd34, and Mpo), illustrating expression dynamics across hematopoietic differentiation trajectories. **d**, Heatmap of the peak matrix, showing Z-scores for differentially accessible chromatin regions across tumour and control skull and tibia samples. **e**, Heatmap of the combined gene score matrix across pseudo time, highlighting dynamically regulated genes (Spib, Siglech, Ppp1r16b, Arl5c, and Dusp2) associated with differentiation. **f**, Heatmap of the motif matrix, showing activity scores of transcription factor motifs (Cebpb, Cebpg, Zbtb16, Fosb, Bach1, and Smarcc1) across pseudotime, with increased motif activity in EP*^ZFTA-RELA^*-bearing samples indicative of enhanced regulatory engagement. **g**, Quantification of the number and types of colonies (colony forming unit (CFU)- granulocytes/erythroids/macrophages/megakaryocytes (CFU-GEMM), colony-forming unit- granulocytes/macrophages (CFU-GM) and colony-forming unit-erythroid (CFU-E) in colony forming cell assays derived from skull bone marrow from EP*^ZFTA-RELA^*-bearing and control bearing mice, (n=6/group, mean±s.e.m, unpaired two-tailed Student’s t-test).

**Supplementary Fig. 9 Single-cell RNA-sequencing highlights CSFR2A expression on skull and dura in *EP^ZFTA-RELA^*-bearing mice. a**, Heatmap depicting GM-CSF (Csf2)-associated cytokine enrichment scores generated using CytoSig for annotated cell types across skull, dura, and brain from control and EP^ZFTA-RELA^ -bearing mice. **b**, Dot plot of receptor expression in skull bone marrow cells of *EP^ZFTA-RELA^*-bearing mice, scaled by gene expression and percentage of cells expressing the gene. **c**, Smooth density plots of CSFR2a enrichment scores across selected immune cell types in the Skull and Dura tissues. **d**, Dot plot of CSF2 expression in across cells and tissues of *EP^ZFTA-RELA^*-bearing and control bearing mice, scaled by gene expression and percentage of cells expressing the gene.

**Supplementary Fig. 10. Increased T cell infiltrate in single-cell RNAseq of intratumoural immune compartment following treatment with mavrilimumab in EP*^ZFTA-RELA^* mice. a**, Experimental design for the treatment of 3 week-old *EP^ZFTA-RELA^*-bearing mice with a single intracisternal magna (ICM) injection of 10 uL of IgG isotype control (5 mg/kg, *n* = 6 per group or mavrililumab (5 mg/kg, *n* = 6 per group) created with biorender.com. **b**, Representative flow cytometry plots for the enrichment of extravascular CD45^+^ cells. **c**, Uniform manifold approximation and projection visualisation of CD45^+^ cells in integrated vehicle and mavrilimumab single-cell RNAseq datasets. **d**, Dotplot of expression of T cell lineage and phenotype markers across annotated cell types. **e-g**, Featureplot of key phenotypic marker expression levels including **(e)** lineage and memory, **(f)** exhaustion and activation, **(g)** effector molecule, and **(h)** transcription factors genes. **i**, Flow cytometry quantification of the proportion of CTLA-4^+^ CD8 T cells in tumour parenchyma of treated mice at recurrence, n = 12 mice per group.

