## Supplementary Tables for "Childhood brain tumours instruct cranial haematopoiesis and immunotolerance"

**Supplementary Table 1: List of mouse strains**

| Mouse Strain | Company |
|---|---|
| Mouse: *Rosa-Cre-ER* $^{T2}$ | Jackson Laboratory |
| Mouse: Nestin-CreERT2 | Collaoration |
| Mouse: C57BL/6 | Jackson Laboratory |
| Mouse: B6.Cg-Gt(ROSA)26Sortm14(CAG-tdTomato)Hze/J | Jackson Laboratory |
| Mouse: C57BL/6-Tg(Tcra2D2,Tcrb2D2)1Kuch/J | Jackson Laboratory |
| Mouse: B6.Cg-Tg(TcraTcrb)425Cbn/J | Jackson Laboratory |
| Mouse: B6.Cg-Rag2tm1.1Cgn/J | Jackson Laboratory |
| Mouse: B6.SJL-Ptprca Pepcb/BoyJ | Jackson Laboratory |
| Mouse: C57BL/6-Tg(CAG-OVAL)916Jen/J | Jackson Laboratory |
| Mouse: B6.Cg-Rag2tm1.1Cgn/J | Jackson Laboratory |
| Mouse: B6.129X1(FVB)-H2-Ab1b-tm1.2Koni/KmmJ | Jackson Laboratory |
| Mouse: C57BL/6-Tg(Tal1-cre/ERT)42-056Jrg/J | Jackson Laboratory |

**Information**

RRID: IMSR_JAX:008463

MGI:5304277

RRID:IMSR_JAX:000664

RRID:IMSR_JAX:007914

RRID:IMSR_JAX:006912

RRID:IMSR_JAX:004194

RRID:IMSR_JAX:008449

RRID:IMSR_JAX:002014

RRID:IMSR_JAX:005145

RRID:IMSR_JAX:008449

RRID:IMSR_JAX:037709

RRID:IMSR_JAX:037466
