## Supplementary figures and images for "Childhood brain tumours instruct cranial haematopoiesis and immunotolerance"

### SUPPLEMENTARY FIGURE 1

#### SUPPLEMENTARY FIGURE 1

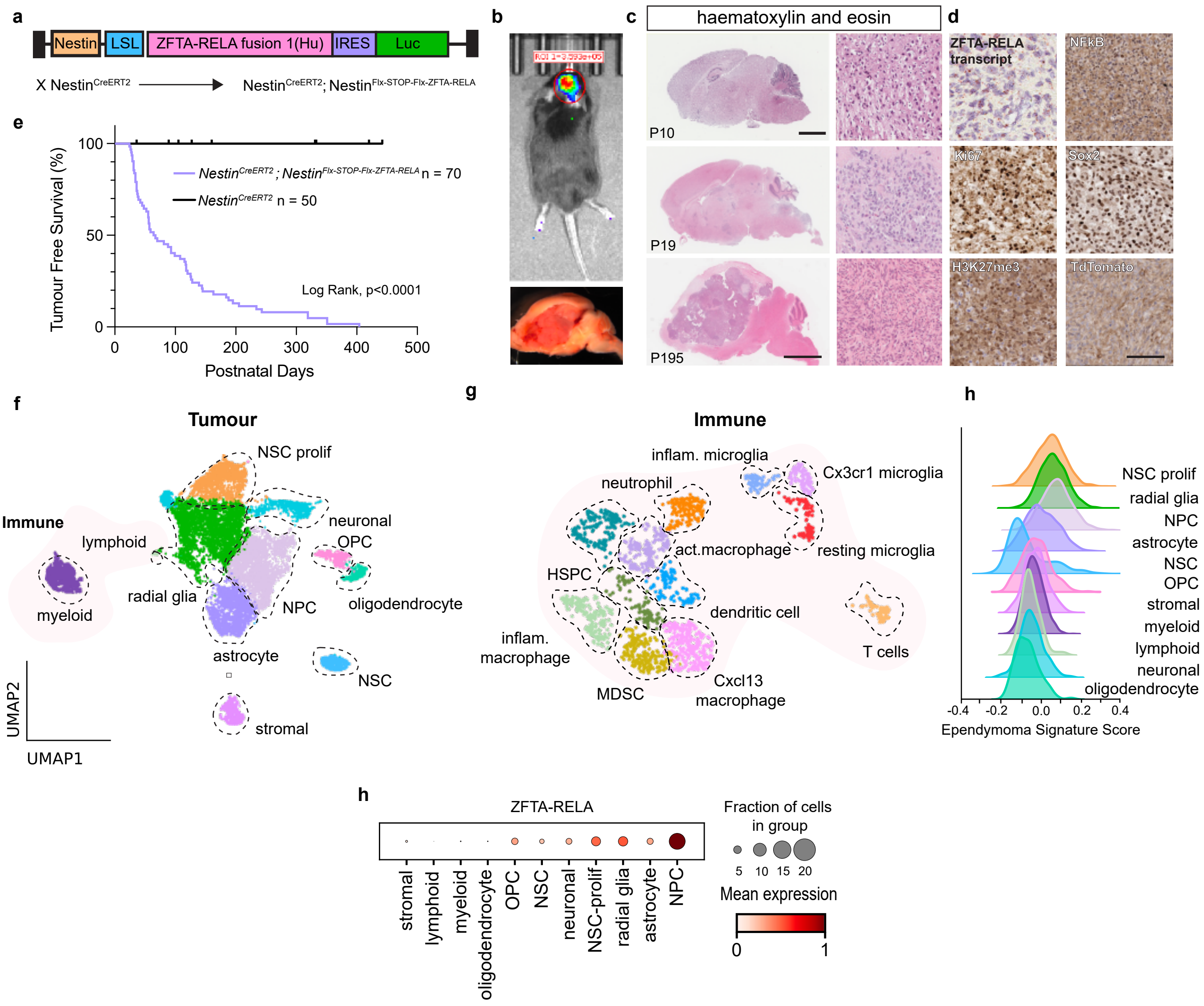

### SUPPLEMENTARY FIGURE 2

#### SUPPLEMENTARY DATA FIGURE 2

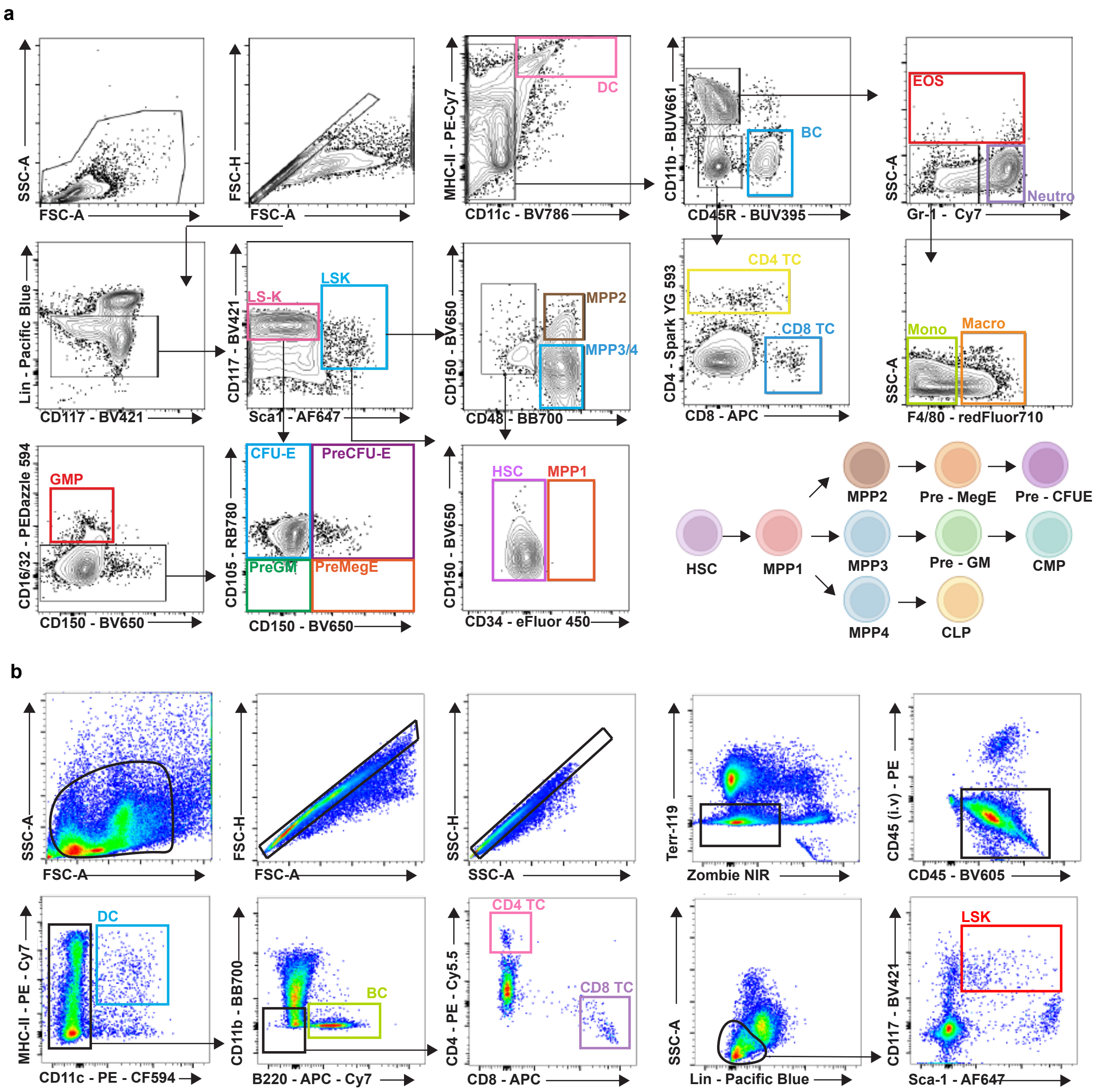

### SUPPLEMENTARY FIGURE 3

#### SUPPLEMENTARY FIGURE 3

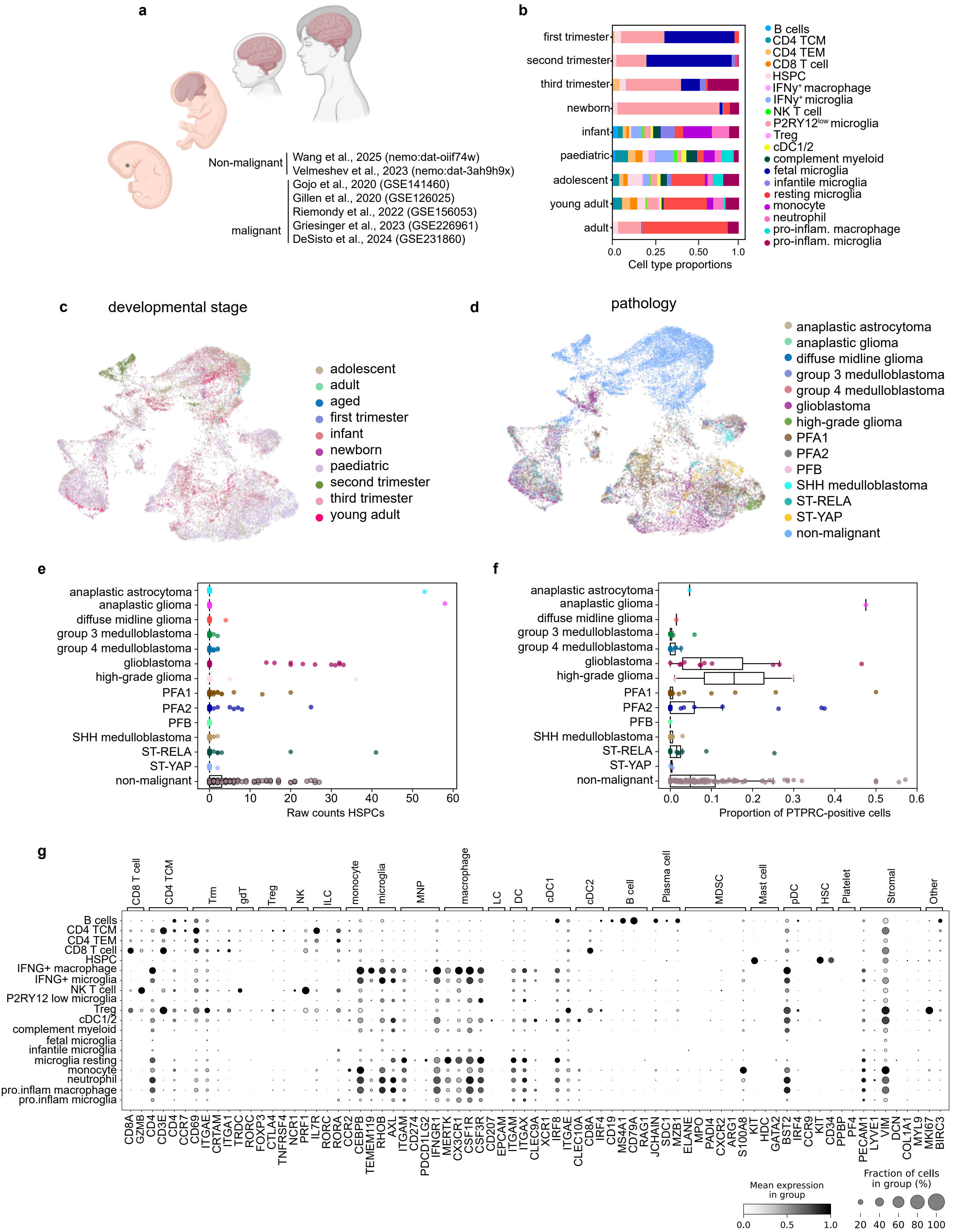

### SUPPLEMENTARY FIGURE 4

**SUPPLEMENTARY FIGURE 4**

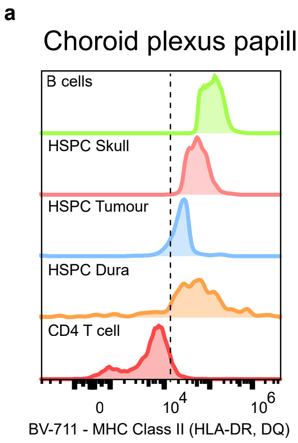


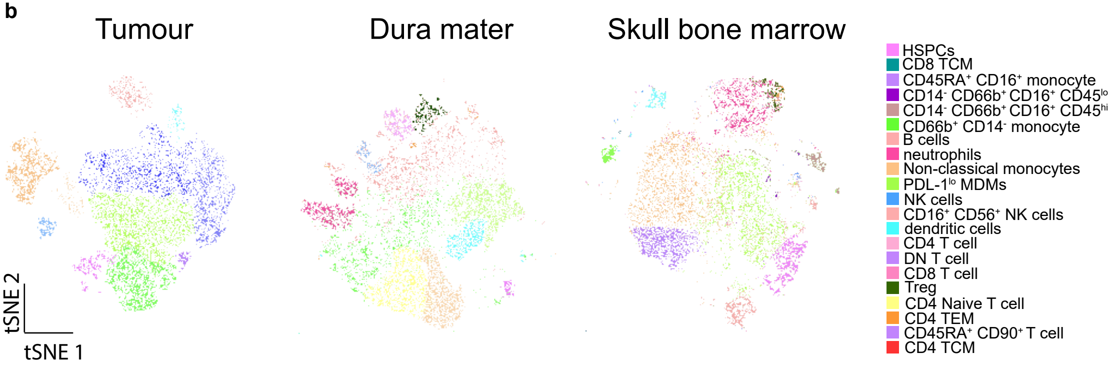

### SUPPLEMENTARY FIGURE 5

#### SUPPLEMENTARY FIGURE 5

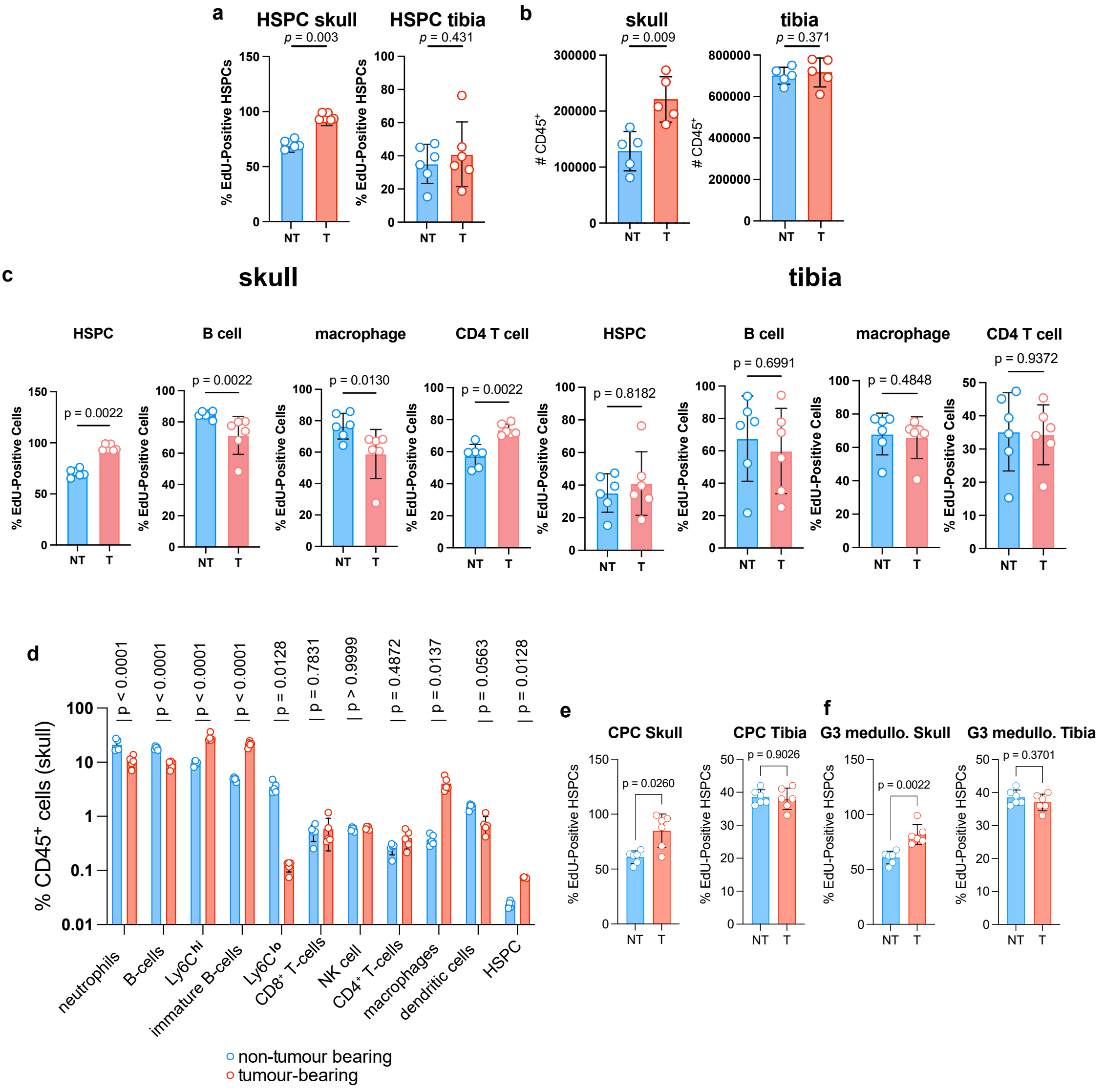

### SUPPLEMENTARY FIGURE 6

##### SUPPLEMENTARY FIGURE 6

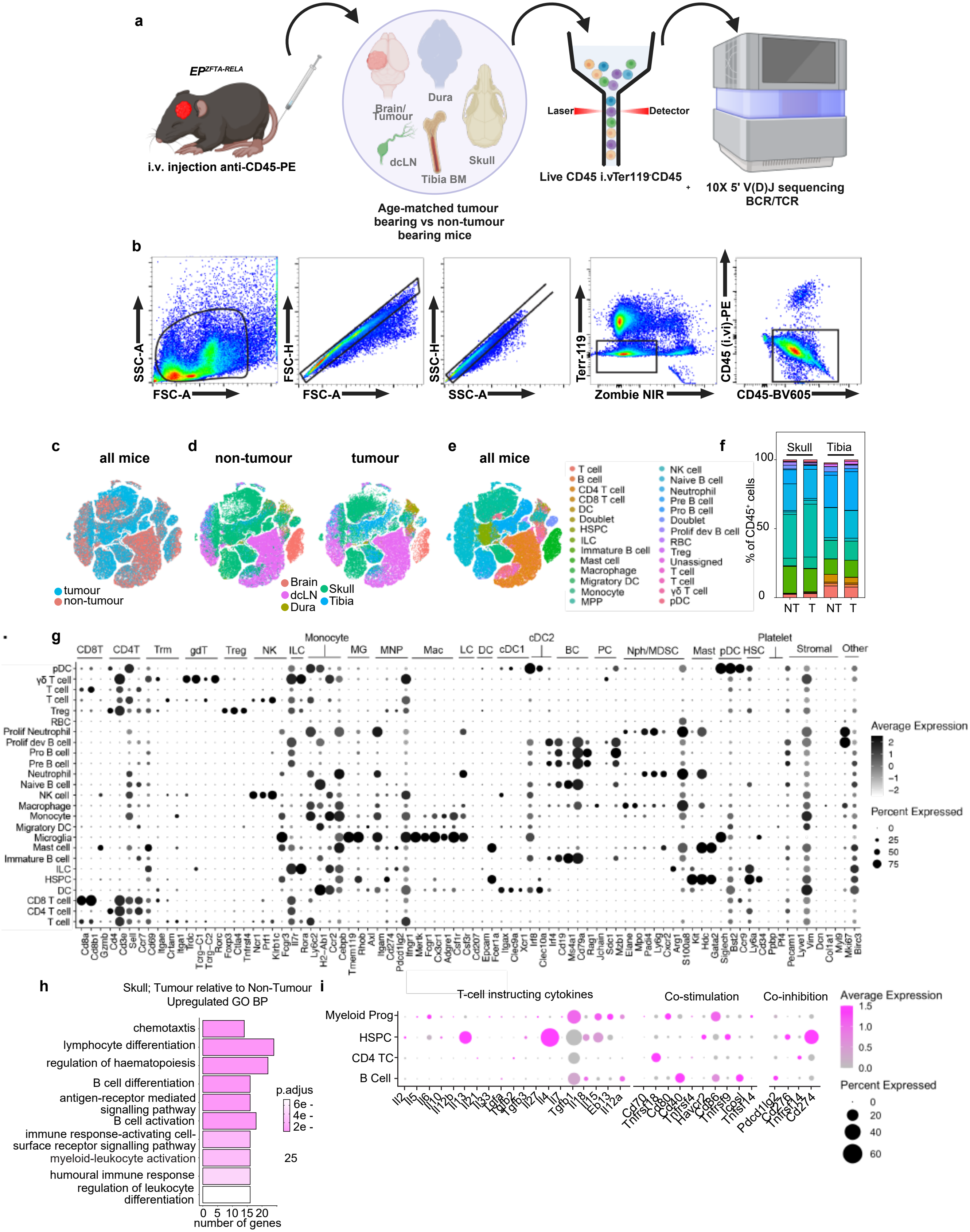

### SUPPLEMENTARY FIGURE 7

#### SUPPLEMENTARY DATA FIGURE 7

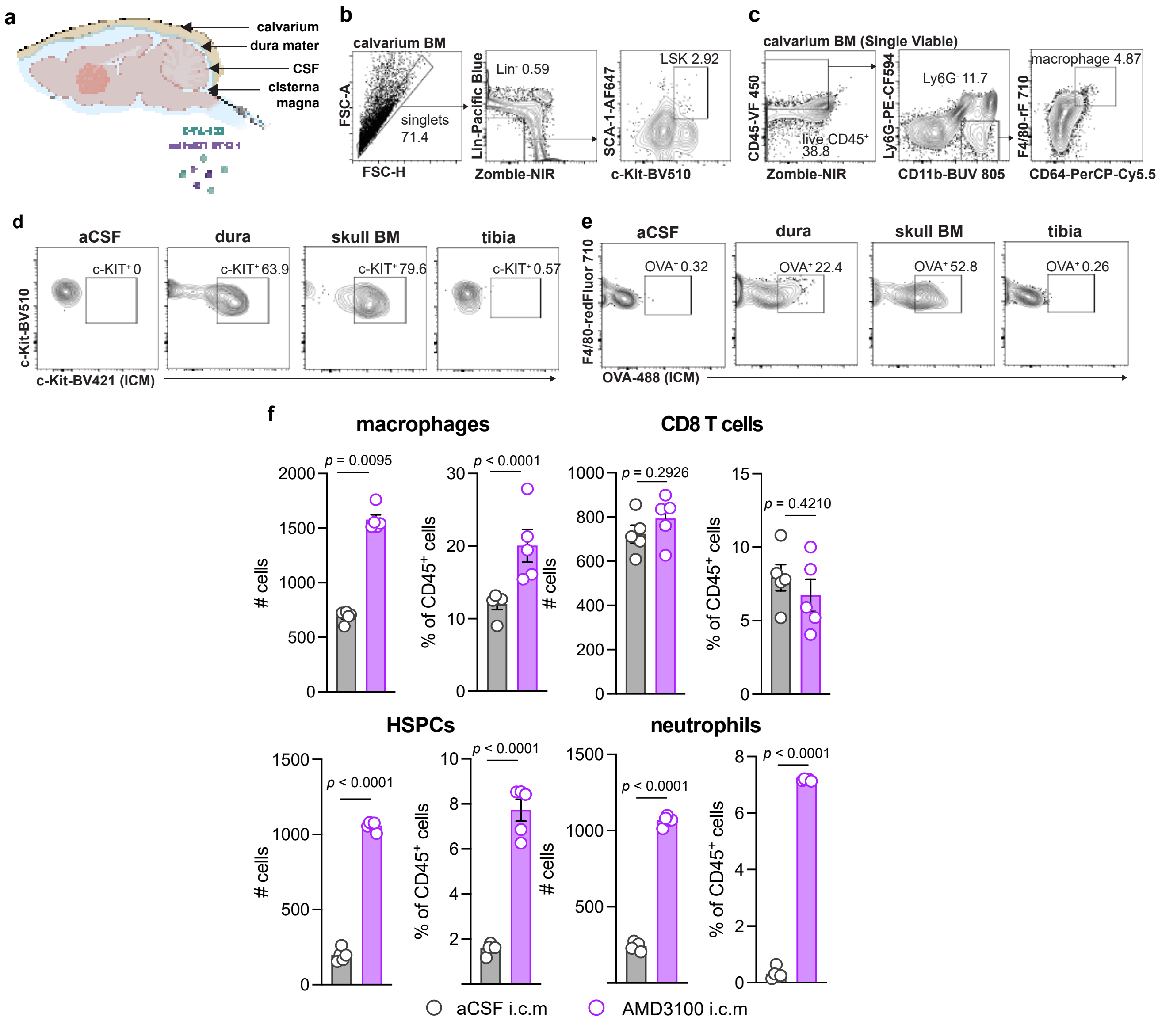

### SUPPLEMENTARY FIGURE 8

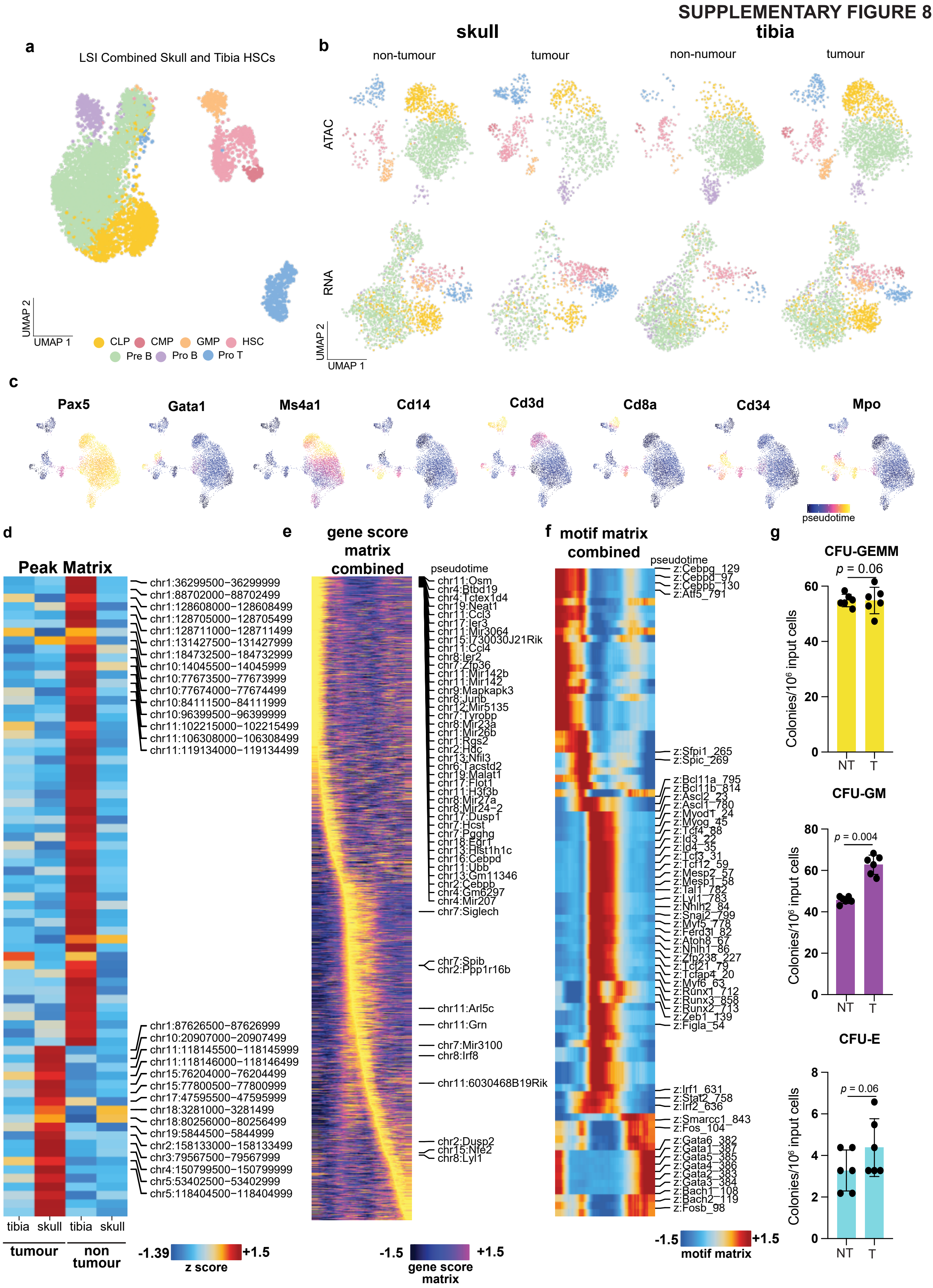

### SUPPLEMENTARY FIGURE 9

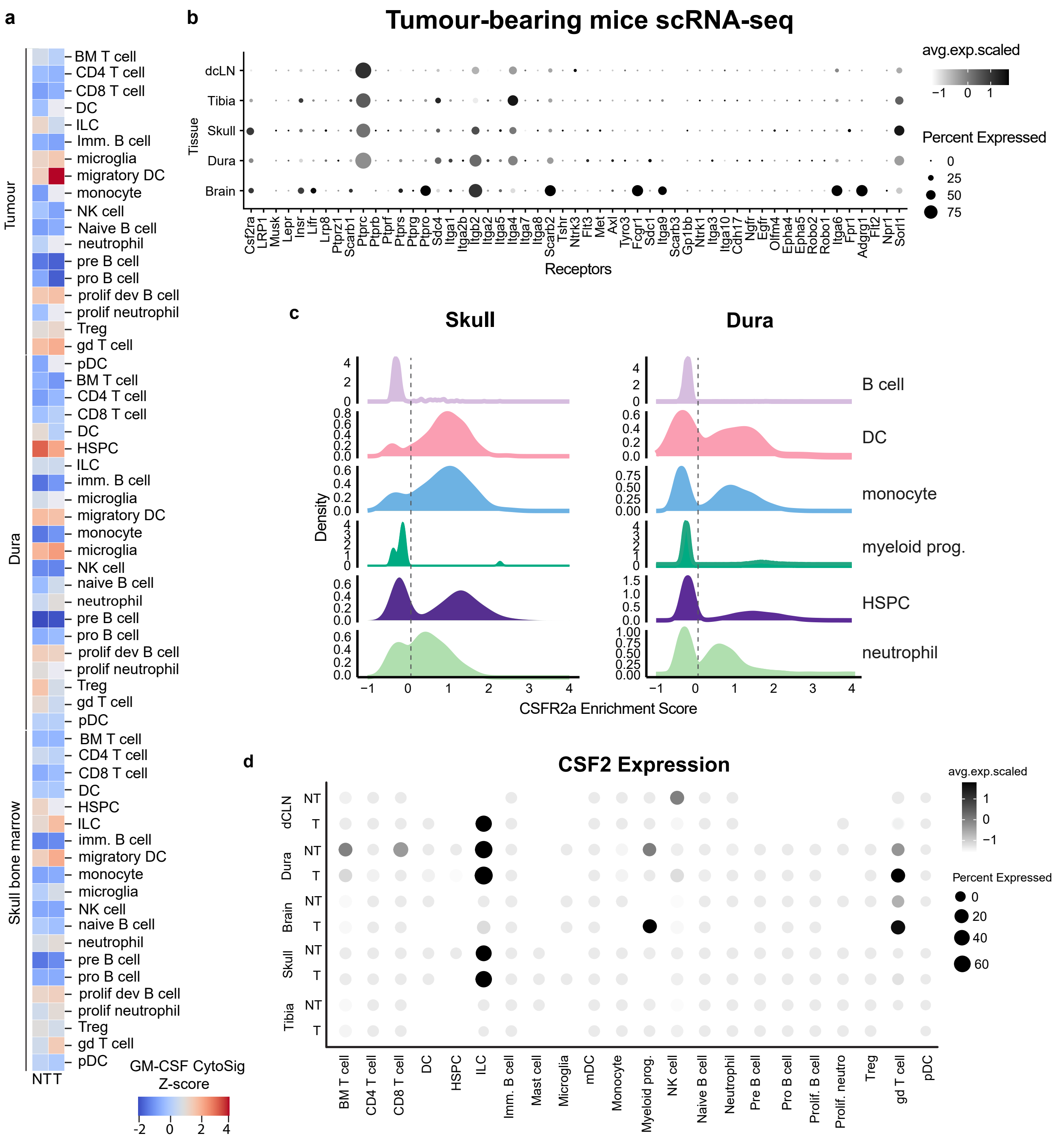

### SUPPLEMENTARY FIGURE 10

#### SUPPLEMENTARY FIGURE 10

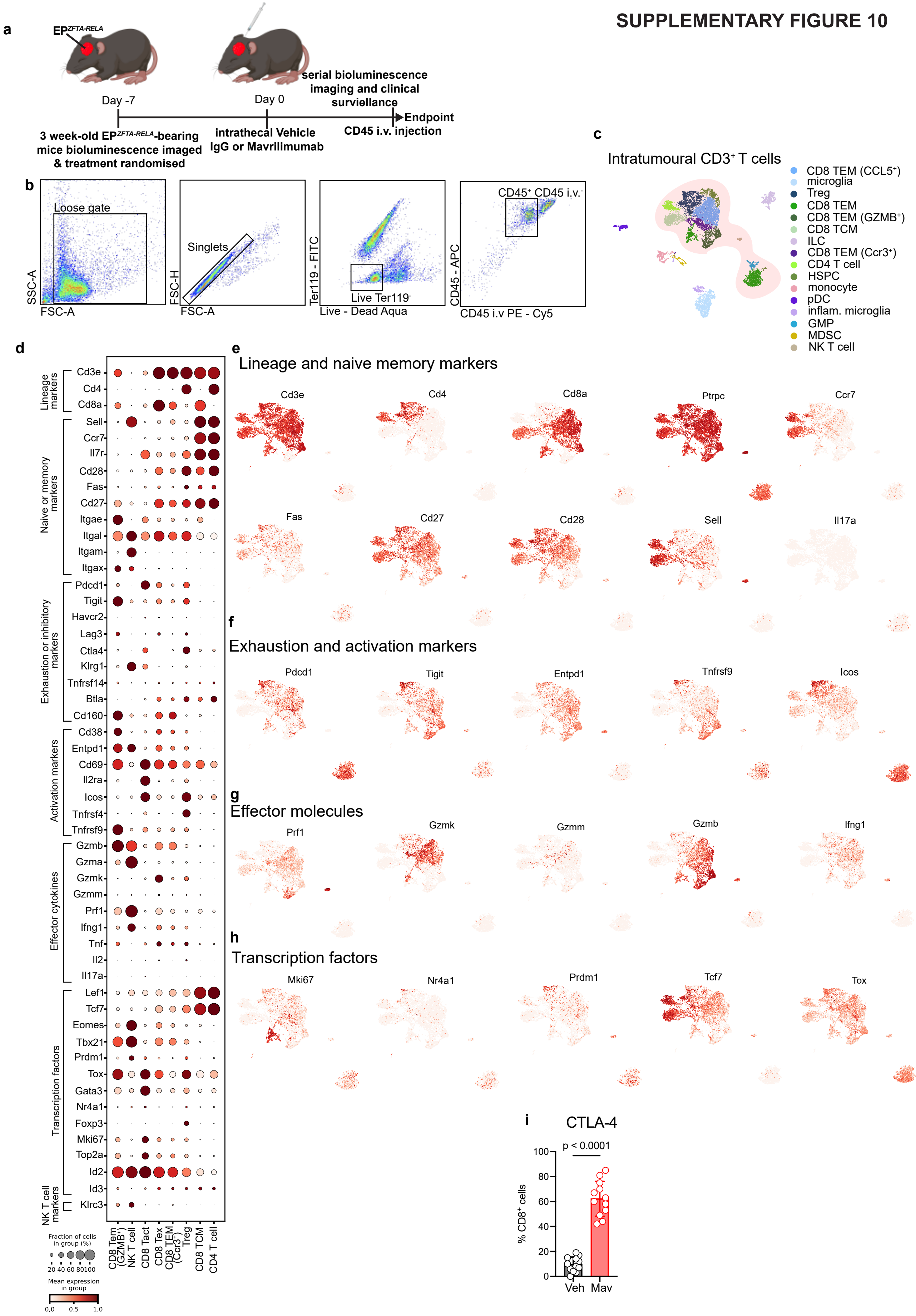
